# Estimating spatially adjusted temperature-dependent time-varying reproduction numbers for vector-borne diseases

**DOI:** 10.64898/2026.05.29.728636

**Authors:** Luis J Ponce, Esther Li Wen Choo, Diyar Mailepessov, Somya Bansal, Kelvin Bryan Tan, Kris V Parag, Jue Tao Lim

**Affiliations:** Lee Kong Chian School of Medicine, Nanyang Technological University, Singapore; Environmental Health Institute, National Environment Agency, Singapore; Saw Swee Hock School of Public Health, National University of Singapore, Singapore; Ministry of Health, Singapore; Department of Engineering, King’s College London, London, UK; MRC Centre for Global Infectious Disease Analysis, Imperial College London, London, UK

## Abstract

Estimating the effective reproduction number is crucial for understanding and managing infectious disease outbreaks. For vector-borne diseases like dengue, transmission depends on environmental and spatial conditions: temperature affects the extrinsic incubation period in mosquitoes, altering transmission timing, while spatial proximity can lead to clusters of transmission. We integrated a temperature-dependent (TD) generation time (GT) distribution and a spatial decay function weighting transmission likelihood by distance into the Wallinga & Teunis estimation framework. Simulations compared scenarios where the true generation time for spatially adjusted disease cases were TD or temperature independent (TI), versus their corresponding model assumptions. Daily reproduction numbers (*R*_*t*_) estimated were evaluated via percent error against true values. We found error to be predominantly driven by variance rather than bias, indicating that stochastic uncertainty in infector assignment was the primary driver of inaccuracies rather than systematic model miscalibration. Assuming a spatial weighting function assigning higher probabilities of transmission to geographically close infector-infectee pairs (Gaussian spatial decay) percent errors measuring deviation between estimated and true *R*_*t*_ outperformed those of alternative spatial kernels. Mis-specifying temperature-dependence under a Gaussian spatial kernel yielded higher errors when the ground truth was TD (16–38% vs 36–58% deviation), and similar errors when it was TI (32–52% vs 35–56% deviation), indicating limited sensitivity to temperature dependence when it was not present in the underlying transmission process. Application to dengue case data in Singapore showed that spatially adjusted models produced more variable *R*_*t*_ trajectories than time-series smoothing approaches, with Gaussian decay yielding more stable estimates than exponential decay and the TD model producing modest refinements, suggesting that explicitly modeling spatial heterogeneity and TD transmission dynamics increases responsiveness to even fluctuations in case counts. Our framework is robust across climates, shown by simulations using a broader temperature range, and suggests a useful application in retrospective analyses.

## Background

Accurate estimation of infectious disease transmissibility is particularly critical for vector-borne diseases (VBDs), where transmission is strongly shaped by vector ecology and environmental conditions. Interventions such as *Wolbachia* deployment, insecticide measures, or source-reduction policies aim to reduce transmission potential and have shown promising effectiveness in curbing dengue case counts.^1–4^ These more traditional approaches to assess intervention effectiveness against vector-borne infections often rely on incidence, attack rates, or prevalence, typically within study designs such as cluster-randomized controlled trials, pre-post intervention comparisons, or quasi-experimental methods like synthetic control.^1–3,5–7^ While such methods offer valuable insights such as quantifying reductions in case counts or the speed of epidemic growth, they offer limited information on whether declines in incidence can be attributed to imposed interventions. Further, approaches like randomized controlled trials require robust surveillance, making them difficult to implement, especially in resource-limited regions. Alternatively, to better understand whether interventions interrupt transmission, we require methods that reflect how many onwards infections each case is likely to generate, and how this varies over space and time.

The time-varying reproduction number, *R*_*t*_, provides a dynamic measure of the average number of secondary infections generated by an infected individual at a given time, *t. R*_*t*_ serves as a real-time indicator of epidemic trends and could inform intervention impact by assessing whether reductions in transmission potential are consistent with the timing of intervention implementation, making it a powerful tool for disease monitoring and response. Intervention effectiveness has been evaluated by using *R*_*t*_ across several diseases such as COVID-19 and dengue, demonstrating its potential to identify high-risk transmission areas and aid in outbreak management.^8–15^

Several methodological frameworks have been proposed to estimate *R*_*t*_, each offering distinct advantages and limitations. One foundational approach is the Wallinga-Teunis (WT) method, which uses the generation interval distribution, defined as the time between exposure of a primary case and their secondary case(s), to probabilistically infer who infected whom (infector-infectee pairs) from reported dates of symptom onset, leading to the estimation of individual- or cohort-level reproduction numbers, i.e., the estimation of the number of infections caused by each person/cohort.^16^ Bayesian approaches such as EpiEstim use the serial interval, defined as the time between symptom onset in a primary case and its secondary case, and case incidence data to estimate *R*_*t*_ in real time.^17^ Another approach, EpiFilter, employs a sequential Bayesian filter to iteratively refine estimates, providing an adaptive approach that accounts for observational uncertainty and addresses the censoring issues from the WT and EpiEstim methods.^18^ Although these methods differ in implementation, they all depend heavily on correct specification of the generation interval distribution. In practice, the generation interval is often approximated using the serial interval, which can introduce substantial bias when the mean, variance, or overall shape of the true generation interval are misspecified.^19,20^ This issue is particularly important for vector-borne diseases, where transmission involves both human and vector processes, resulting in more complex and temperature-dependent transmission dynamics that are poorly captured by simplified generation interval assumptions. Consequently, inaccurate specification of the generation interval can lead to biased *R*_*t*_ estimates and unreliable assessment of intervention effects.^20^

However, most existing reproduction number methods largely operate at the population level and rarely model the impact of space and temperature on vector-borne transmission. While some spatially adjusted methods have been developed to include geographical heterogeneity in vector-borne disease dynamics by estimating reproduction numbers for each individual case *l* (*R*_*l*_), allowing for refined assessments of local transmission potential, these do not account for a dynamic generation interval distribution.^21,22^ In this approach, the probability that some case *l* infected case *k* is determined by a temporal weighting function capturing the generation time distribution, and a spatial weighting function. This spatial function describes how the geographic distribution of past infections contributes to ongoing transmission. This is commonly assumed to decay monotonically with Euclidean distance to reflect decreasing transmission likelihood between geographically separated cases. Unlike *R*_*t*_ estimation methods that summarize transmission at the population level, these spatially adjusted approaches capture how transmission risk varies at the individual level, potentially revealing clusters of high transmission or localized outbreak sources that might not be captured in aggregate population-level estimates. This spatial approach is particularly relevant for VBDs, where mosquito dispersal is heterogenous across space, and other factors such as environmental (temperature and diurnal temperature range) and anthropogenic confounders (precipitation and vegetation density) may drive spatially clustered transmission patterns.^23–25^

Moreover, temperature is a well-established determinant of transmissibility for many vector-borne and endemic diseases, affecting pathogen replication, vector survival, and the duration of key transmission intervals.^9,26,27^ Temperature and diurnal temperature range (DTR) have been established as specific climatic factors affecting viral replication within *Aedes* mosquitoes, and can influence the duration of the extrinsic incubation periods (EIPs), and consequently, vector-borne diseases’ generation times.^28–30^ However, many existing models, including all the aforementioned ones, assume a static generation time distribution, which overlooks the temperature-dependent nature of disease transmission. Dengue, for example, exhibits substantial variation in generation time due to temperature-driven changes in the EIPs and mosquito biting rates.^9,31^ Warmer temperatures tend to decrease the EIP and are associated with a higher number of cases, and a wider DTR at low mean temperatures also produce a similar effect.^26,27,29^ Neglecting these effects can lead to biased estimates of *R*_*t*_, potentially misrepresenting transmission dynamics.

Motivated by these key limitations, we developed a framework which can explicitly model the dependence of generation time distributions on temperature and space to infer the effective reproduction number of vector-borne diseases by extending the framework of the WT method. We applied this framework to a series of simulations that incorporated realistic spatial and temporal transmission dynamics and exposed how misspecified or unmodelled transmission details limit our ability to infer and associate transmissibility changes with interventions. Using both simulated and empirical dengue data, we show that incorporating these dependencies yields more accurate and epidemiologically consistent estimates of *R*_*t*_, and better captures both local transmission heterogeneity and temperature-driven fluctuations in transmissibility.

## Methods

### Infection network representation

First, we followed the setup of Wallinga and Teunis (2004).^16^ VBD transmission can be modeled as a directed network, where nodes represent individual cases, and directed edges indicate successful infections. Each case has exactly one primary infector, except for imported cases, which have none. Given an outbreak with *n* cases, of which *q* are imported, the remaining *n* − *q* cases form the infection network. The total number of possible infection networks is determined by the number of ways these locally transmitted cases can be assigned a primary infector. We assumed that each case has exactly one primary case, and therefore each node in the infection network has exactly one incoming edge. Cases cannot infect themselves and no edges are directed to their own node.

We can uniquely represent our transmission network as a vector *v* of length *n* − *q*, of which each element *v*(*i*) represents the label of the primary case (infector) that has infected the case with label *i* (infectee). The directed edge from *ν*(*i*) to *i* defines who infected whom within the network. We set *ν*(*i*) = 0 for exogenous infections outside the population. The set of candidate infection networks which satisfy the above constraints is labelled *V*, where card(*V*) = (*n* − *q*)^*n*−1^ represents its cardinality (i.e., the total number of possible infection networks). For each of the *n* − *q* non-imported cases, there are *n* − 1 potential infectors.

### Likelihood-based inference of the infection network

We inferred the most likely infection network using a likelihood-based model that assumes transmission occurred only among reported cases. We considered a probability model to infer the likelihood that a candidate infection network *v* ∈ *V*, which specifies who infected whom, underlies the observed epidemic curve, *ξ*. This curve is the aggregate time series of infections and in practice (due to surveillance limitations) does not distinguish between locally generated and imported infections. A key component of this model is the probability density function of the spatiotemporal generation time, which describes how transmission likelihood varies jointly with the elapsed time of infection *t* (approximated by symptom onset date) from case *j* to case *i*, and the spatial distance *d* between the two cases, *w*(*t*_*i*_ − *t*_*j*_, *d*_*i*_ − *d*_*j*_|*θ*), where *w*(0, *d*_*i*_ − *d*_*j*_ | *θ*) = 0 (i.e., transmission is never instantaneous). The arguments {*t*_*i*_ − *t*_*j*_, *d*_*i*_ − *d*_*j*_} represent the interval of time and space between cases *i, j*, while θ is the vector of parameters that specify the probability distribution. In the absence of ξ, each infection network is considered equally likely, assuming that there is independence between unobserved transmission events from case *j* to case *i* and from case *j* to any other case *k*. This is referred to as the “independence condition.”

We expand the WT approach to consider the candidate epidemic curve *ξ*, with parameters *θ*, for a space- and time-dependent generation interval. Let *v* represent a candidate infection network, with *t*_*i*_ and *d*_*i*_ denoting the time of infection and location of individual *i*, respectively.^16^ Then the likelihood of that infection network is given by:

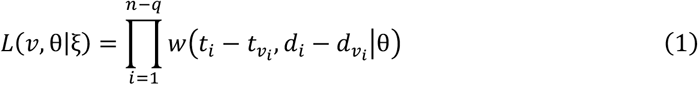

Let 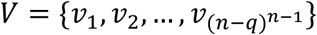 be the set of candidate infection networks among the *n* − *q* locally transmitted cases, and let *c*(*v*|θ) represent the weight function for each infection network. Each of these *n* − *q* cases can potentially have any of the remaining *n* − 1 cases as their primary infector (i.e., no self-infection), and every distinct combination of such assignments corresponds to one possible infection network in *V*. Then for the likelihood of all possible infection networks with *n* − *q* locally infected nodes, *c*(*v*|θ) is implied to be a constant by the independence condition, denoted as *c*. Under this assumption, the likelihood of the full transmission network can be factorized into the product of the likelihood contributions from each individual case. Consequently, rather than summing over all possible transmission networks jointly, the summation can be separated into independent sums over all possible infectors for each case *i*:

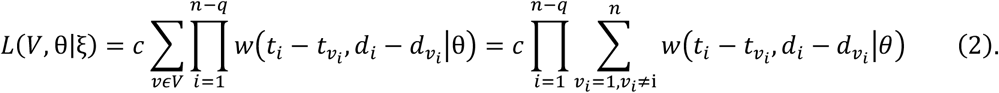

In the case of individual *l* infecting individual *k* over the set of infection networks *V*,

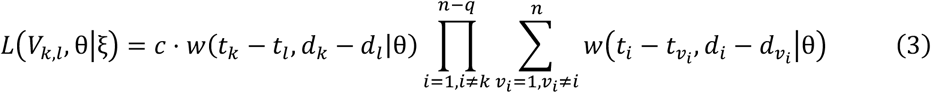

The relative likelihood that *l* infects *k* is:

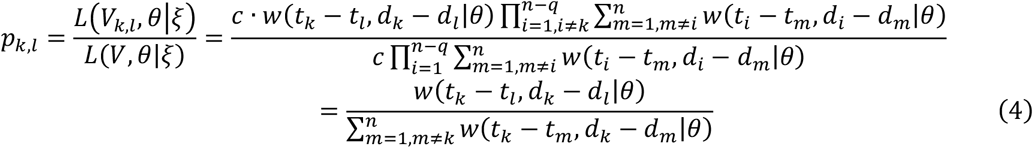

We assumed that the spatiotemporal generation time is separable into space and time components, that the time component is temperature (*T*) dependent (i.e., a vector-borne disease where the EIP depends on temperature at the time of human symptom onset), and that geographical variations in temperature across the small city-state of Singapore are negligible relative to variations over time. In addition, due to Singapore’s low daily variations in temperature and consistently high mean temperatures, we assumed a negligible effect on the EIP due to DTR.^29^ Other climatic factors such as humidity were considered, but not incorporated in this study as they have not shown to have any effect on viral replication during the EIP.^30^ The formulation is generalizable and can accommodate case-specific temperatures, *T*_*k*_, without changes to the underlying likelihood structure, such that the spatiotemporal weight of transmission from individual *l* to individual *k* is:

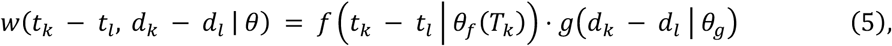

where *θ*_*f*_ (*T*_*k*_) denotes the temperature-dependent parameters of the generation time distribution for case *k*, and *θ*_*g*_ denotes the set of parameters describing the spatial transmission function. We can then write Eq (4) as:

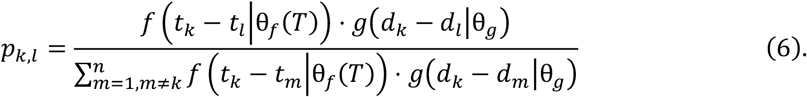

Then the distribution of the individual reproduction number for individual *l* is given by

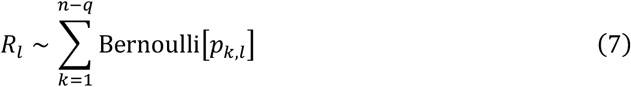

with expectation

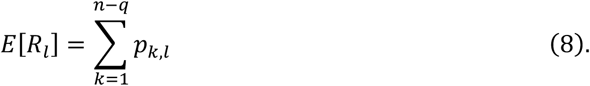

We derived confidence intervals (CIs) for the estimated reproduction numbers using a bootstrap approach. To quantify uncertainty, we resampled individuals with replacement from the simulated epidemic data. For each bootstrap sample, *R*_*l*_ was recalculated for all cases in the sampled network, and the time-varying reproduction number *R*_*t*_ for each day was computed as the arithmetic mean of *R*_*l*_ values corresponding to cases with onset on that day within the same bootstrap sample, 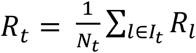, where *N*_*t*_ is the number of cases with onset on day *t*, and *I*_*t*_ denotes the indices of cases with onset on day *t*. Thus, *R*_*t*_ represents the average number of secondary infections per case per day, calculated separately for 100 bootstrap samples. The 2.5 and 97.5 percentiles of these bootstrap distributions were then used as the bounds of the 95% CIs, providing inference that accounts for stochasticity in individual transmission and temporal variability in the generation time.

### Temperature-dependent generation times

The generation time distribution was modeled as a function of temperature due to its strong influence on the EIP in mosquitoes. Because transmission occurs through a vector, the generation time consists of four sequential components: the intrinsic incubation period (IIP) in humans, the EIP in mosquitoes, the time for a human to infect a mosquito, and the time for an infectious mosquito to transmit the virus to a human. Each of these components follows a distinct probability distribution. Adapted from Codeço et al. (2018) and Chan and Johansson (2012), the IIP was modeled as a gamma-distributed interval, capturing the variability in the time it takes for an infected human to become infectious, with shape *α*_*IIP*_ = 16 and rate *λ*_*IIP*_ = 2.7.^9,28^ Its mean *μ*_*IIP*_ is given by:

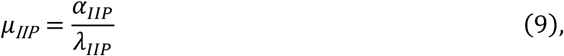

Following the work of Chan and Johansson (2012), the EIP, which describes the time required for viral replication within the mosquito was also modeled as a gamma distribution, but with shape *α*_*EIP*_ = 4.3 and now the rate parameter *λ*_*EIP*_(*T*) = (*β*_0_ − *β*_1_*T*)^−1^ (where *β*_0_ = 7.9, *β*_1_ = −0.21) is temperature-dependent, allowing for dynamic adjustments based on environmental conditions, with mean *μ*_*EIP*_

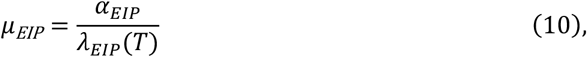

where β_0_ is the baseline scaling parameter, and β_1_ quantifies the sensitivity of the rate parameter to temperature *T*.

The remaining two components, human-to-mosquito transmission (*t*_H → M_) and mosquito-to-human transmission (*t*_M → H_), were both assumed to follow exponential distributions, as computed by Codeço et al. (2018)^9^:

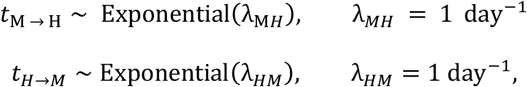

reflecting the memoryless nature of mosquito biting behavior.

Since the total generation time is the sum of these independent time components, we derived its overall distribution using moment-generating functions.^32^ This resulted in a compound gamma-exponential distribution, with the temperature-dependent rate parameter of the EIP influencing the overall shape and scale of the generation time distribution. By incorporating daily temperature data into this framework, we allowed the generation time distribution to vary dynamically throughout the epidemic, providing a more accurate representation of transmission dynamics under fluctuating environmental conditions. The relationship between temperature and generation time is visualized in **Figure 1A**. When temperature dependence is ignored, the temporal kernel *f* (*t*_*k*_ − *t*_*l*_ | *θ*_*f*_(*T*)) is replaced by a fixed *f*(*t*_*k*_ − *t*_*l*_ | *θ*_*f*_), such that a single, constant generation time distribution is applied across all temperature values instead of one that varies dynamically with temperature.

**Figure 1.**
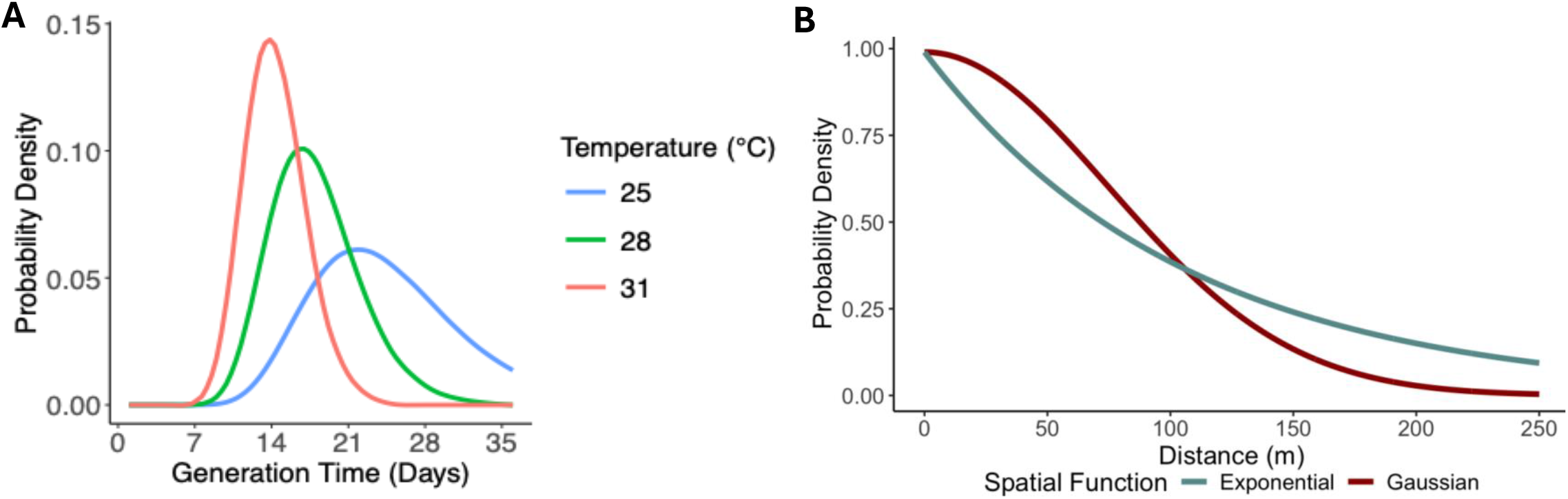
Generation time distributions across different temperature values (A) and exponential and Gaussian decay spatial functions dependent on distance.

To incorporate spatial dependencies in the estimation of *R*_*l*_, spatial weights were calculated using the Euclidean distance between cases based on their geographical coordinates. These spatial weights were derived using both a Gaussian and an exponential decay function to reflect the influence of proximity and the limited flight range of *Ae. aegypti/albopictus*. The exponential decay function decays faster than the Gaussian for transmission pairs closer in distance, but decays more slowly as distance between them increases (**Figure 1B**).

Let *d*_*k,l*_ represent the distance between successive cases *k* and *l*. The spatial weight for each pair was calculated as either a Gaussian decay function:

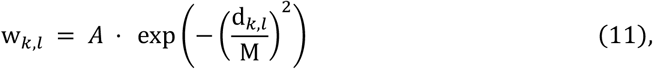

or as an exponential decay function:

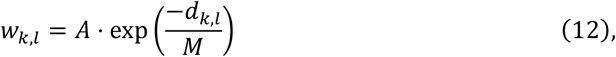

where *A* is the amplitude representing the maximum weight a distance approaches zero, and *M* is a parameter adjusting the decay rate, corresponding to the assumed maximum flight range of *Ae. aegypti* (106 meters).^33^ When we ignore spatial effects on transmissibility, we assume that the likelihood of transmission is uniform with respect to space, such that *g*(·) in Eq (5) is set to a constant value of 1.

This formulation ensures that weights decrease rapidly as the distance between cases increases, reflecting the localized nature of mosquito-borne transmission. Closer cases exert greater influence, while the influence of cases beyond *M* is small. The resulting spatiotemporal weight matrix *w*(*t*_2_ − *t*_1_, *d*_2_ − *d*_1_ | *θ*) quantifies the spatial contribution of each case to others and is incorporated into the spatiotemporal *R*_*l*_ model to account for spatial structure in the data. In the case of *Aedes* mosquitoes and their limited flight range, the spatial component of transmission likelihood approaches zero even between individuals separated by only a few hundred meters, highlighting the highly localized nature of transmission.

### Simulation-based evaluation of effective reproduction numbers

To assess the accuracy of the proposed estimation method, we simulated dengue epidemics in Singapore and compared our estimated *R*_*l*_ with their true values from each simulation. We assumed true *R*_*l*_ to be derived under the model described in Eq (7), where the probability of infection between individuals depended on both temperature and spatial distance. The temporal component of the transmission likelihood varied daily with temperature through the EIP, and the spatial component of the likelihood decayed with distance. Both an exponential and Gaussian decay kernel were tested as being the ground truth. Detailed parameter values are provided in the Supplementary Materials. The accuracy of these estimates was evaluated using the mean absolute error (MAE) and percent error as measures of deviation from true *R*_*l*_ values. In addition, we computed the area under the receiver operating characteristic curve (AUC-ROC) to assess how well the method classified cases based on their transmission potential. For this analysis, we categorized *R*_*l*_ into less than 1 or greater than 1, representing whether an individual sustains the transmission chain or not, respectively.

The simulation setup involved 10,000 individuals over a period of 365 days or until all individuals became infected. Each individual was assigned random geographical coordinates within Singapore (longitude 103.60º to 104.03º and latitude 1.23º to 1.47º). The epidemic began with the random selection of a single index case, whose infection time was set to *t* = 1. To match the simulated epidemics to real data, *t* = 1 was set as 1 January 2019, and daily average temperature data in Singapore (2019) from Meteorological Service Singapore were subsequently used to compute Eq (6).^34^ At each subsequent time step *t*, infectious individuals attempted to infect susceptible individuals based on the probability *p*_*k,l*_ defined in Eq (6). The infection process was modeled as a Bernoulli draw for each susceptible individual *k*. The resulting “true” individual reproduction number, *R*_*obs,l*_, representing the total number of successful infections for the infector *l*, follows the distribution described in Eq (7) with the estimated reproduction number, *R*_*estim,l*_, given by Eq (8).

The mean absolute error (MAE) between the true and expected reproduction numbers was calculated as:

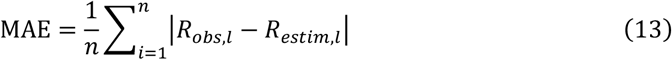

where *n* is the total number of individuals. The mean squared error (MSE) between true and expected reproduction numbers was also calculated, using the following equation:

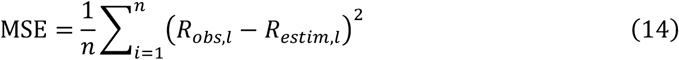

Across 100 simulation runs, the mean MAE and MSE were computed to summarize the performance of the *R*_*l*_ estimation method. Percent errors were similarly calculated to measure relative deviation, defined as (*MAE*/*mean*(*R*_*obs*_)) × 100.

To evaluate the population-level trend in disease transmissibility, individual-level *R*_*l*_ estimates were aggregated by day of infection to calculate the time-varying reproduction number *R*_*t*_. Specifically, *R*_*t*_ was computed as the arithmetic mean of all *R*_*l*_ values corresponding to cases infected on the same day and similarly assessed for its deviation from true *R*_*t*_.

To identify sources of error, we further decomposed MSEs into the variance and squared bias of the estimation error,^35^ *R*_*estim*_ − *R*_*obs*_, as follows:

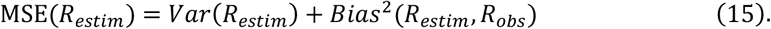

*R*_*k,l*_ was defined as the linear sum of independent random variables *Z*_*k,l*_ where each *Z*_*k,l*_ represents the transmission contribution between cases determined by exogenous spatial and temporal kernel weights. Under the assumption that all *Z*_*k,l*_ are independent and identically distributed, the expectation *E*[*R*_*k,l*_] corresponds to the mean estimate of *R*_*k,l*_. The variance of *R*_*k,l*_ can therefore be expressed as the variance of a linear sum of independent random variables, which expands to the sum of individual variances plus covariance terms. However, because *Z*_*k,l*_ are independent, all covariance terms are zero, making the variance of *R*_*k,l*_ equal to the sum of variances of the individual *Z*_*k,l*_ terms. In the implementation, *R*_*l*_ corresponds to the aggregation of *R*_*k,l*_ over all infectious pairs within each simulation and therefore inherits this variance structure at the simulation level. At the daily temporal level, bias was defined as the mean deviation between estimated and observed *R*_*t*_: *E*[*R*_*estim*_ − *R*_*obs*_]. Variance, on the other hand, was computed as the empirical variance of the estimation error across infection times within each simulation. The decomposition was verified by comparing Eq (15) with the direct empirical MSE = *E*[(*R*_*estim*_ − *R*_*obs*_)^2^], which were numerically equivalent.

### Robustness checks

We first examined how alternative parameterizations of the generation time distribution affect estimated reproduction numbers. The baseline model incorporated a temperature-dependent generation time, where the EIP followed a gamma distribution with a rate parameter that varied with temperature. To test this assumption, we also considered static generation time distributions, *f*_*mis*_(*t*|*μ, σ*), that have been used in previous literature.^21,31^ Specifically, we evaluated two different gamma-distributed intervals with fixed parameters with means of 20 and 18.2 days (μ = 20, 18.2), and with standard deviations of 9 and 6.1 days (σ = 9, 6.1), altering Eq (6) to become:

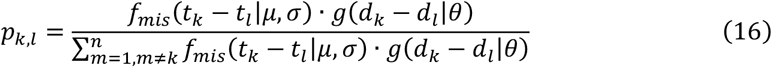

where the misspecified temporal weight is given by the gamma distribution

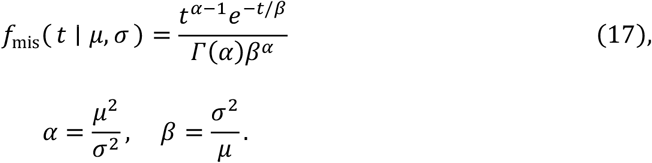

Our temperature-dependent generation time distributions varied for each 0.1 difference in temperature between 20–36.8ºC. Therefore, the means of each distribution ranged from 9.8 to 31.4 days, with higher temperatures having shorter mean generation times. Similarly, the standard deviations tended to be shorter for higher temperatures, ranging from 2.1 to 7.5 days.

We next examined how omission of spatial structure affects *R*_*l*_ estimation. In this scenario, we assumed a generation interval that depended only on time, with no spatial component. By removing the spatial term *g*(*d*_1_ − *d*_2_ | θ_*g*_) from the model, the calculation of *p*_*k,l*_ simplified to:

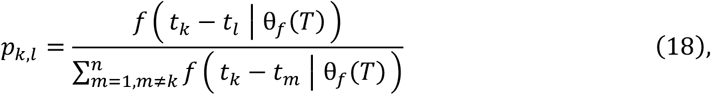

where *θ*(*T*) is the set of parameters that specify the temperature-dependent probability distribution.

Furthermore, we tested various cutoff values for the maximum possible distance between two successive infected cases and for the maximum possible generation times. Our baseline model assumed a maximum distance of 1.5km and a maximum generation time of 5 weeks, based on previous literature assessing maximum flight ranges of *Ae. aegypti* and maximum generation times of dengue.^36–38^ Our sensitivity analyses included simulations of all possible combinations of temperature-dependent and independent generation times, distance cutoffs of 500m, 1km, and 1.5km, and generation time cutoffs of 4, 5, and 6 weeks, for both Gaussian decay and exponential decay distance kernels. Likewise, we considered and simulated scenarios without distance dependence by only varying the generation time cutoffs for both temperature-dependent and independent generation times. 100 simulations were performed for each possible combination of parameters. Additionally, we tested varying amplitudes through sensitivity analyses for *A* = 0.8, 0.5, 0.3 in Eq (11) to assess the effect of misspecifying the spatial decay weighting function.

Finally, the performance of the model under a wider temperature than that observed in Singapore was assessed. Historical daily temperature data from Kaohsiung, Taiwan in 2019 were used, as Kaohsiung also observes a high number of dengue cases, but has a wider range of temperature compared to Singapore.^39^

### Application to the 2019 dengue outbreak in Singapore

To examine the performance of our model in a real-world scenario, we applied the present spatiotemporal temperature-dependent *R*_*t*_ estimation procedure Eq (6) to empirical dengue case data in Singapore from 1 January 2019 to 28 December 2019. Singapore is a multi-ethnic Asian city-state where dengue is endemic, with at least three serotypes documented to be in constant circulation.^40,41^ Dengue is legally notifiable to the Ministry of Health within 24 hours from laboratory confirmation with confirmatory diagnostic testing for dengue via NS1 antigen assays/IgM enzyme-linked immunoassay is widely available across healthcare settings.^40,41^ 2019 data was utilized to match the daily temperature data used in the simulations above.

Models were applied 100 times for both spatial decay weighting functions Eq (11) and Eq (12), and without a spatial component Eq (18) to the same dataset. The same set of models were applied using a temperature-independent generation time distribution Eq (16). Individual reproduction numbers and daily time-varying reproduction numbers, *R*_*l*_ and *R*_*t*_, were both estimated, along with their 95% CIs calculated through bootstrapping. Model specificity was assessed by comparing predicted absence of transmission (*R*_*t*_ < 1) against observed absence of cases, calculated as the proportion of correctly predicted no-transmission days.

We also evaluated the time-varying reproduction numbers for the same data but with EpiFilter, which assumes that the observation model is a Poisson renewal equation and treats *R*_*t*_ as a latent state that evolves smoothly over time. The grid on the reproduction numbers was set to have a minimum (Rmin) of 0.01, a maximum (Rmax) of 4, and a size (m) of 5000. The diffusion noise (*η*) was set to 0.25, and we assumed that *R*_*t*_ was gamma-distributed with shape=5 and scale=0.5. For the total infectiousness, we used a gamma-distributed, temperature-independent generation time with mean = 18.2 and SD = 6.1 days.

## Results

### Accuracy of individual reproduction number estimates

We first evaluated the accuracy of *R*_*l*_ estimates using MAEs across models with different spatial transmission kernels and generation time assumptions. Among models with a Gaussian decay spatial kernel, MAE values ranged from 1.10–1.17, and AUC-ROC scores from 0.58–0.61, when applying a temperature-dependent estimation method to epidemics simulated with temperature-dependent generation times. In contrast, models with an exponential decay kernel yielded MAE values of 1.35–1.43 and AUC-ROC scores of 0.59–0.65. Detailed values for each simulation scenario are listed in **Table S2**.

### Aggregated reproduction number estimation

Daily reproduction numbers, *R*_*t*_, were computed as the arithmetic mean of *R*_*l*_ values for all cases infected on the same day. Estimation accuracy was assessed via percent errors relative to each simulation’s true daily *R*_*t*_, and models were also evaluated on their ability to correctly classify *R*_*t*_ as below or above 1 using AUC-ROC scores. **Table 1** summarizes results under matched spatial kernel assumptions between the simulated epidemics and estimation models, where both used either exponential or Gaussian decay kernels. When both the simulations and estimation models assumed temperature-dependent generation times, percent errors were 16–38% under a Gaussian kernel and 31–363% under an exponential kernel. When the ground-truth simulations instead used temperature-independent generation times, with the estimation model also assuming temperature-independent generation times, percent errors in were 32–52% under a Gaussian spatial kernel and 26–49% under an exponential spatial kernel. When both the simulated true generation times and the estimation model assumed temperature-dependent generation times, AUC-ROC values ranged from 0.77–0.96 for Gaussian decay kernels and 0.89–1.00 for exponential decay kernels. In contrast, when both the simulations and estimation assumed temperature-independent generation times, AUC-ROC values ranged from 0.79–0.98 and 0.82–1.00 for Gaussian and exponential kernels, respectively. Full results are shown in **Table 1** and **Figure S4** and **S5**.

**Table 1.** Mean percent errors were calculated according to Eq (13) divided by the average true R_t_ multiplied times 100 (MAE/mean(R_t_obs_)*100). The range depicts the minimum and maximum values obtained from each of the 100 simulations. Area Under the Receiver Operating Characteristic curve (AUC-ROC) was calculated by comparing binary observed transmission status (*R*_*t*_ ≥ 1 vs < 1) against continuous mean *R*_*t*_ values. The ROC curve was generated by varying the threshold on predicted *R*_*t*_, and the area under this curve was used as the measure of discriminatory ability. Similarly, the AUC-ROC ranges were obtained from the minimum and maximum values yielded by each of the sets of 100 simulations. TD: temperature dependent. TI: temperature independent.

| Spatial kernel | Generation times in simulations | Generation times in estimation | Mean percent errors (range, %) | Mean MSE (range) | Mean bias<sup>2</sup> (range) | Mean variance (range) | AUC-ROC (range) |
| --- | --- | --- | --- | --- | --- | --- | --- |
| Exponential decay | TD | TD | 206 (40–373) | 15.48 (10.55–25.22) | <0.01 (<0.01–<0.01) | 15.48 (10.55–25.22) | 0.97 (0.89–1) |
|  | TI | TI | 36 (26–49) | 2.31 (1.76–3.08) | <0.01 (<0.01–<0.01) | 2.31 (1.76–3.08) | 0.94 (0.82–1) |
|  | TD | TI | 171 (103–273) | 4.29 (2.19–6.93) | 0.74 (0.61–0.93) | 3.55 (1.26–6.31) | 0.97 (0.90–1) |
|  | TI | TD | 55 (36–77) | 4.30 (2.72–7.62) | <0.01 (<0.01–<0.01) | 4.30 (2.72–7.62) | 0.94 (0.84–1) |
| Gaussian decay | TD | TD | 27 (16–38) | 2.81 (2.65–3.02) | <0.01 (<0.01–0.02) | 2.81 (2.64–3.02) | 0.88 (0.77–0.96) |
|  | TI | TI | 43 (32–52) | 1.81 (1.71–1.89) | 0.09 (<0.01–0.26) | 1.72 (1.56–1.87) | 0.89 (0.79–0.98) |
|  | TD | TI | 39 (36–58) | 2.49 (2.36–2.65) | <0.01 (<0.01–0.02) | 2.49 (2.34–2.65) | 0.88 (0.76–0.95) |
|  | TI | TD | 45 (35–56) | 2.16 (2.01–2.31) | 0.07 (<0.01–0.32) | 2.08 (1.79–2.30) | 0.90 (0.80–0.98) |

### Sensitivity to model assumptions

We computed the MAE and percent error of estimated daily *R*_*t*_ across combinations of distance cutoffs (500m, 1km, 1.5km), generation time thresholds (4, 5, and 6 weeks), and generation time assumptions (temperature-dependent [GT1], temperature-independent with mean = 20 days and standard deviation = 9 days [GT2], and with mean = 18.2 days and standard deviation = 6.1 days [GT3]). For models using a Gaussian spatial decay weighting function, mean MAE values ranged from 0.25 to 0.32, and mean percent errors from 21.5% to 27.9% (**Figure 2A**) across all combinations. For models using an exponential decay spatial weighting function, mean MAE values ranged from 0.39 to 3.00, and mean percent errors from 27.5% to 204.6% (**Figure 2B**). Non-spatial models resulted in mean MAE values ranging from 1.72 to 19.7, and mean percent errors ranging from 23.7% to 277% (**Figure 2C**).

**Figure 2.**
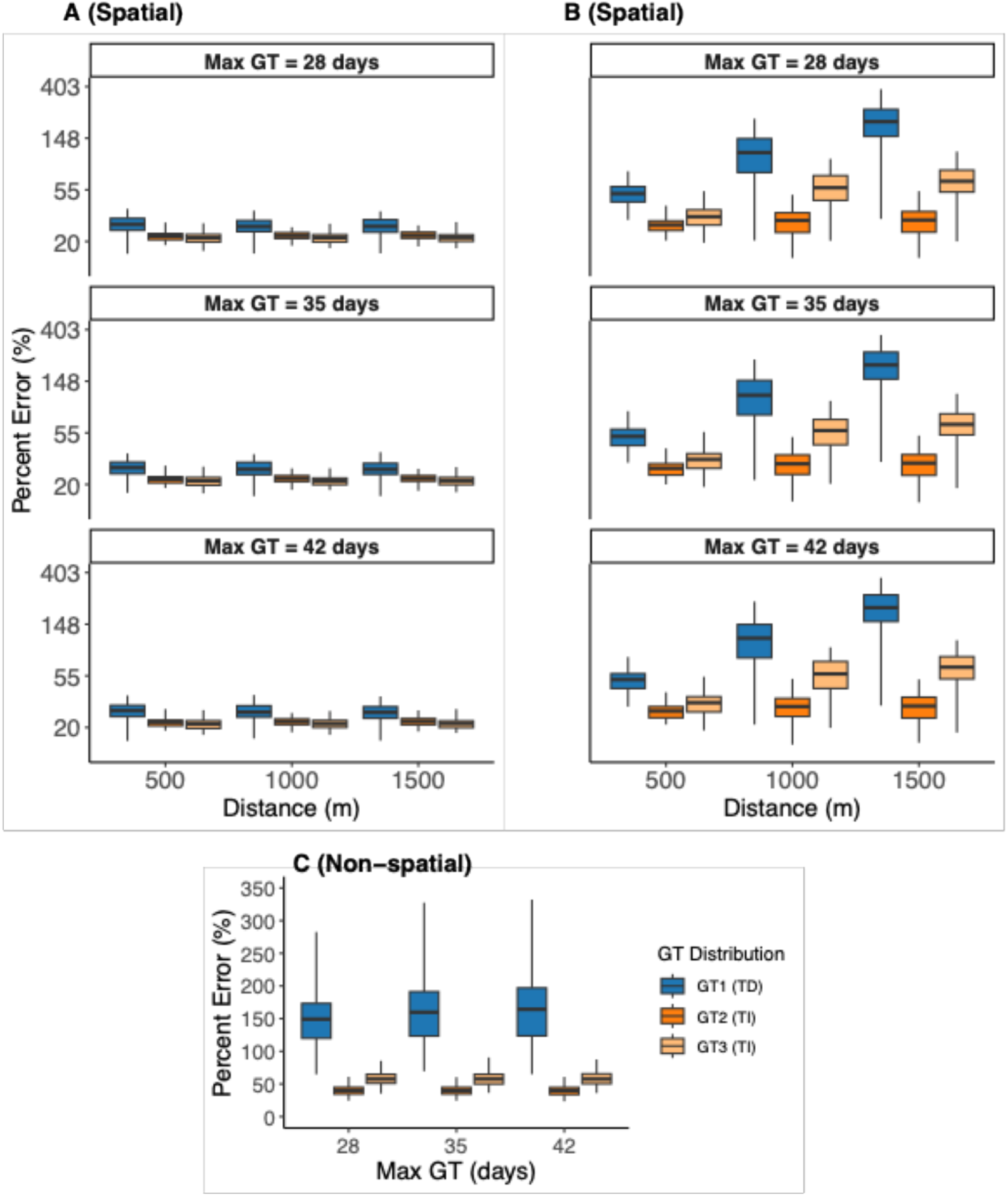
Errors in *R*_*t*_ estimation compared to true values. GT: Generation time. **A** and **B** depict the percent errors for simulations involving Gaussian and exponential decay spatial weighting functions, respectively, stratified by different maximum distances between successive cases (x-axis) and maximum GTs. To standardize the y-axis scale, percent errors for log transformed. **C** depicts the model results without spatial heterogeneity, separated by maximum GT on the x-axis. GT1 (TD): temperature-dependent GT distribution. GT2 (TI) and GT3 (TI): temperature-independent GT distribution with mean = 20, SD = 9, and mean = 18.2, SD = 6.1 days, respectively.

Using Eq (12) as the spatial weighting function and under non-spatial models, temperature-independent generation time distributions tended to result in lower errors compared to their temperature-dependent counterparts. However, for models using Eq (11) as the spatial weighting function, temperature-dependent GT distributions consistently yielded lower errors than the temperature-independent ones. Apart from temperature-dependent exponential decay spatially weighted models, the non-spatial models resulted in larger errors than all the rest of the models. Differences between maximum generation times and distance cutoffs produced minimal variations in the resulting errors. Full MAE and percent error statistics for each configuration are presented in **Figure 2** and **Table S2–S6**.

To examine whether the difference in estimation accuracy between models arose primarily from differences in the mean or the shape of the temperature-dependent generation time distribution, we compared results from temperature-dependent simulations under conditions where the mean generation time matched that of the temperature-independent model (26.9ºC). The mean error in R_t_ estimates on days where the mean generation time was equal to that of the temperature-independent mean (18.2 days) was 16.5% (ranging from 2.7–49.5%) within the simulations that assumed a temperature-dependent generation time distribution. Under the same conditions, the temperature-independent model (GT3) resulted in a mean percent error of 21.6% (17.3–28.1%) across all simulations (**Table S4**).

In addition to misspecified generation times, we performed sensitivity analyses to examine the effect of misspecified spatial weight functions by altering *A* in Eq (11), which represents the maximum transmission weight for cases at nearly zero distance from each other. Varying *A* tests the sensitivity of *R*_*t*_ estimates to assumptions about how strongly nearby cases contribute to transmission. For *A* = 0.8, 0.5, 0.3, the respective mean percent errors for *R*_*t*_ were 32.63, 32.96, and 33.24 (ranges 20.09–50.33, 18.64–37.25, and 20.93–38.63%), respectively. Box plots with these errors are depicted on **Figure S1A**.

Similarly, the effects of a wider temperature range were examined by utilizing mean daily temperature data from Kaohsiung, Taiwan in 2019, which ranged from 16.8–30.9ºC, whereas Singapore’s temperature ranged from 24.9–30.1ºC. The mean percent error from the 100 simulations under the wider temperature range was 33.29% (range 19.71–50.15%). Summary results are illustrated on **Figure S1B**.

### Real-world model performance

We analyzed a total of 3490 dengue cases in Singapore, reported from 1 January to 28 December 2019, with incidence fluctuating across the year, with a large observable peak in July and a smaller one in May (**Figure 3A**). Estimates of *R*_*t*_ from EpiFilter captured a couple of the incidence peaks, during which it accurately evaluated *R*_*t*_ to be above 1, and consistently remained usually between 1 and 3, with occasional troughs dropping below 1, and associated 95% CIs (**Figure 3B**). To improve interpretability and facilitate comparison of the models developed in this study with EpiFilter, 7-day rolling averages of *R*_*t*_ are shown for panels C–E in Figure 3. Spatially explicit models using exponential and Gaussian decay functions produced more variable and generally lower *R*_*t*_ trajectories compared to EpiFilter, with estimates occasionally dropping to zero (**Figure 3C–D**). The non-spatial model produced a similar pattern (**Figure 3E**). Across **Figure 3C–E**, both temperature-dependent and temperature-independent formulations were evaluated, with the temperature-dependent version shown in red and the temperature-independent version in blue. Specificity, defined as the proportion of correctly predicted no-transmission days, was similarly high across all models: 0.985 for the exponential decay model, 0.977 for the Gaussian decay model, and 0.988 for the non-spatial model.

**Figure 3.**
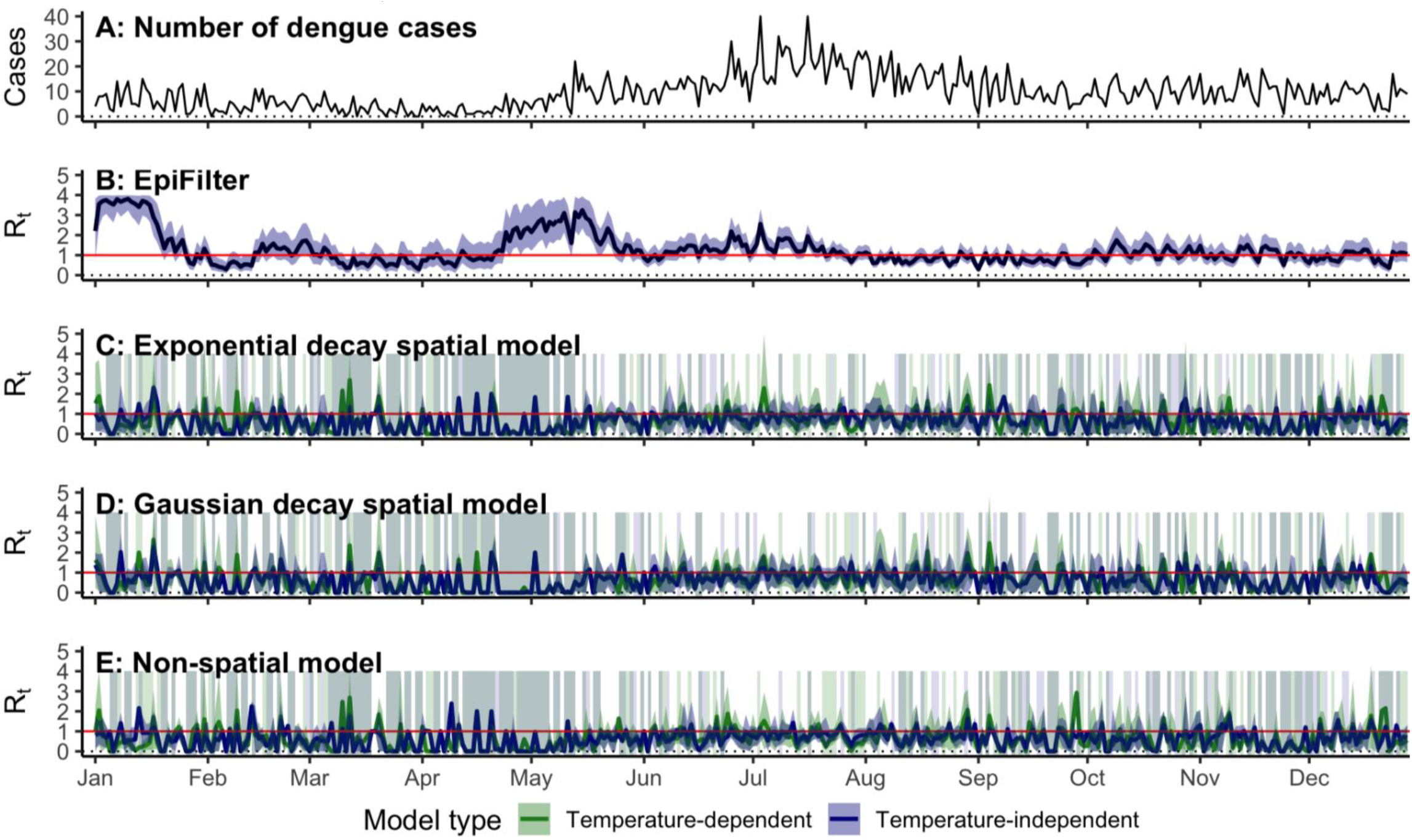
Singapore 2019 dengue *R*_*t*_ estimates across different models over time. Daily reported dengue cases in Singapore (2019) are depicted in **A**. Daily *R*_*t*_ estimates using EpiFilter **B** and the different models developed in this study (exponential and Gaussian decay spatial models, and non-spatial model, **C–E**) are plotted with their 95% CIs for both temperature-dependent (green) and -independent (blue) generation times. Shaded regions indicate timepoints where *R*_*t*_ estimates and their 95% CIs do not include 1 in the respective temperature-dependence color. *R*_*t*_ = 1 is marked in red.

## Discussion

This study demonstrates that the benefits of incorporating temperature-dependent generation times in *R*_*t*_ estimation depend partly on appropriate specification of spatial transmission dynamics, as adding temperature dependence alongside misspecified spatial kernels can reduce accuracy relative to simpler temperature-independent, spatially unstructured models for infectious diseases with strong spatial and environmental heterogeneity. We further applied the same temperature-dependent, spatially adjusted models across settings with both narrow (Singapore) and wider (Taiwan) temperature ranges and observed relatively small differences in predictive accuracy between them. This suggests that the framework is broadly transferable across different temperature regimes, although the magnitude of improvement from temperature dependence may vary depending on the underlying temperature variability and its representation in the model (**Figure S1**). This may partly reflect greater within-day temperature variability in Taiwan that is not captured when using daily mean temperatures, suggesting the need to also incorporate diurnal temperature ranges into the estimation model for regions with higher temperature variability. This may also reflect uncertainty in the timing of the EIP, for which the temperature on the day of the infectee’s onset is used as a proxy, because such approximations are unlikely to strongly affect estimates in a narrow temperature range such as Singapore’s but may affect the apparent benefit of temperature-dependent modeling in settings with wider temperature variation. We examined the performance of the temperature-dependent formulation when the mean of the generation time distribution was set equal to that of the temperature-independent generation time distribution. Across simulations, percent errors were similar to those of the temperature-independent model (16.5% vs 21.6%), suggesting that, in this setting, variation in the EIP shape may have had limited additional impact on *R*_*t*_ estimation accuracy.

Furthermore, we found that models that did not incorporate spatial heterogeneity within the likelihood of transmission and similarly assumed a maximum generation time of 35 days and temperature-dependent EIPs yielded errors of up to 277% (range: 74–277%) when estimating *R*_*t*_ assuming a TD ground truth (**Table S4**). On the other hand, when using a Gaussian decay spatial weighting function, the range of errors was significantly reduced, ranging from 16–38% error from true *R*_*t*_ values (**Table 1**). When the ground truth was assumed to be temperature-independent, both Gaussian and exponential spatial models produced comparatively similar and moderate errors, indicating that baseline approaches perform reasonably well when temperature does not influence transmission dynamics (**Table 1**).

To better understand the sources of these errors, we decomposed the mean squared error into variance and squared bias components across simulations. This decomposition showed that the majority of the error was driven by variance rather than bias in most scenarios. Bias was highest at the beginning and end of simulated epidemics, when the number of infectious individuals and candidate infector-infectee pairs was small, leading to structural constraints on estimation and systematic deviations from the true *R*_*t*_. In contrast, during the middle phases of the epidemic, when the number of infectious individuals was larger and the set of candidate transmission links expanded, variance dominated the error, reflecting increased stochasticity in infector assignment under the likelihood-based reconstruction framework. Variance was substantially higher in simulations using the exponential decay kernel compared to those using the Gaussian kernel, likely reflecting the fact that the exponential kernel produces a broader set of possible transmission pairs due to its heavier tail, increasing uncertainty in infector-infectee assignment. Both Eq (11) and Eq (12) capture the decreased probability of successful transmission as distance grows and incorporate average *Aedes* flight ranges,^33^ but they differ in shape. Eq (11) assigns higher transmission probability weights to nearby individuals, and very small weights to those further away, narrowing the candidate infector pool and reducing stochastic variability in the reconstructed transmission network. This strong spatial localization in the Gaussian kernel limits the effective set of candidate infectors, therefore reducing the propagation of temporal uncertainty into transmission reconstruction. On the contrary, Eq (12) begins with lower weights assigned to nearby individuals and declines more gradually. So, under the exponential decay kernel, a larger set of spatially plausible infector-infectee pairs allows temporal uncertainty to propagate across a wider network of candidate transmission trees, amplifying variance. Consequently, although correctly specified temperature-dependent models reduce bias, the additional variance introduced under narrow temperature ranges or broader spatial kernels may outweigh these gains, such that simpler or partially misspecified models with slightly higher bias can nevertheless achieve lower overall error through reduced variance. This likely explains the counterintuitive pattern observed in **Table 1**, where correctly specified generation time models did not consistently yield the lowest errors.

A similar mechanism explains the elevated variance observed in temperature-dependent ground truth simulations. Unlike temperature-independent generation time assumptions, which assign near-zero transmission probabilities for infections temporally close (e.g., 1–7 days apart), the temperature-dependent formulation retains non-zero transmission probabilities even for short generation times (e.g., 2–3 days under high-temperature conditions). This increases the number of temporally plausible infectors per case, consequently expanding the number of candidate transmission pathways and increasing stochastic variability in *R*_*t*_. When combined with spatial uncertainty, this leads to substantially higher variance, which explains the elevated mean absolute errors and percent errors for estimated *R*_*t*_ observed in the exponential decay, temperature-dependent ground truth simulations.

By explicitly accounting for the impact of daily temperatures on the extrinsic incubation period, we captured a biologically realistic driver of *R*_*t*_ fluctuations. For instance, an increase in temperature would shorten dengue’s incubation period in mosquitoes, temporarily boosting transmission potential.^24,42^ Mechanistically, shorter EIPs would be expected to lead to infections occurring sooner after a mosquito becomes infectious, which could increase the likelihood of transmission becoming more localized given the mosquitoes’ limited flight range, potentially favoring spatial kernels that assign higher weights to pairs of nearby individuals. Our estimation method detects temperature-driven shifts in transmission timing, whereas conventional approaches averaging generation times across seasons would overlook them. Specifically, a static generation time may obscure the shortening and lengthening of the EIP during warmer and colder seasons.

We had hypothesized that for warmer temperatures (i.e., shorter EIPs), infections would be more localized and thus a spatial kernel assigning higher weights to shorter distances would better capture true transmission dynamics, which we observed from the lower mean errors and narrower variability in errors yielded by the Gaussian decay spatial kernel. If spatial heterogeneity is ignored and homogeneous mixing is assumed, transmission from clustered cases is blurred across the entire population. This obscures localized clusters with higher spread, leading to an underestimation of *R*_*t*_ in “hotspots” and an overestimation elsewhere. *Ae. aegypti* and *albopictus* are highly localized near their breeding sites due to their shorter flight ranges,^38^ and areas with dense vegetation or high precipitation have shown an association with dengue case clusters,^25^ thus potentially leading to large differences in transmission potential even between adjacent neighborhoods. This misestimation could lead to the inadequate allocation of interventions and resources between higher- and lower-risk areas.

To further test these findings, we explored several major assumptions. The Gaussian decay spatial weighting function’s (Eq (11)) superior performance held across sensitivity analyses. Temperature dependence without consideration of spatial heterogeneity resulted in large percent errors, suggesting that incorporating only temperature and not the distance between individuals can mislead transmissibility estimates. This phenomenon is likely observed because the simulated epidemics were spatially structured, with many potential infectors located far from susceptible individuals. In non-spatial models, all pairs of individuals are treated as equally likely to transmit infection regardless of how far apart they are. By contrast, even imperfect parameterizations of the exponential or Gaussian decay spatial weighting functions (Eqs 11, 12) weight transmission probability by the actual distance between cases, correctly assigning lower weights to long-distance transmission and higher weights to short-distance transmission. When temperature was not part of the ground truth, incorporating temperature dependence had minimal impact under the Gaussian kernel but increased error under the exponential kernel, suggesting that additional temporal flexibility is well-tolerated under appropriate spatial constraints but may introduce unnecessary variability when spatial structure is misspecified.

Application of the models to empirical 2019 dengue case data in Singapore showed that the temperature-dependent spatiotemporal approaches developed in this study produced *R*_*t*_ estimates that were frequently near or below 1 and generally lower in magnitude than those from EpiFilter during several periods (**Figure 3B–E**). All WT-based models produced comparatively variable trajectories, with estimates oscillating between 0 and ∼3 over time, whereas EpiFilter showed smoother temporal patterns, reflecting the more sensitive responsiveness of the WT-based models to short-term changes in observed incidence. The difference in fluctuations between approaches is consistent with their underlying formulations. In this study, *R*_*t*_ represents an instantaneous reproduction number derived from individual-level transmission probabilities rather than a cohort-averaged quantity over a generation interval. Each individual’s reproduction number depends on pairwise infection probabilities, *p* ∝ *f* (*t*_*k*_ − *t*_*l*_ | *θ*_*f*_(*T*)) · *g*(*d*_*k*_ − *d*_*l*_ | *θ*_*g*_), so if either the temporal, *f*(·), or spatial, *g*(·), components assign very small weights to most infector-infectee pairs, then *R*_*l*_ (and consequently, *R*_*t*_) will approach 0 when cases are temporally or spatially isolated. Thus, stochastic variation in incidence propagates directly into *R*_*t*_, leading to more variable trajectories. Unlike renewal-based approaches, WT-style estimates quantify transmission potential at the individual level and can therefore respond more rapidly to short-term changes in transmission. However, in the present application, the WT-based trajectories were dominated by substantial short-term stochastic variability, and due to the relatively low number of cases, we did not observe clear consistency between incidence dynamics and *R*_*t*_ patterns. In contrast, EpiFilter formulates *R*_*t*_ as a latent stochastic process evolving through a state space model, with observed incidence linked via a renewal equation and estimates updated through Bayesian forward-backward smoothing. The tendency of early EpiFilter *R*_*t*_ estimates to approach *R*_*max*_ likely reflects edge effects caused by sparse early incidence and low initial infectiousness rather than true transmission intensity. When case counts are sparse, this framework tends to regularize estimates towards the prior, producing smoother trajectories. Differences among the exponential, Gaussian, and non-spatial WT models were modest in terms of mean *R*_*t*_ at the aggregated scale. However, the Gaussian decay model yielded comparatively less extreme fluctuations than the exponential kernel, consistent with simulation findings that more concentrated spatial weighting can limit the propagation of space-time variability. Overall, in this empirical application, spatial structure influenced the stability and dispersion of estimates more than it altered average *R*_*t*_ levels.

These findings elaborate upon a growing body of work on quantifying the transmissibility of climate-sensitive pathogens. Earlier studies established the role of temperature in vector-borne pathogens’ EIP,^42^ but formally simulating this phenomenon to estimate *R*_*t*_ has lagged. Spatially, we reproduced a previous spatially explicit framework,^21^ but introduced a temperature-dependent generation time, synthesizing two separate dimensions of transmission. Our temperature-dependent spatiotemporal model demonstrated high accuracy in predicting whether *R*_*t*_ exceeded or fell below the epidemic threshold of 1, as reflected by consistently high ROC AUC values across simulations (**Table 1, Figure S4, S5**). However, while the framework reliably captured epidemic growth or decline, it was more challenging to recover precise *R*_*t*_ magnitudes and to correctly reconstruct individual transmission links, reflecting the inherent stochastic uncertainty in identifying exact infector-infectee pairs within complex mosquito-borne transmission networks. The high level of noise in the WT-based models observed in its error decomposition and empirical application (**Figure 3**) suggests limited resourcefulness in real-time monitoring of epidemic growth or decline compared to smoothing approaches such as that of EpiFilter, so they may be better suited for retrospective analyses and individual-level transmission heterogeneity.

Certain limitations were present in the models developed in this study. First, temperature snapshots at the time of case reporting were used to construct the temperature-dependent generation time distribution because the exact timing at which temperature affects the temperature-dependent EIP occurs is unknown. We assumed temperature to directly affect the rate parameter, and therefore the mean, of the gamma distribution representing the EIP (Eq (10)), while the IIP and biting intervals remain temperature independent. In reality, the EIP occurs after the human incubation period and a subsequent delay for mosquito biting, so the temperature on the actual day the mosquito becomes infected could differ from the snapshot used. This could alter the mean and shape of the generation time distribution for that individual, potentially affecting the estimated reproduction number. However, temperature in tropical and subtropical regions does not normally fluctuate much within a single week, so results would not be significantly altered. Second, we assumed a single pair of coordinates per individual, which means each person has a static location, and we disregarded human mobility. Certain studies on vector-borne disease dynamics have shown that human mobility may affect transmission potential, and future models may benefit from incorporating this granularity.^43,44^ Although research in this direction remains scarce and has mostly been applied to human-to-human transmission infections such as COVID-19,^22,45^ incorporating human mobility into the present model could more accurately infer “who infected whom” thereby further improving the accuracy of *R*_*t*_ estimates, even for vector-borne diseases. However, incorporating mobility data comes with practical and computational challenges, particularly given the limited availability of fine-scale movement data in dengue-endemic settings. As an initial step, our static-location framework provides a tractable approximation that can later be extended as richer mobility datasets become available. If human mobility data were available, future research could incorporate dynamic coordinates into the spatial weighting function, allowing transmission likelihoods to be modeled continuously across individuals’ movement patterns. Moreover, our model implicitly assumes that the recorded location corresponds to the place of infection. In reality, surveillance data more often capture place of residence, which introduces a source of measurement error and may bias spatial inference if the true site of exposure differs from where cases live. Finally, intervention-free simulations omit how vector control tools such as *Wolbachia* releases can truncate transmission networks, potentially overestimating *R*_*t*_ in real-world settings.^1–4^ These gaps do not invalidate our core findings but highlight contextual dependencies for implementation.

The strengths of the present study lie in its biologically logical input parameters and easily interpretable outputs. We incorporated viral dynamics, which are affected by temperature, within vectors into generation times, making *R*_*t*_ estimates interpretable mechanistically. For example, a wave of warmer temperature could indicate an upcoming spike in *R*_*t*_. Spatial explicitness also enables township-level risk stratification. Public health interventions could prioritize areas where *R*_*t*_ is persistently >1 despite citywide declines. Such granularity is feasible in practice: temperature data are widely available, and cases are routinely surveilled in vector-borne disease-endemic nations. Additionally, model performance did not decrease drastically even after applying it to wider temperature data with seasonal fluctuations (**Figure S1**), meaning the model can be generalizable outside of a Singapore-specific context. Holding all other metrics equal, the mean percent error for *R*_*t*_ using Kaohsiung, Taiwan’s mean daily temperature data (where winter months are colder and summer months are warmer), was 33% (20–50%), whereas a more consistent set of temperature data such as Singapore’s led to a mean percent error of 27% (16–38%). These results suggest that the framework remains nearly equally robust across different climatic settings, which is crucial for informing whether special public health intervention planning is needed for regions that experience wider seasonal fluctuations or not.

For public health, this work invites immediate applications. Currently ongoing interventions aimed to curb vector-borne transmission, such as *Wolbachia*-infected mosquito releases to reduce dengue transmission,^1–4^ could benefit from models that more accurately estimate *R*_*t*_ to guide intervention schedules and prioritize locations that would most benefit from the intervention. Future research should test this in diverse settings, such as Brazil’s urban-rural setting, incorporate human movement data (e.g., mobile-derived mobility networks), and merge with immunity-aware models to reflect population susceptibility. Ultimately, refining *R*_*t*_ estimation can help build surveillance systems that see epidemics as dynamic, spatially explicit, and environmentally mediated to better reflect vector-borne disease dynamics observed in the real world.

## Supplementary Materials

**Table S1.** List of parameters used and their associated values.

| Parameter | Description | Value |
| --- | --- | --- |
| $\alpha_{IIP}$ | Shape parameter of the intrinsic incubation period (IIP); gamma distribution | $\alpha_{IIP} = 16$ |
| $\lambda_{IIP}$ | Rate parameter of the IIP; gamma distribution | $\lambda_{IIP} = 2.7 \text{ day}^{-1}$ |
| $\mu_{IIP}$ | Mean of the gamma-distributed IIP | $\mu_{IIP} = 5.93 \text{ days}$ |
| $\alpha_{EIP}$ | Shape parameter of the extrinsic incubation period (EIP); gamma distribution | $\alpha_{EIP} = 4.3$ |
| $\lambda_{EIP}(T) = (\beta_0 - \beta_1 T)^{-1}$ | Temperature-dependent rate parameter of the EIP; gamma distribution | $\beta_0 = 7.9, \beta_1 = -0.21 \text{ day}^{-1}$ |
| $\lambda_{MH}$ | Rate parameter of the mosquito-to-human transmission period; exponential distribution | $\lambda_{MH} = 1 \text{ day}^{-1}$ |
| $\lambda_{HM}$ | Rate parameter of the human-to-mosquito transmission period; exponential distribution | $\lambda_{HM} = 1 \text{ day}^{-1}$ |
| $T$ | Set of daily temperature values in 2019 | Singapore: 24.9–30.1°C<br>Kaohsiung, Taiwan: 16.8–30.9°C |
| $M$ | *Ae. aegypti* maximum flight range | 106 meters |

**Table S2.**
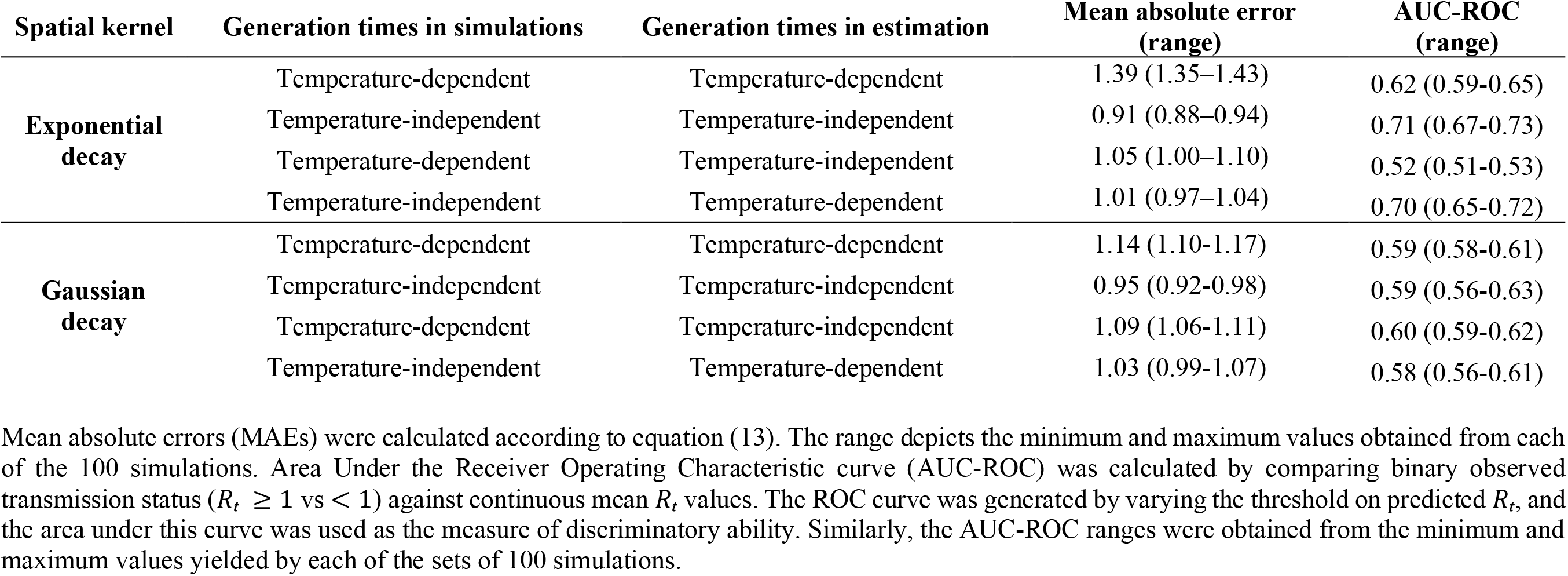
Mean absolute errors of *R*_*l*_ and AUC-ROC scores across different simulation and estimation inputs.

**Table S3.** Mean absolute errors, percent errors and AUC-ROC scores of *R*_*t*_ across different parameter values and estimation inputs (exponential decay spatial kernels).

| Maximum distance (m) | Maximum generation time (days) | Generation time distribution | Mean MAE | MAE Range | Mean Error (%) | Error Range (%) | Mean AUC | AUC Range |
| --- | --- | --- | --- | --- | --- | --- | --- | --- |
| 500 | 28 | GT1 | 0.73 | 0.44–1.34 | 50.77 | 30.45–78.3 | 0.93 | 0.69–0.99 |
|  |  | GT2 | 0.4 | 0.28–0.66 | 27.67 | 20.43–40.16 | 0.92 | 0.68–0.99 |
|  |  | GT3 | 0.47 | 0.27–0.91 | 32.46 | 19.57–53.21 | 0.92 | 0.7–0.99 |
|  | 35 | GT1 | 0.73 | 0.44–1.42 | 50.98 | 30.58–83.12 | 0.93 | 0.7–0.99 |
|  |  | GT2 | 0.39 | 0.28–0.64 | 27.45 | 20.09–40.42 | 0.92 | 0.69–0.99 |
|  |  | GT3 | 0.47 | 0.27–0.95 | 32.65 | 19.13–55.55 | 0.92 | 0.7–0.98 |
|  | 42 | GT1 | 0.73 | 0.43–1.34 | 50.79 | 30.04–78.72 | 0.93 | 0.7–0.99 |
|  |  | GT2 | 0.39 | 0.28–0.68 | 27.62 | 21.24–39.69 | 0.92 | 0.69–0.99 |
|  |  | GT3 | 0.47 | 0.27–0.92 | 32.46 | 18.95–53.68 | 0.92 | 0.69–0.99 |
| 1000 | 28 | GT1 | 1.65 | 0.23–3.63 | 112.3 | 20.45–216.74 | 0.95 | 0.88–0.99 |
|  |  | GT2 | 0.44 | 0.2–0.85 | 30.17 | 14.58–49.62 | 0.94 | 0.87–0.99 |
|  |  | GT3 | 0.83 | 0.23–1.71 | 56.86 | 20.35–99.54 | 0.95 | 0.86–0.99 |
|  | 35 | GT1 | 1.67 | 0.25–3.8 | 113.21 | 21.86–226.27 | 0.95 | 0.88–0.99 |
|  |  | GT2 | 0.43 | 0.19–0.78 | 29.95 | 14.44–50.34 | 0.94 | 0.87–0.99 |
|  |  | GT3 | 0.83 | 0.23–1.74 | 57.05 | 20.33–101.42 | 0.95 | 0.87–0.99 |
|  | 42 | GT1 | 1.67 | 0.24–3.7 | 113.28 | 21.23–230.51 | 0.95 | 0.88–0.99 |
|  |  | GT2 | 0.44 | 0.19–0.81 | 30.12 | 14.39–51.5 | 0.94 | 0.87–0.99 |
|  |  | GT3 | 0.83 | 0.22–1.63 | 56.66 | 19.91–94.92 | 0.94 | 0.87–0.99 |
| 1500 | 28 | GT1 | 3 | 0.35–6.33 | 204.57 | 31.07–383.74 | 0.96 | 0.9–1 |
|  |  | GT2 | 0.44 | 0.2–0.83 | 30.4 | 14.59–53.2 | 0.95 | 0.88–0.99 |
|  |  | GT3 | 0.94 | 0.23–1.96 | 64.51 | 20.1–114.66 | 0.96 | 0.89–0.99 |
|  | 35 | GT1 | 2.98 | 0.35–6.08 | 203.27 | 31.05–363.34 | 0.97 | 0.9–1 |
|  |  | GT2 | 0.44 | 0.2–0.84 | 30.35 | 14.18–51.79 | 0.95 | 0.88–0.99 |
|  |  | GT3 | 0.94 | 0.21–2 | 64.31 | 18.71–116.55 | 0.96 | 0.89–0.99 |
|  | 42 | GT1 | 3 | 0.34–5.94 | 204.56 | 30.67–365.26 | 0.96 | 0.9–1 |
|  |  | GT2 | 0.44 | 0.21–0.87 | 30.43 | 14.95–50.99 | 0.95 | 0.88–0.99 |
|  |  | GT3 | 0.94 | 0.2–1.86 | 64.57 | 18.19–108.95 | 0.96 | 0.89–0.99 |

**Table S4.** Mean absolute errors, percent errors and AUC-ROC scores of *R*_*t*_ across different parameter values and estimation inputs (Gaussian decay spatial kernels).

| Maximum distance (m) | Maximum generation time (days) | Generation time distribution | Mean MAE | MAE Range | Mean Error (%) | Error Range (%) | Mean AUC | AUC Range |
| --- | --- | --- | --- | --- | --- | --- | --- | --- |
| 500 | 28 | GT1 | 0.32 | 0.19–0.45 | 27.9 | 15.81–37.8 | 0.82 | 0.57–0.96 |
|  |  | GT2 | 0.25 | 0.21–0.32 | 22.2 | 18.68–28.98 | 0.79 | 0.56–0.93 |
|  |  | GT3 | 0.25 | 0.19–0.34 | 21.5 | 16.7–28.58 | 0.79 | 0.61–0.93 |
|  | 35 | GT1 | 0.32 | 0.21–0.45 | 27.8 | 17.06–36.66 | 0.82 | 0.57–0.95 |
|  |  | GT2 | 0.25 | 0.21–0.31 | 22.2 | 18.73–28.93 | 0.79 | 0.57–0.93 |
|  |  | GT3 | 0.25 | 0.2–0.31 | 21.5 | 16.89–28.37 | 0.79 | 0.61–0.92 |
|  | 42 | GT1 | 0.32 | 0.19–0.43 | 27.8 | 15.47–37.33 | 0.82 | 0.58–0.96 |
|  |  | GT2 | 0.25 | 0.21–0.32 | 22.2 | 18.87–28.71 | 0.79 | 0.57–0.93 |
|  |  | GT3 | 0.25 | 0.2–0.33 | 21.5 | 17.4–28.12 | 0.79 | 0.61–0.93 |
| 1000 | 28 | GT1 | 0.31 | 0.18–0.42 | 26.8 | 15.87–36.53 | 0.82 | 0.73–0.92 |
|  |  | GT2 | 0.26 | 0.21–0.32 | 22.5 | 18.48–26.49 | 0.78 | 0.64–0.88 |
|  |  | GT3 | 0.25 | 0.2–0.34 | 21.6 | 17.65–28.33 | 0.78 | 0.66–0.87 |
|  | 35 | GT1 | 0.31 | 0.18–0.44 | 26.9 | 16.01–36.1 | 0.82 | 0.73–0.92 |
|  |  | GT2 | 0.26 | 0.2–0.33 | 22.6 | 18.09–27.45 | 0.78 | 0.65–0.88 |
|  |  | GT3 | 0.25 | 0.2–0.33 | 21.6 | 18.01–27.53 | 0.78 | 0.66–0.87 |
|  | 42 | GT1 | 0.31 | 0.18–0.43 | 27 | 16.2–37.69 | 0.82 | 0.72–0.92 |
|  |  | GT2 | 0.26 | 0.2–0.32 | 22.5 | 18.28–26.6 | 0.78 | 0.64–0.87 |
|  |  | GT3 | 0.25 | 0.2–0.33 | 21.6 | 17.41–27.65 | 0.78 | 0.66–0.88 |
| 1500 | 28 | GT1 | 0.31 | 0.18–0.43 | 26.9 | 15.95–36.02 | 0.82 | 0.72–0.92 |
|  |  | GT2 | 0.26 | 0.2–0.33 | 22.6 | 18.24–27.44 | 0.78 | 0.65–0.87 |
|  |  | GT3 | 0.25 | 0.2–0.36 | 21.7 | 17.56–29.24 | 0.78 | 0.66–0.88 |
|  | 35 | GT1 | 0.31 | 0.18–0.45 | 26.9 | 16.03–37.56 | 0.82 | 0.72–0.92 |
|  |  | GT2 | 0.26 | 0.2–0.33 | 22.6 | 17.75–27.21 | 0.77 | 0.65–0.87 |
|  |  | GT3 | 0.25 | 0.19–0.34 | 21.6 | 17.27–28.13 | 0.78 | 0.66–0.9 |
|  | 42 | GT1 | 0.31 | 0.18–0.42 | 26.9 | 15.56–36.65 | 0.82 | 0.72–0.92 |
|  |  | GT2 | 0.26 | 0.2–0.34 | 22.6 | 18.54–27.98 | 0.78 | 0.66–0.88 |
|  |  | GT3 | 0.25 | 0.2–0.35 | 21.7 | 18.05–28.71 | 0.78 | 0.66–0.88 |

**Table S5.** Mean absolute errors, percent errors and AUC-ROC scores of *R*_*t*_ across different parameter values and estimation inputs (non-spatial models).

| Maximum generation time (days) | Generation time distribution | Mean MAE | MAE Range | Mean Error (%) | Error Range (%) |
| --- | --- | --- | --- | --- | --- |
| 28 | GT1 | 9.98 | 4.82–16.56 | 141.3 | 66.79–257.67 |
|  | GT2 | 3.03 | 1.77–5.02 | 42.4 | 23.92–65.39 |
|  | GT3 | 4.14 | 2.39–5.92 | 58.2 | 34.74–77.88 |
| 35 | GT1 | 10.63 | 5.33–17.8 | 150.6 | 73.97–277.02 |
|  | GT2 | 3 | 1.72–4.9 | 42 | 23.68–63.86 |
|  | GT3 | 4.14 | 2.5–6.02 | 58.3 | 34.6–79.75 |
| 42 | GT1 | 10.85 | 5.46–19.7 | 153.8 | 75.79–306.5 |
|  | GT2 | 3 | 1.79–4.94 | 42.1 | 24.64–64.28 |
|  | GT3 | 4.14 | 2.35–5.91 | 58.3 | 33.65–84.59 |

Calculations were performed under a Gaussian decay spatial weighting function and a maximum distance between two consecutive cases of 1500m. $R_t$ estimates were compared against the respective true $R_t$ values for the same model.

**Figure S1.**
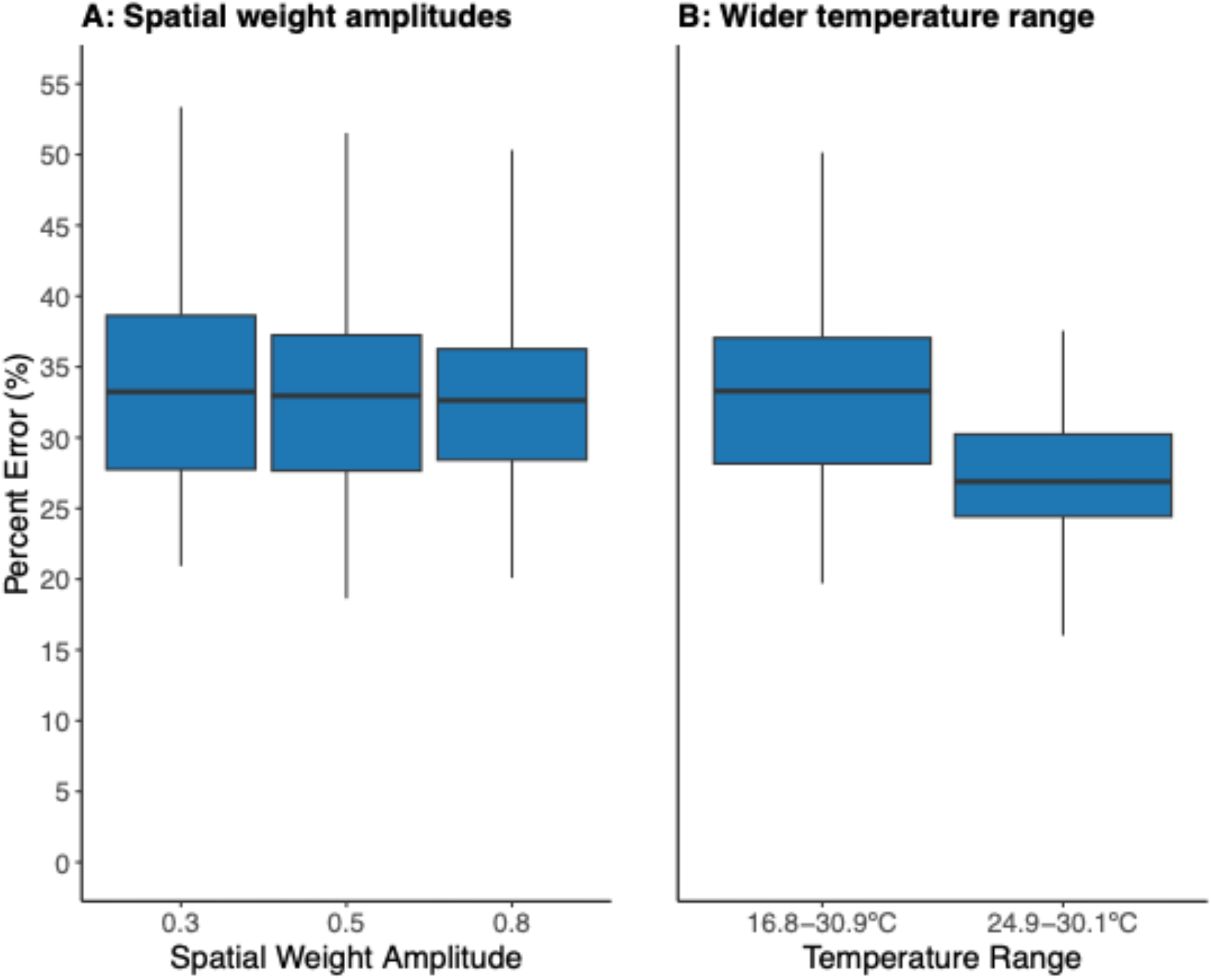
Errors in *R*_*t*_ estimation compared to true values. **A** depicts percent errors under the assumptions that the amplitude within the Gaussian decay spatial weighting function is *A* = 0.3, 0.5, and 0.8, each tested under 100 simulations, with the mean error from these simulations as the box mid-section line, the first and third quartiles as the box edges, and the minimum and maximum values as the whisker ends. **B** compares the percent errors under Singapore’s weather range (right) against a wider temperature range (left) along with the mean, first and third quartiles, and minimum and maximum values.

**Figure S2.**
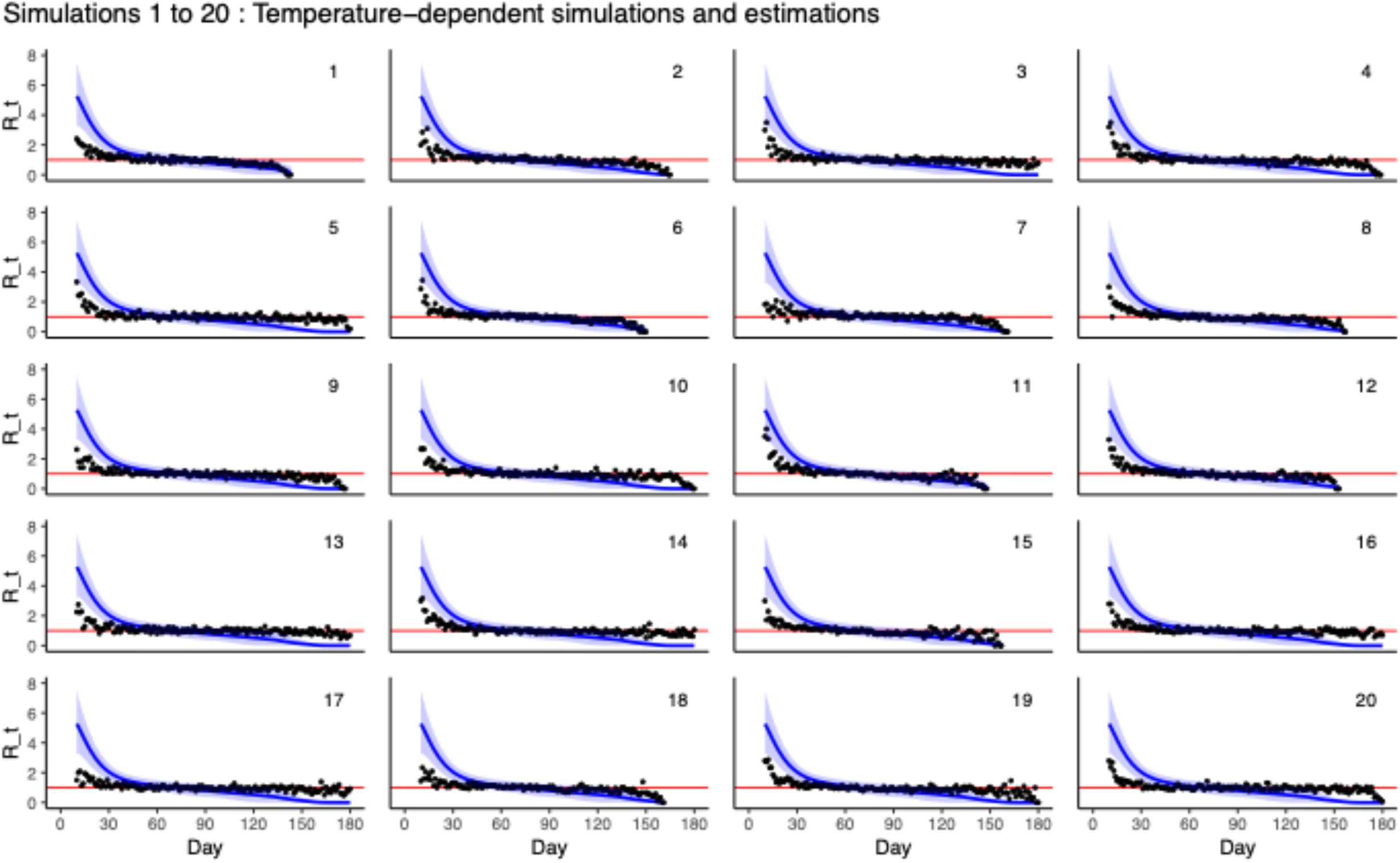
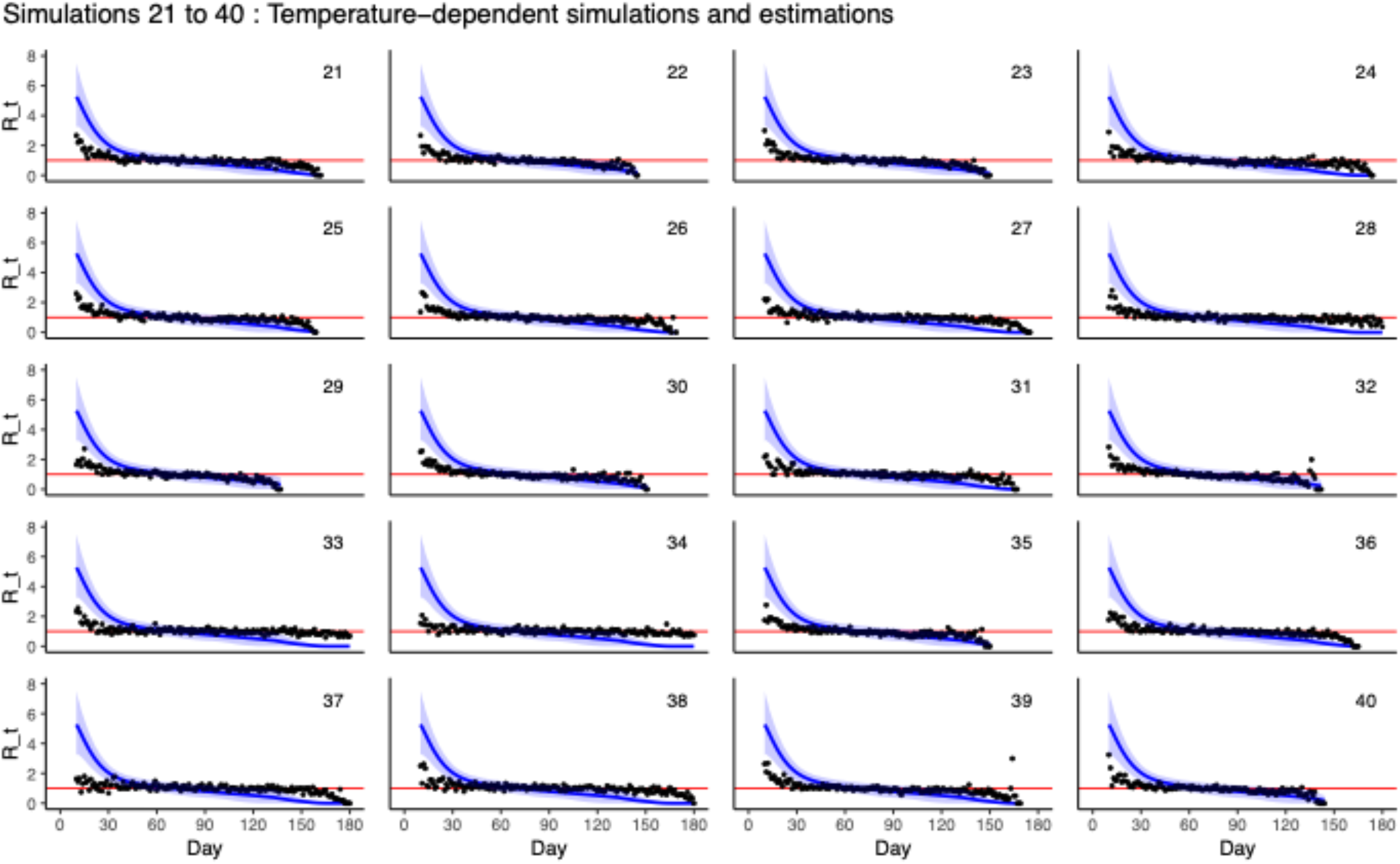
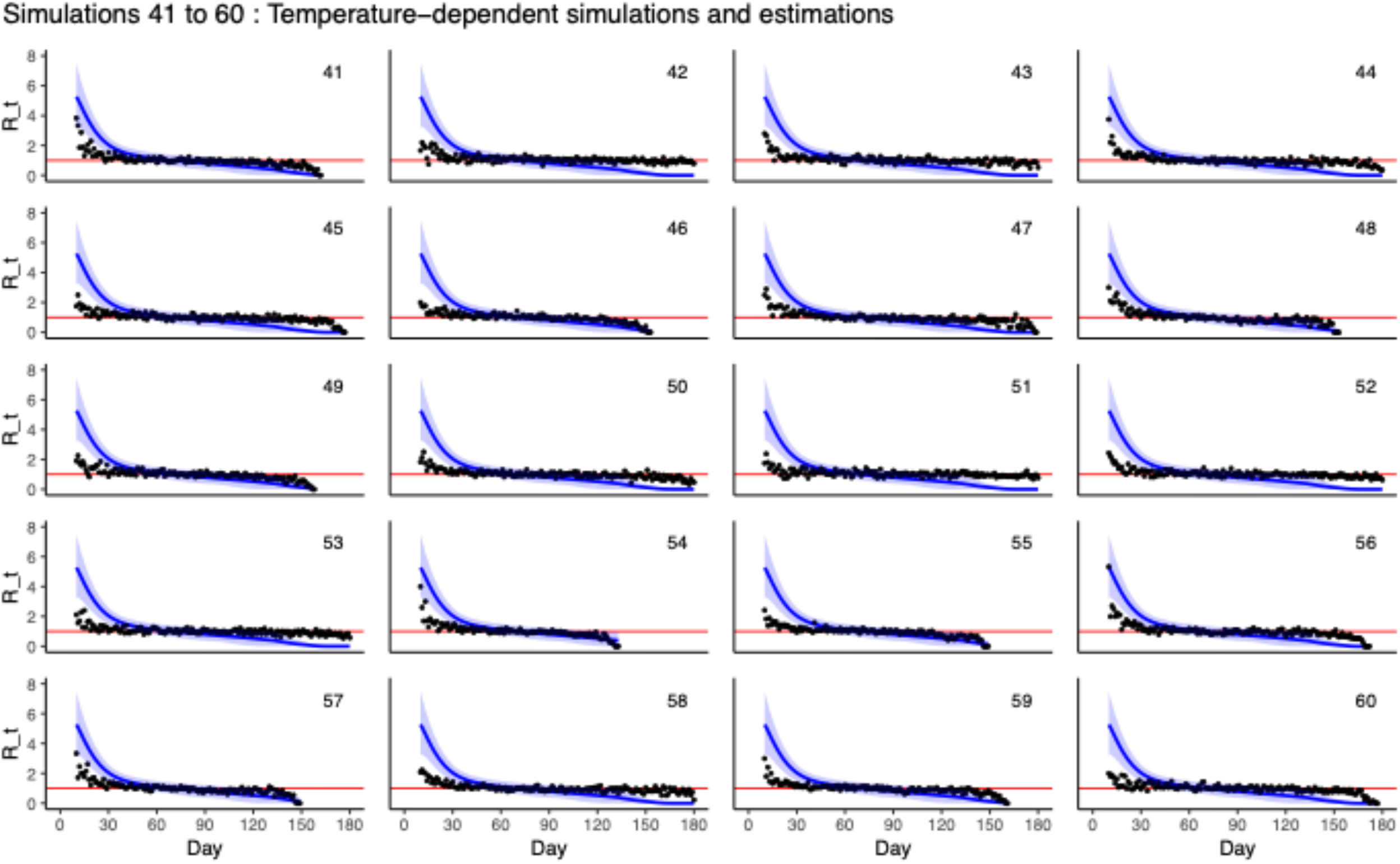
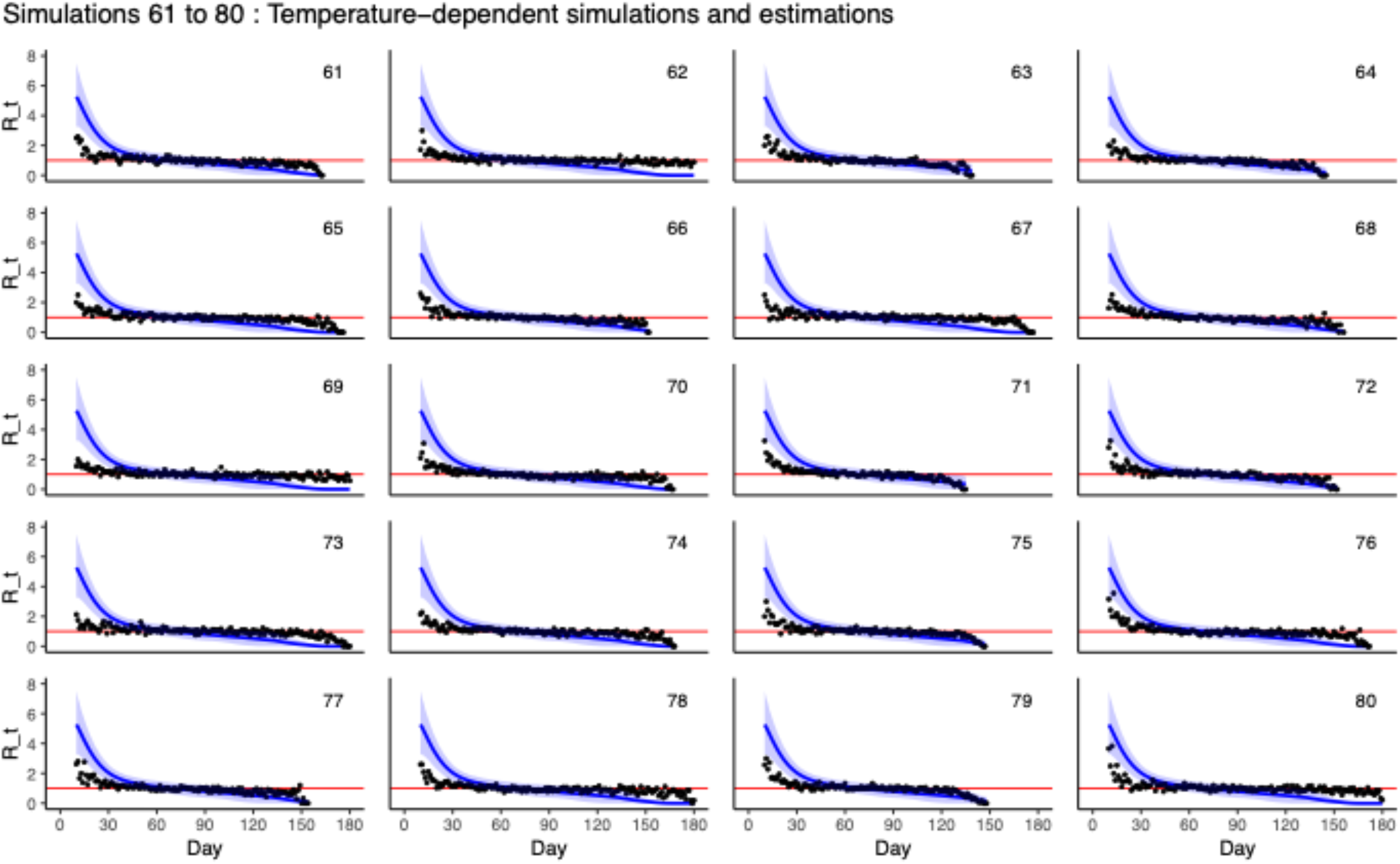
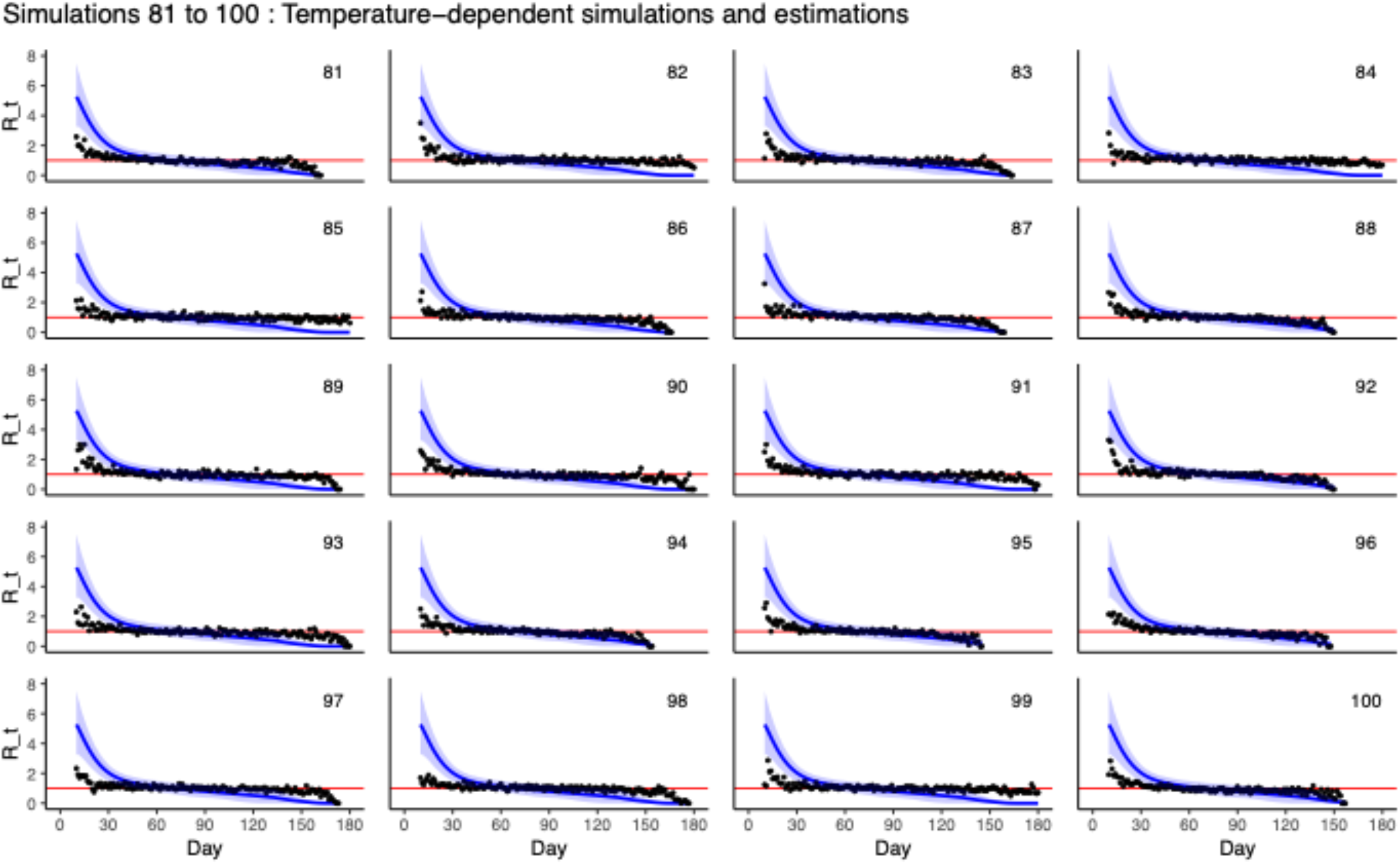
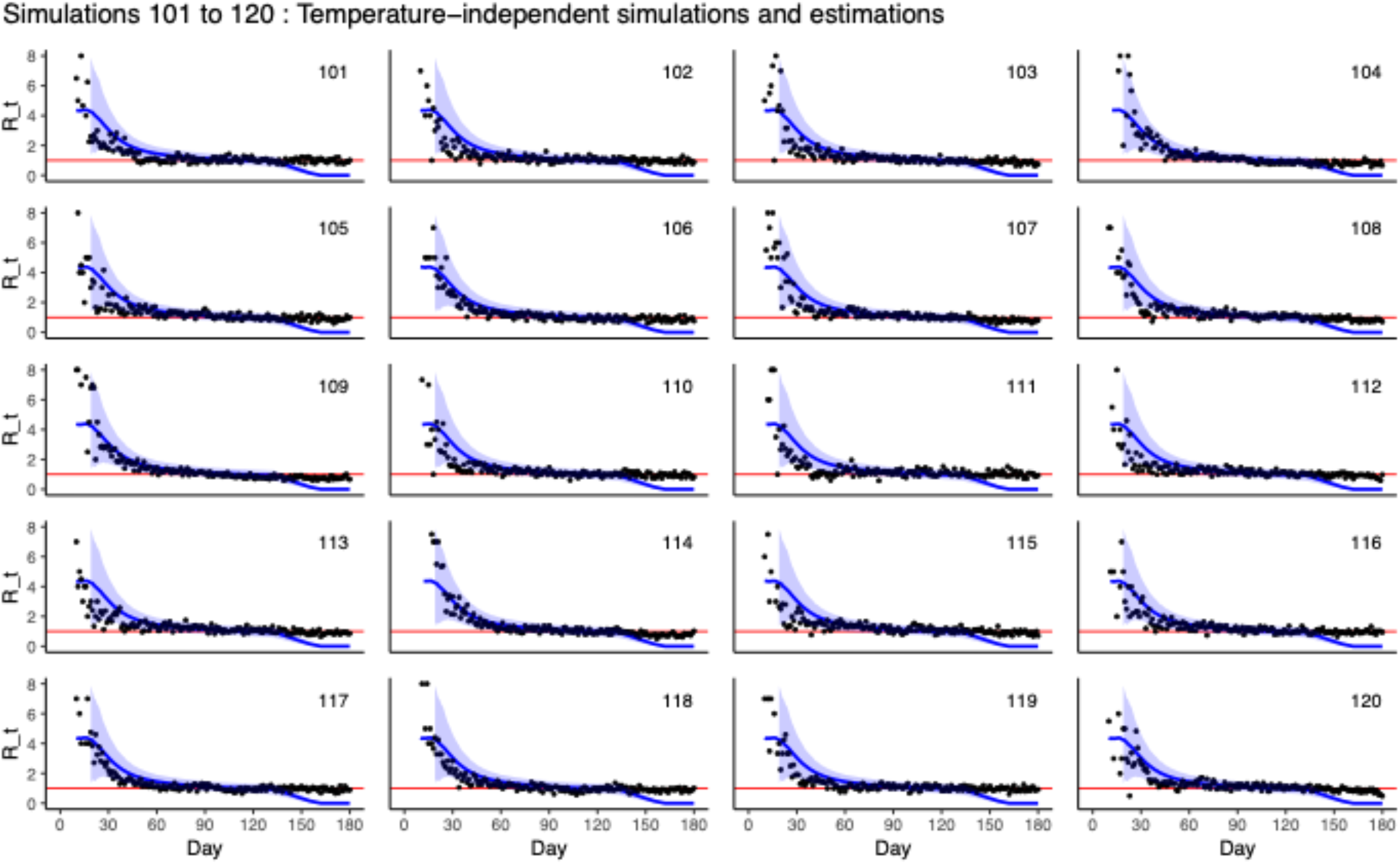
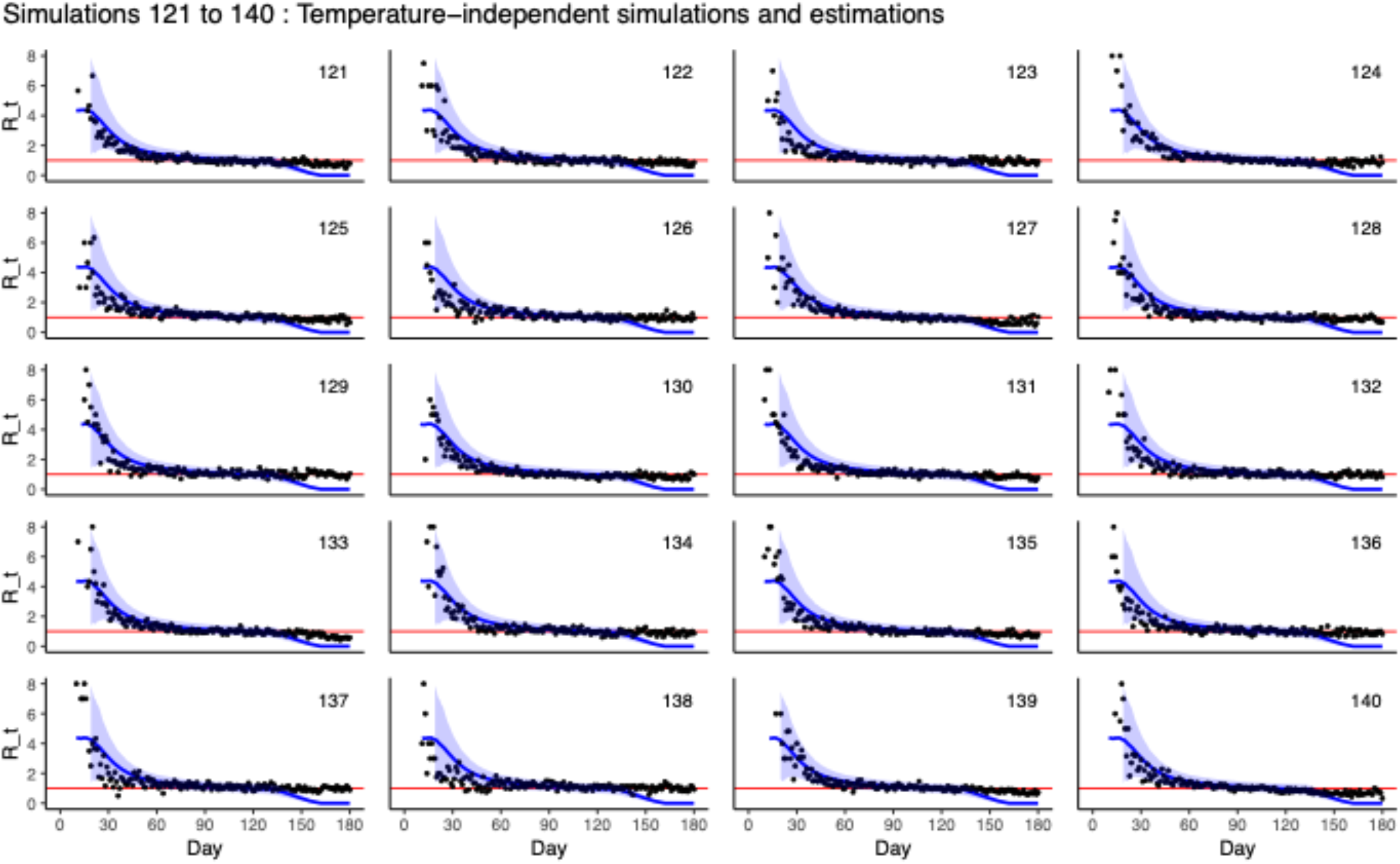
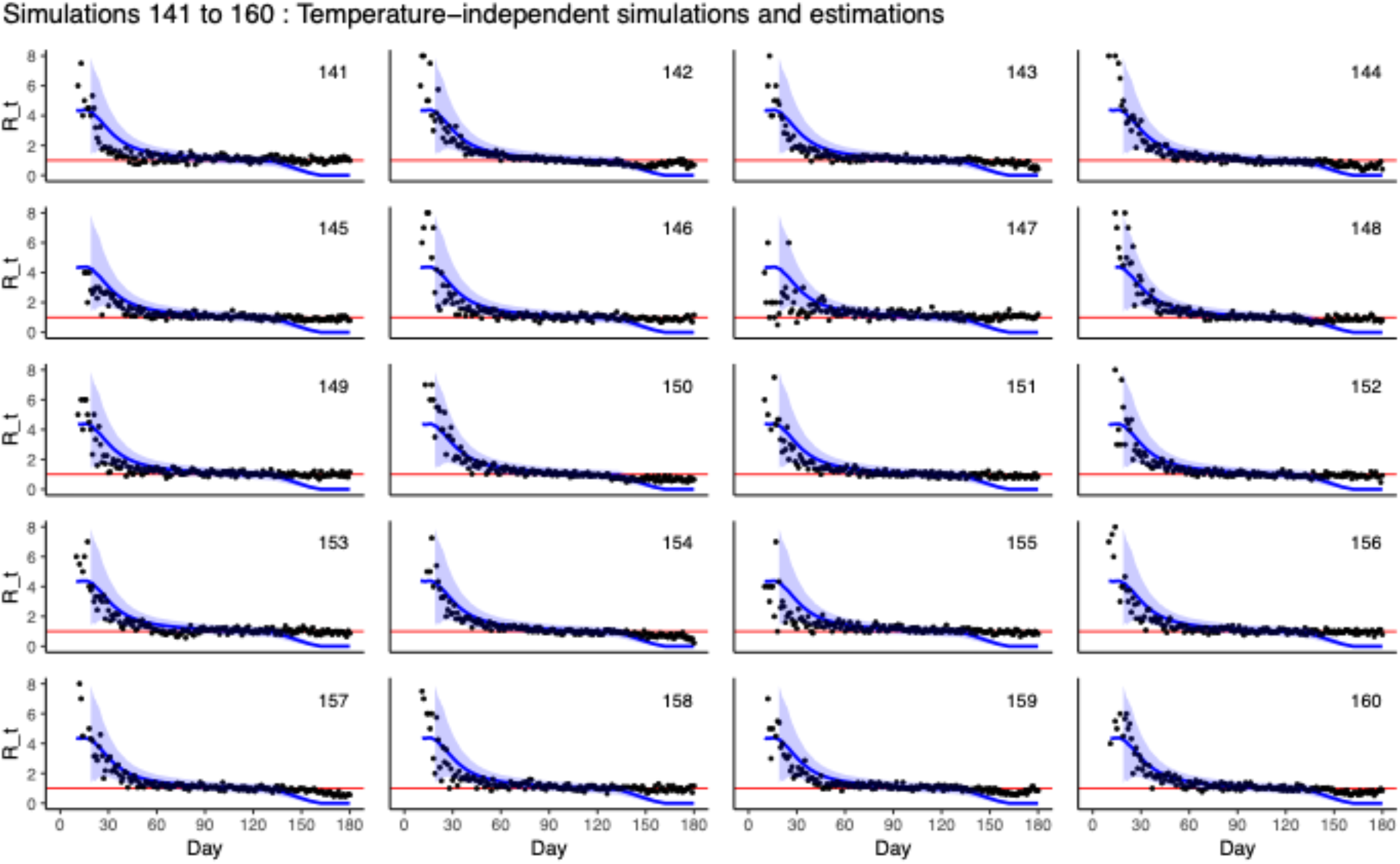
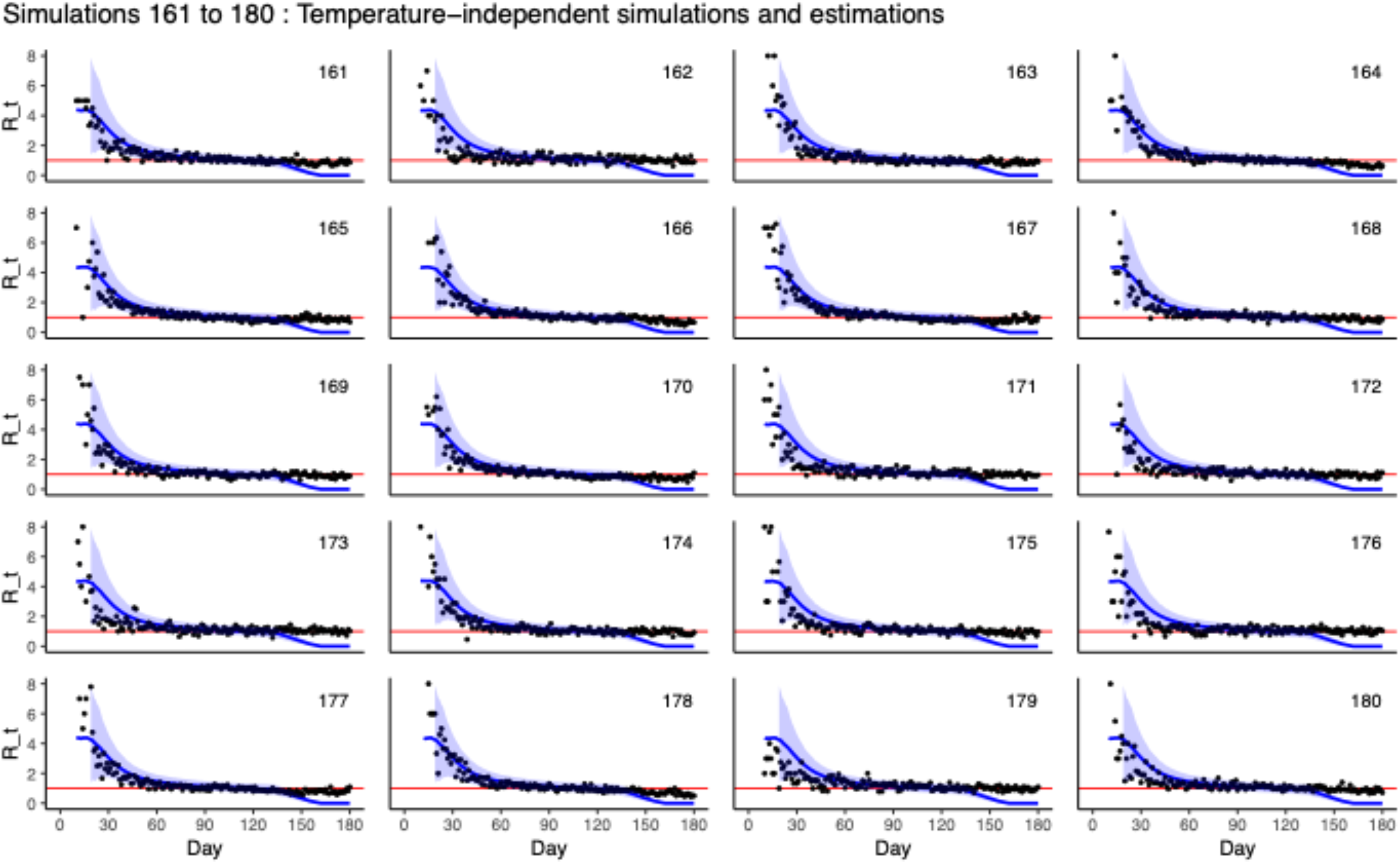
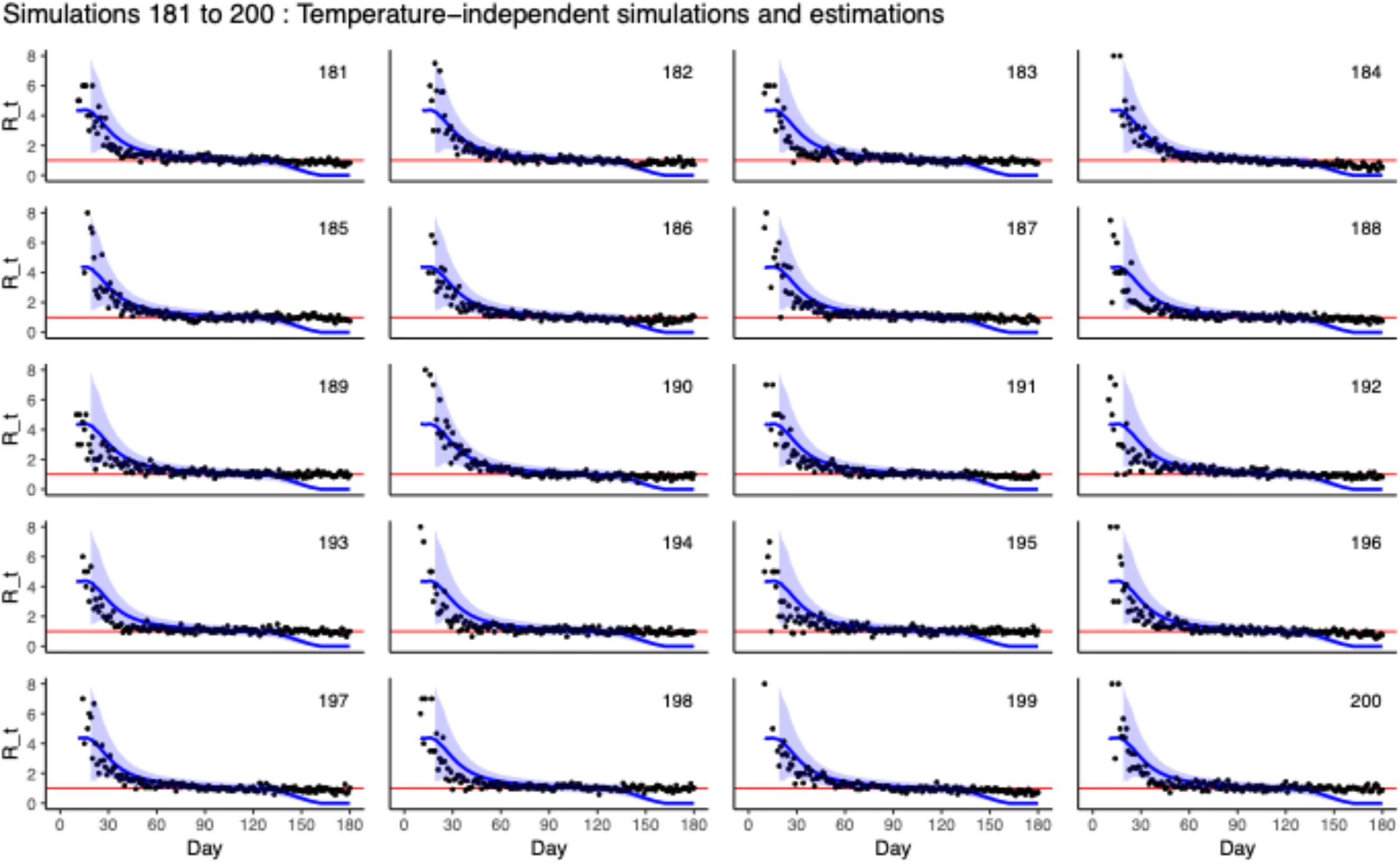
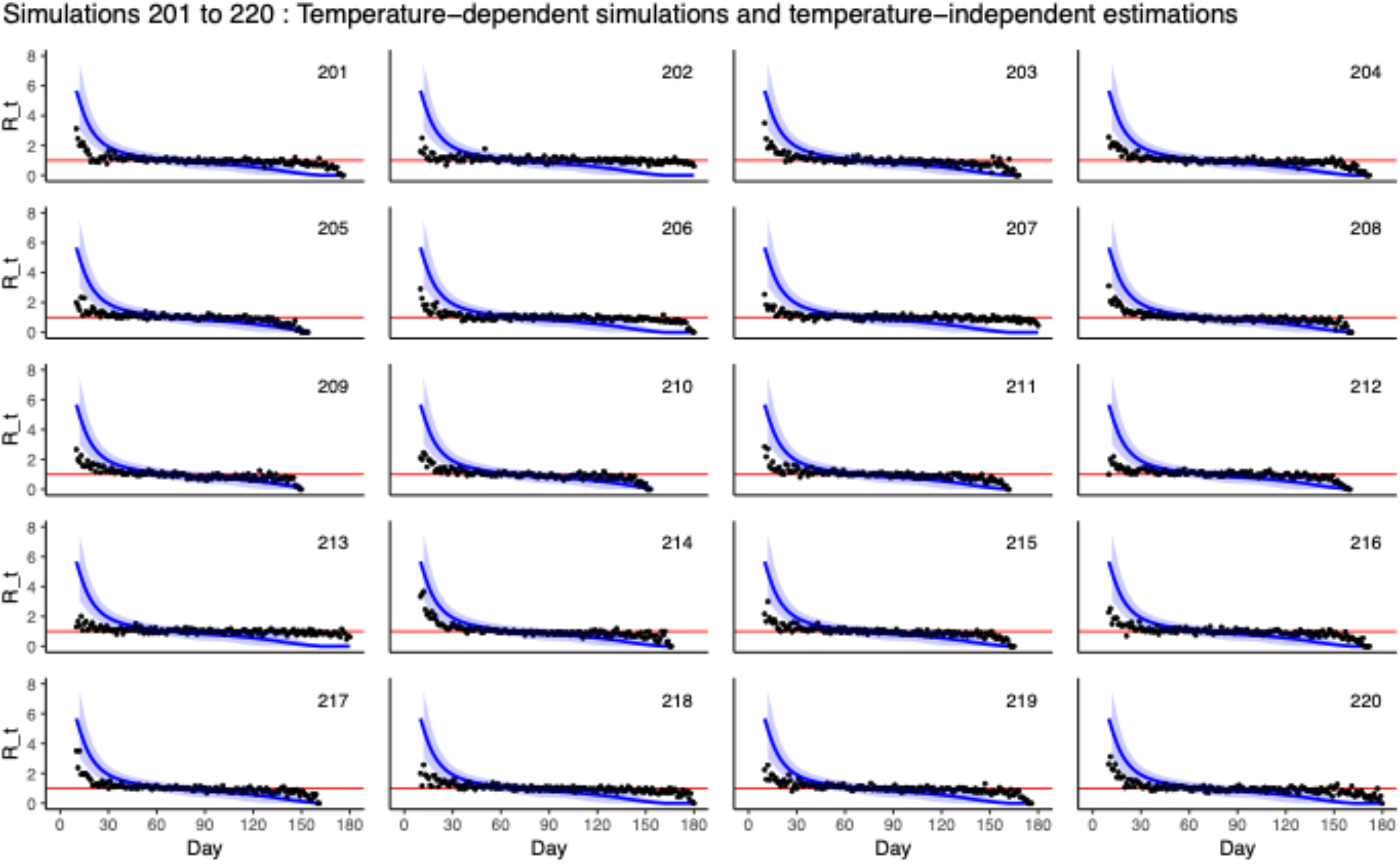
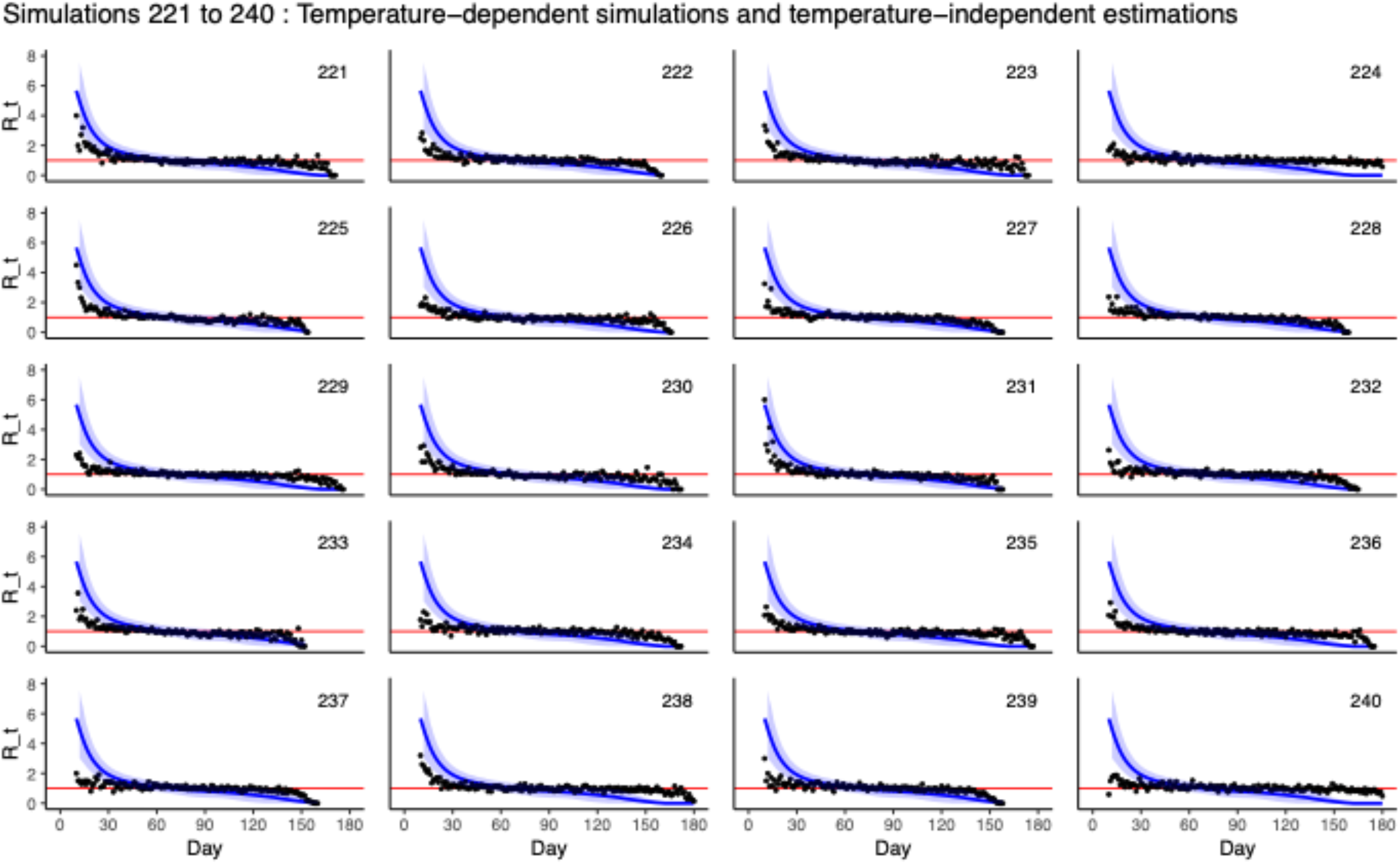
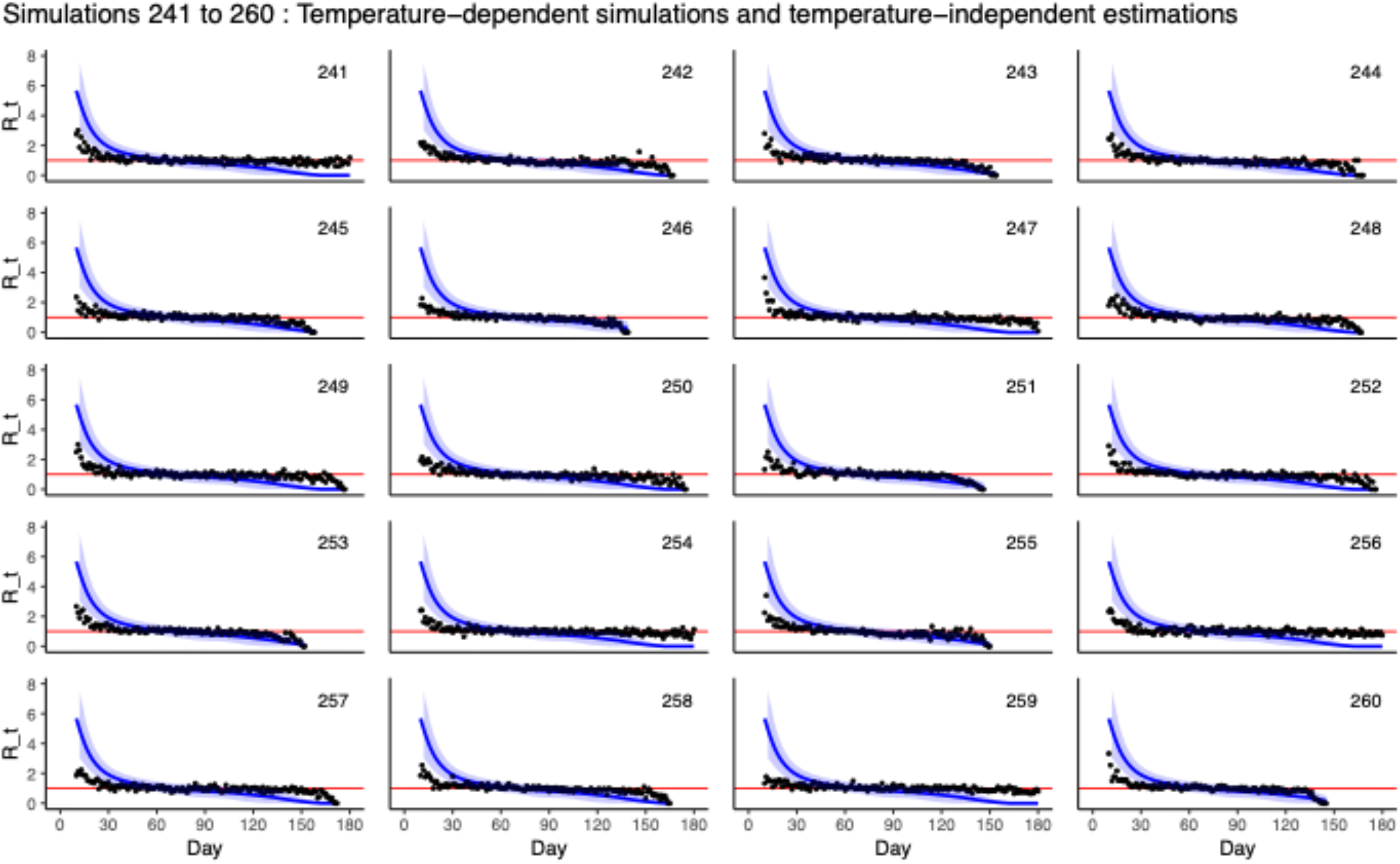
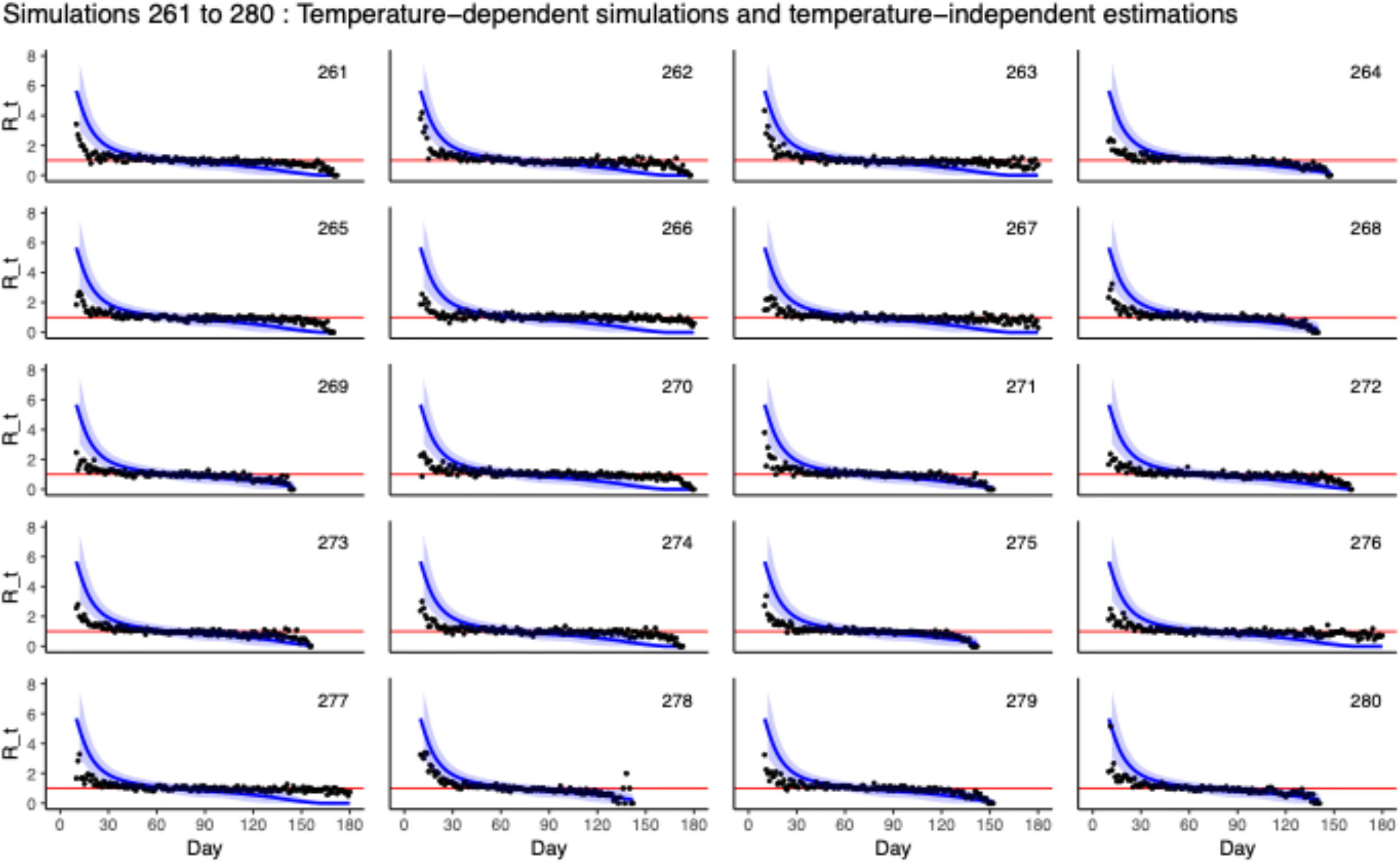
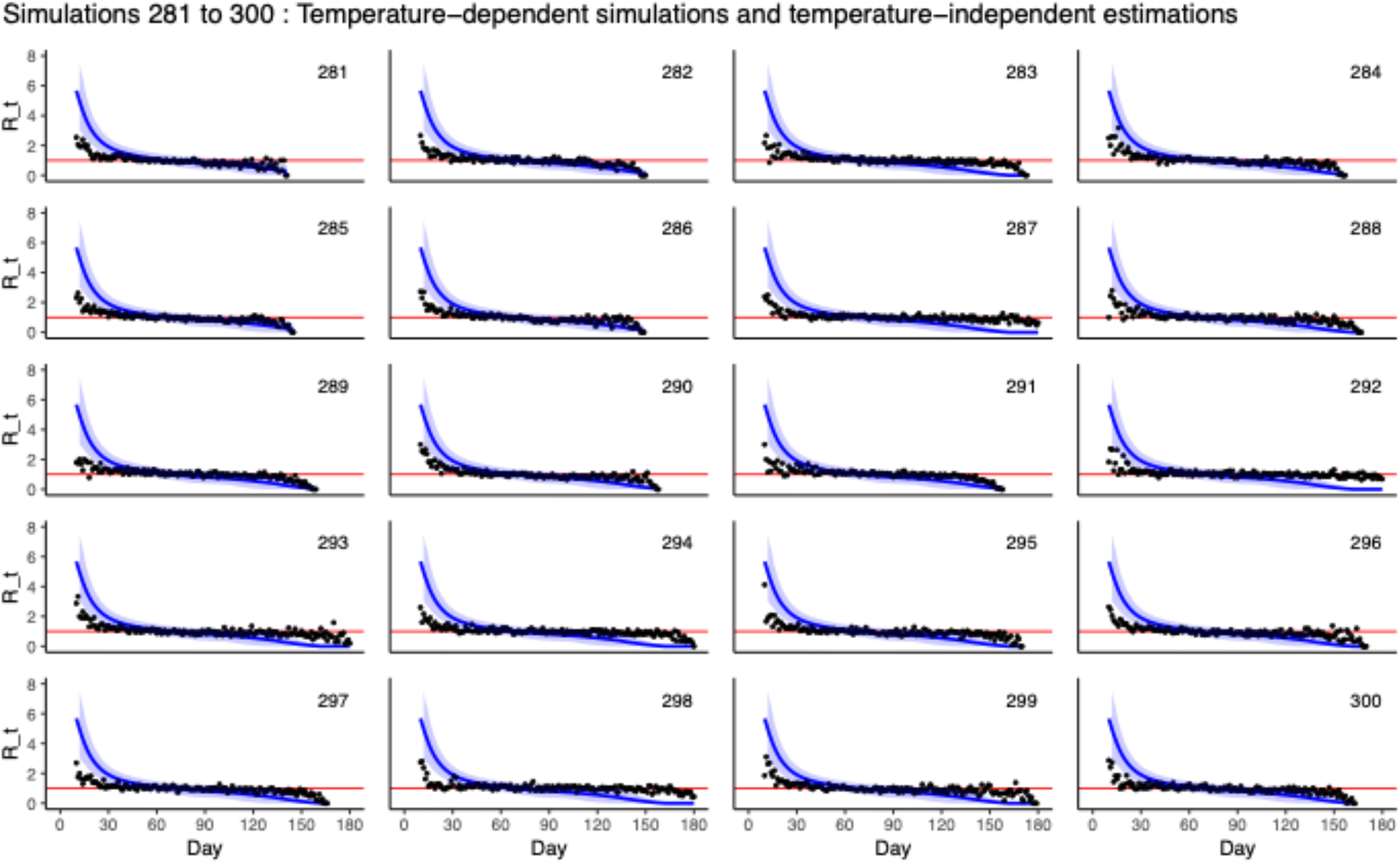
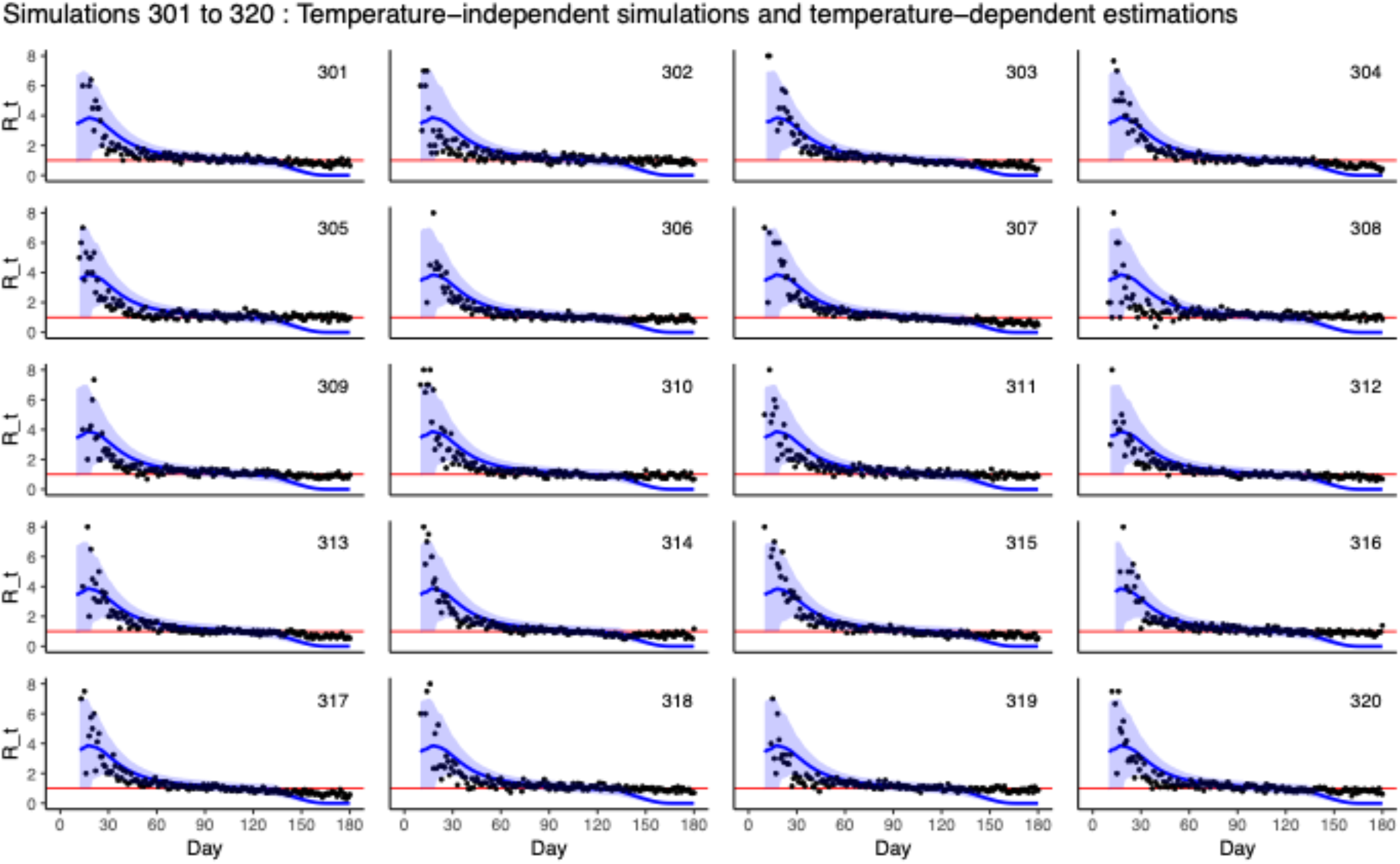
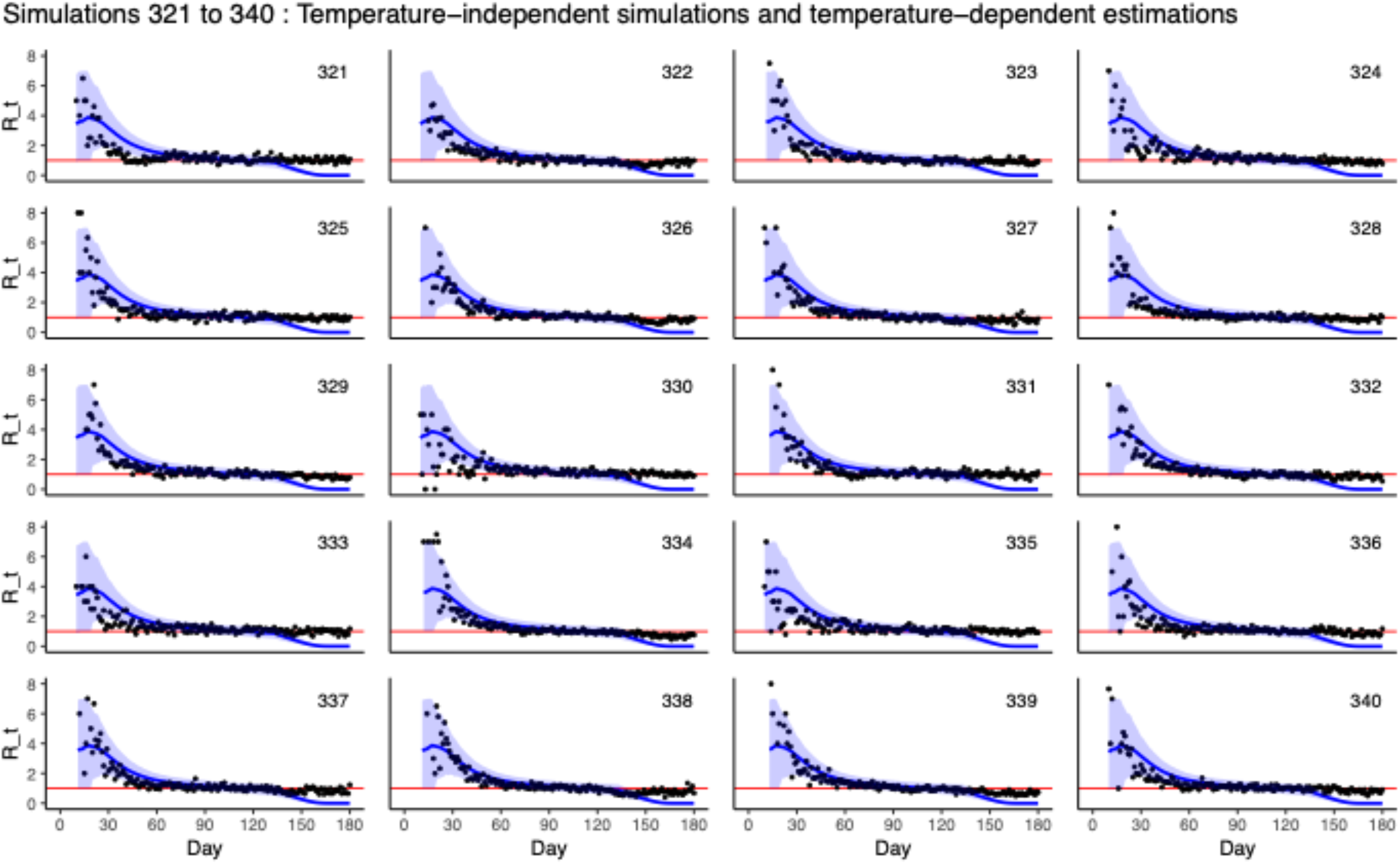
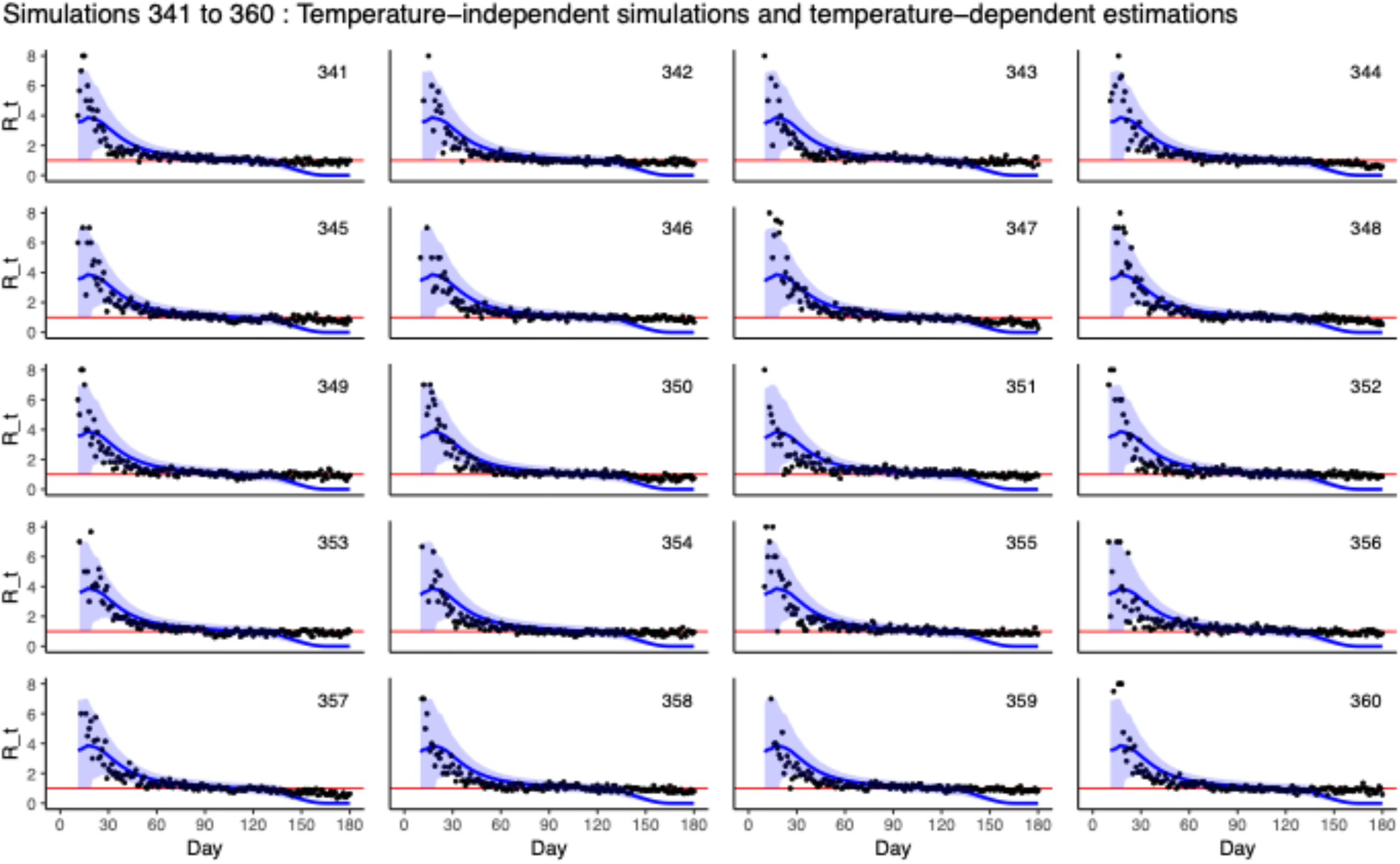
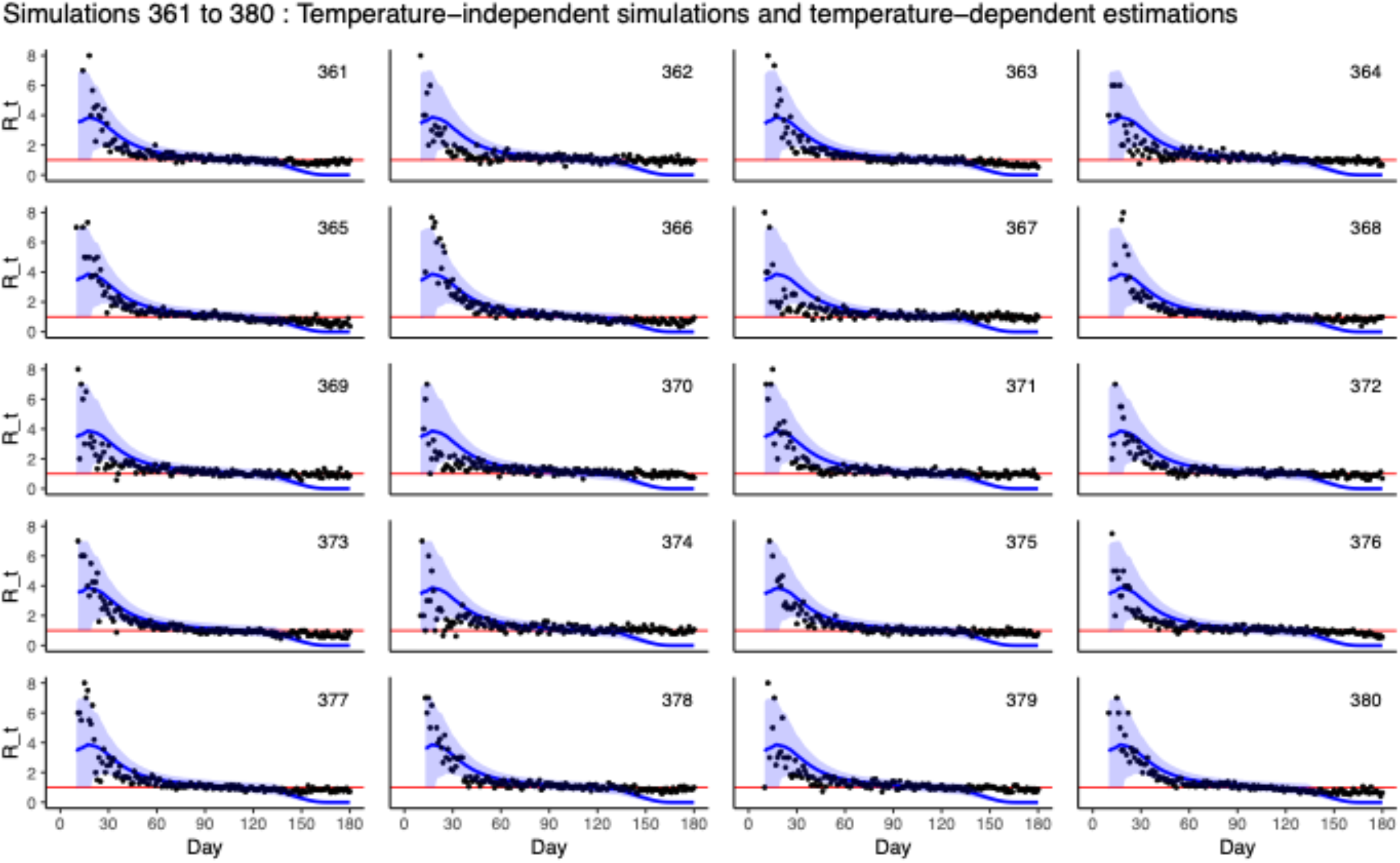
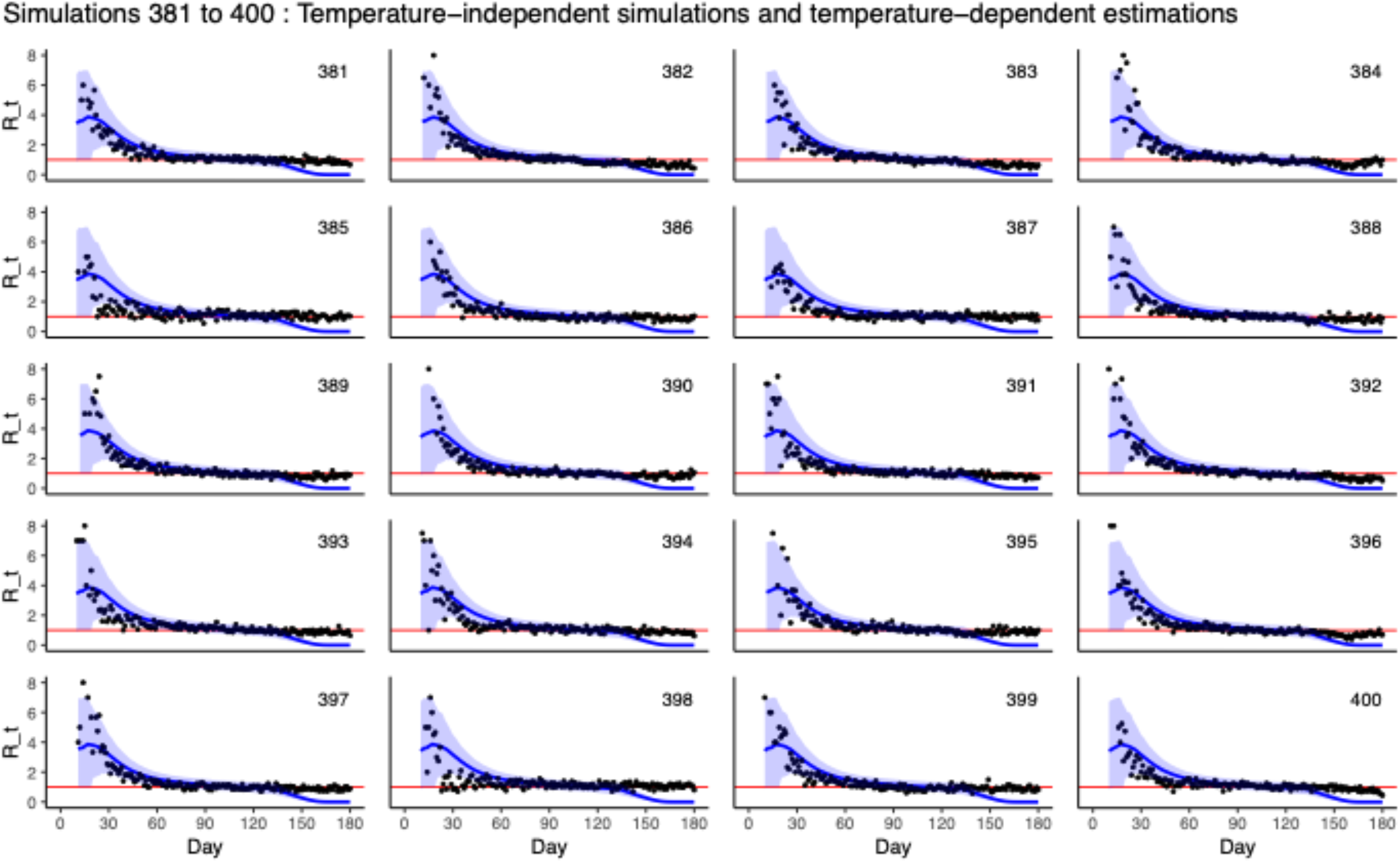
Daily true and predicted *R*_*t*_ values across individual simulations using a Gaussian decay spatial weighting function. For legibility, 20 simulations are depicted per page. The horizontal red line indicates *R*_*t*_ = 1 and the black points indicate true *R*_*t*_ values. The solid blue line depicts the mean *R*_*t*_ value from 100 bootstrap samples, whereas the shaded blue regions depict the 95% confidence intervals based on 100 bootstrap samples. A Gaussian decay spatial weighting function was used for these 400 simulations.

**Figure S3.**
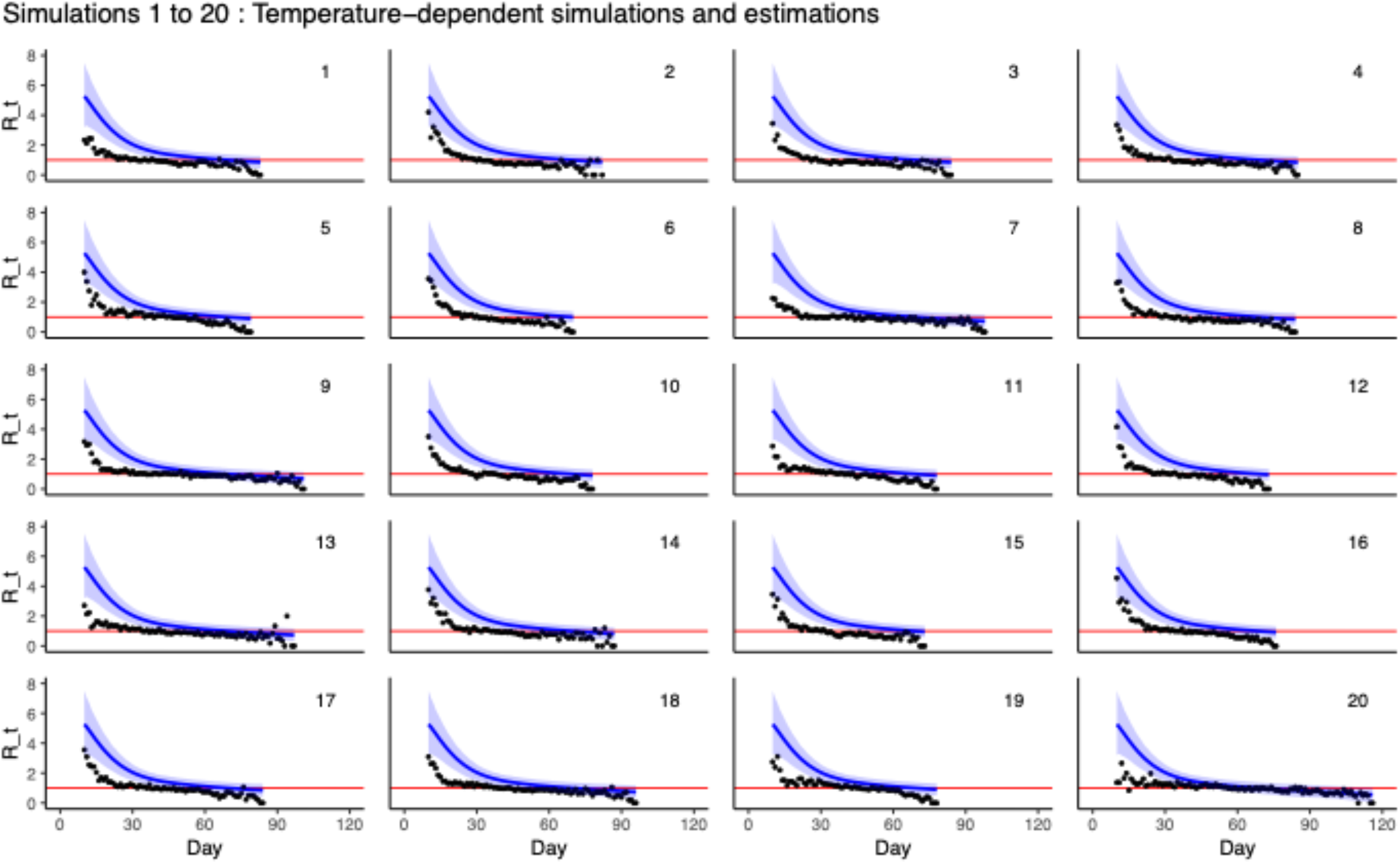
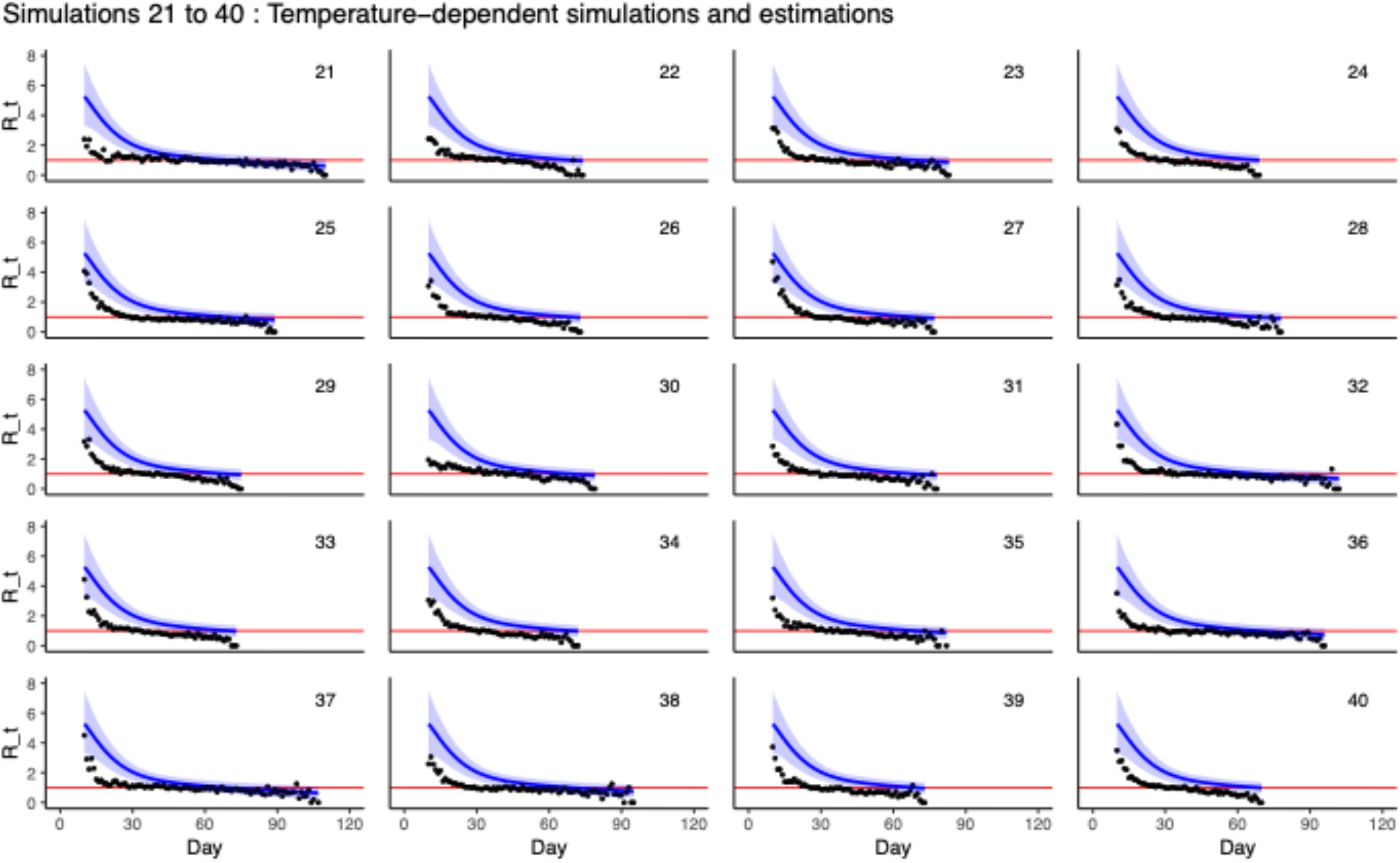
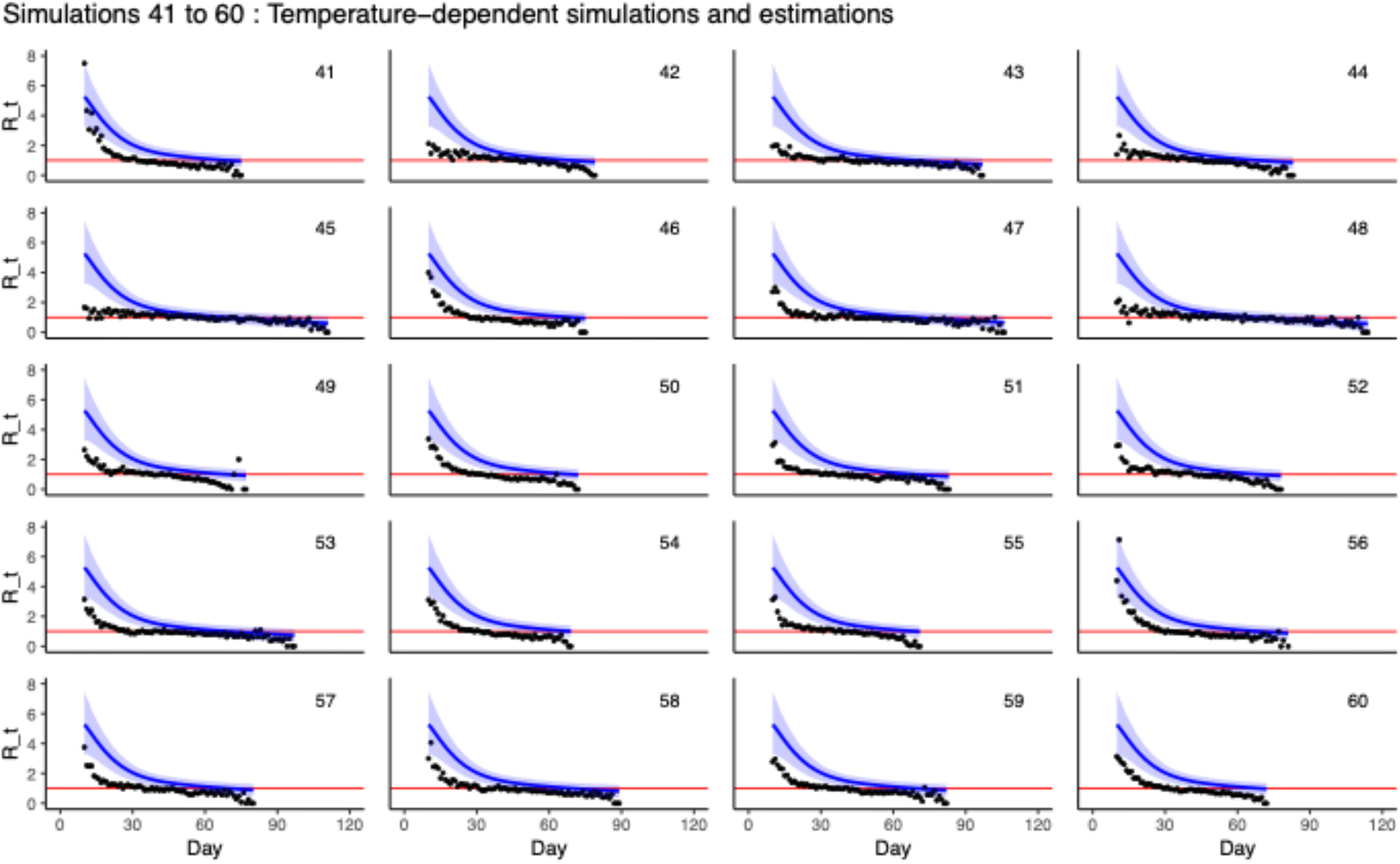
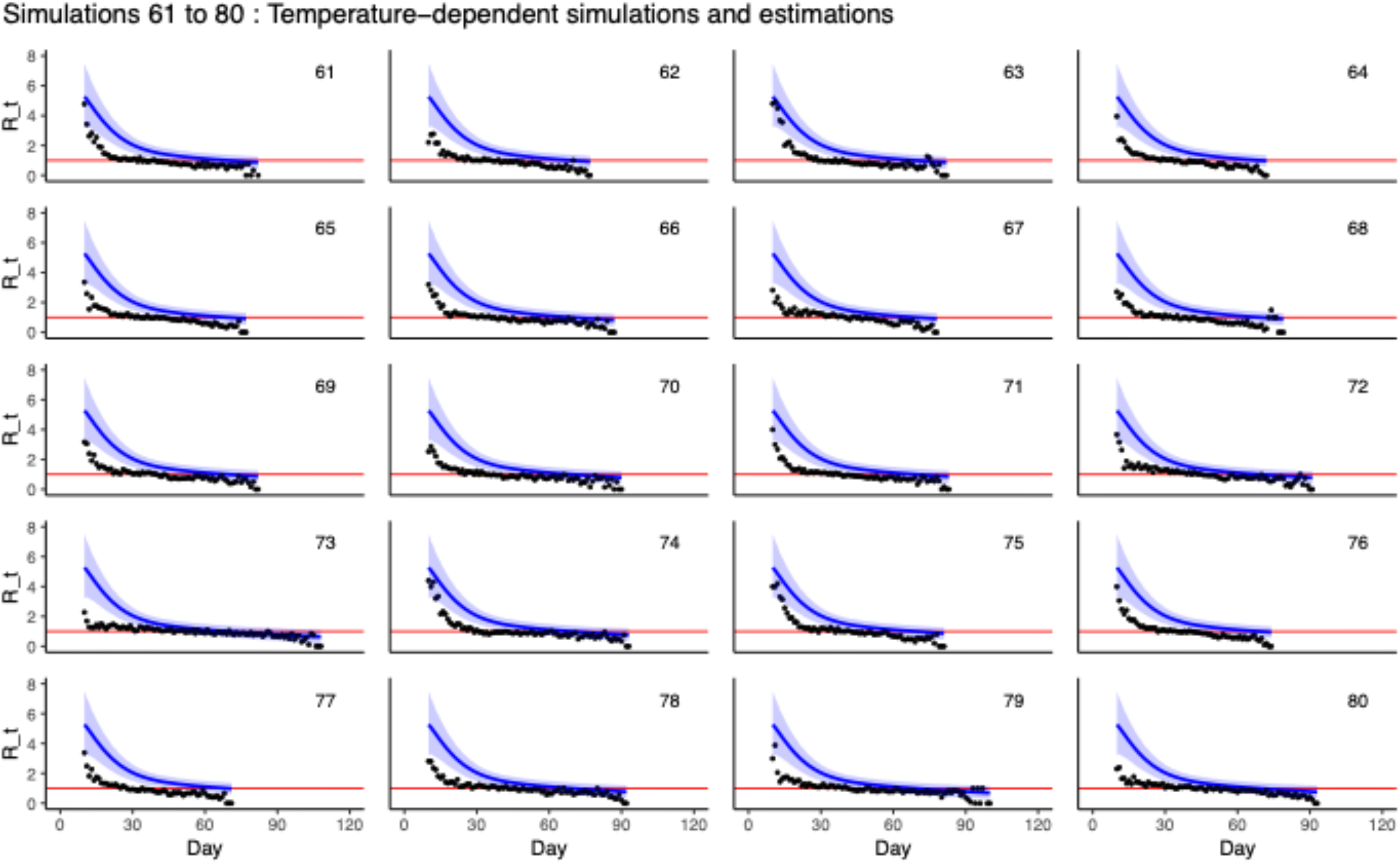
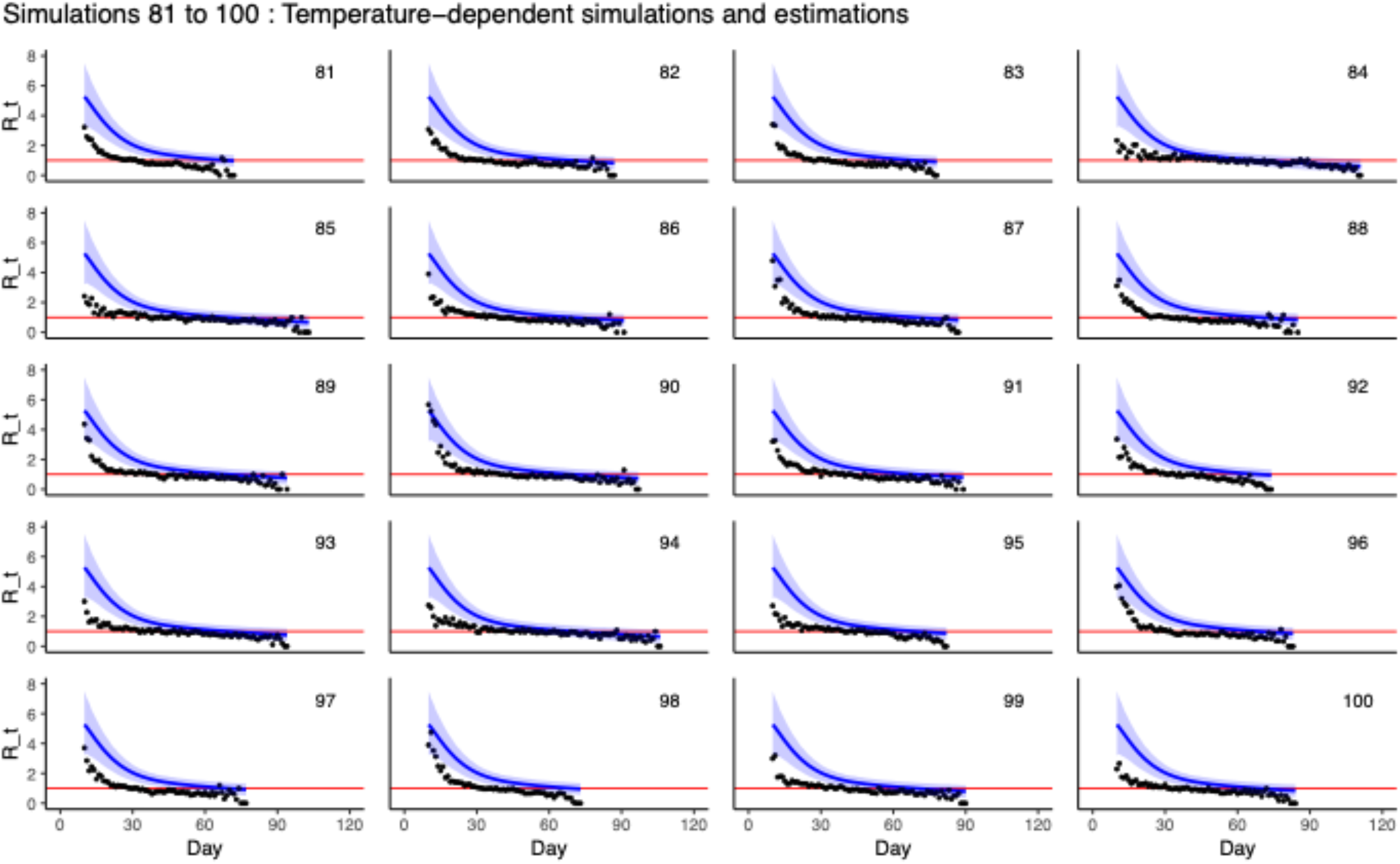
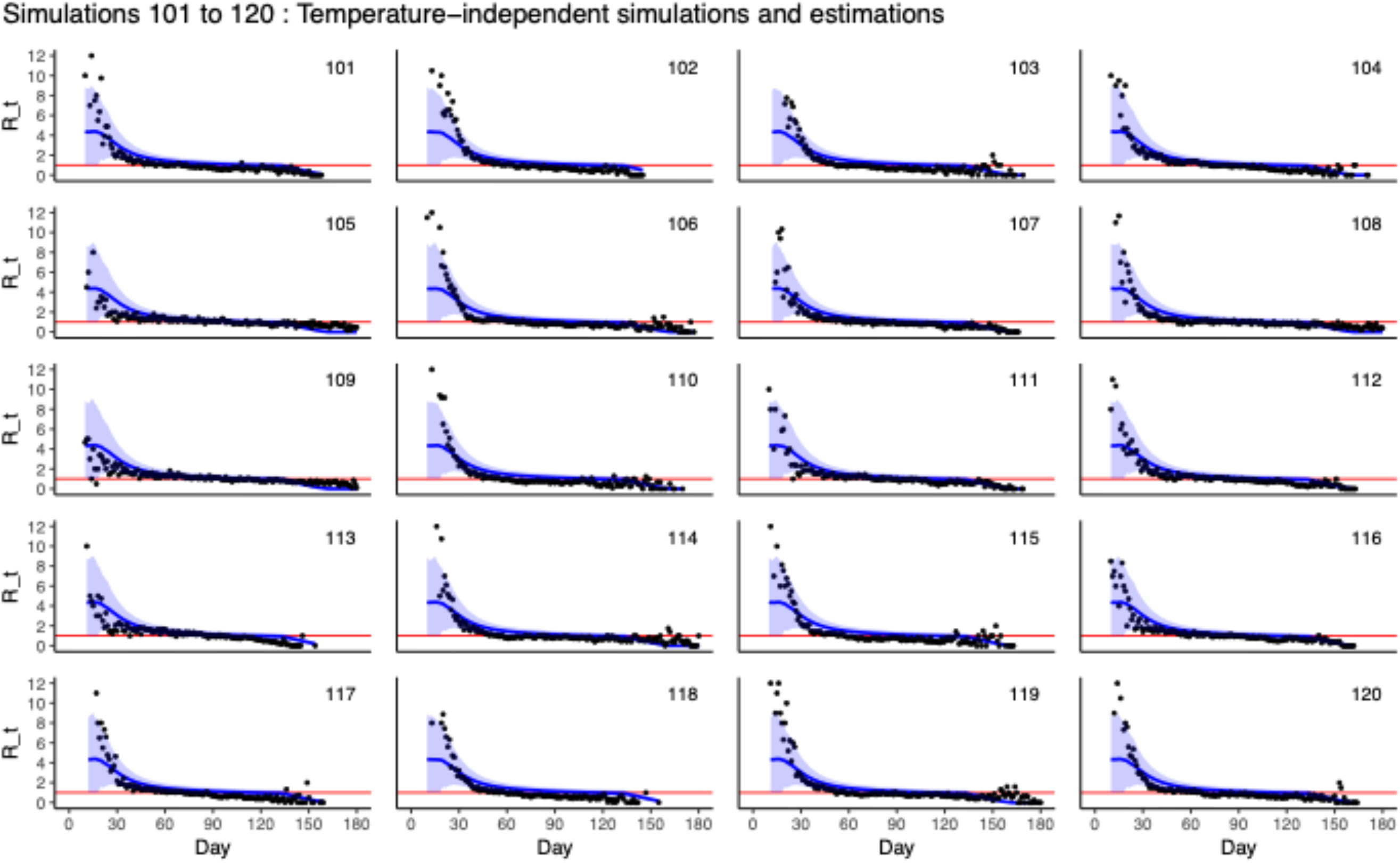
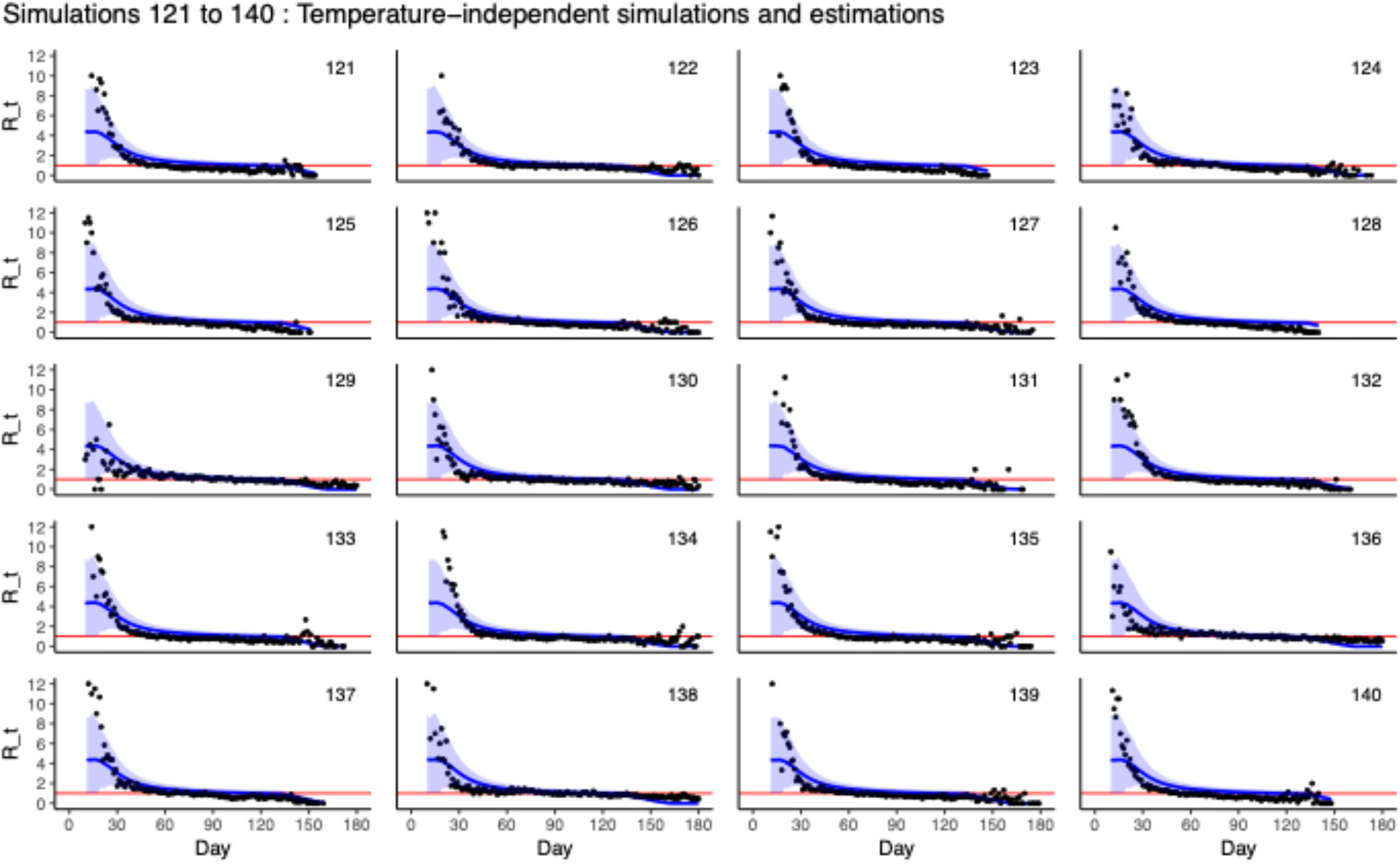
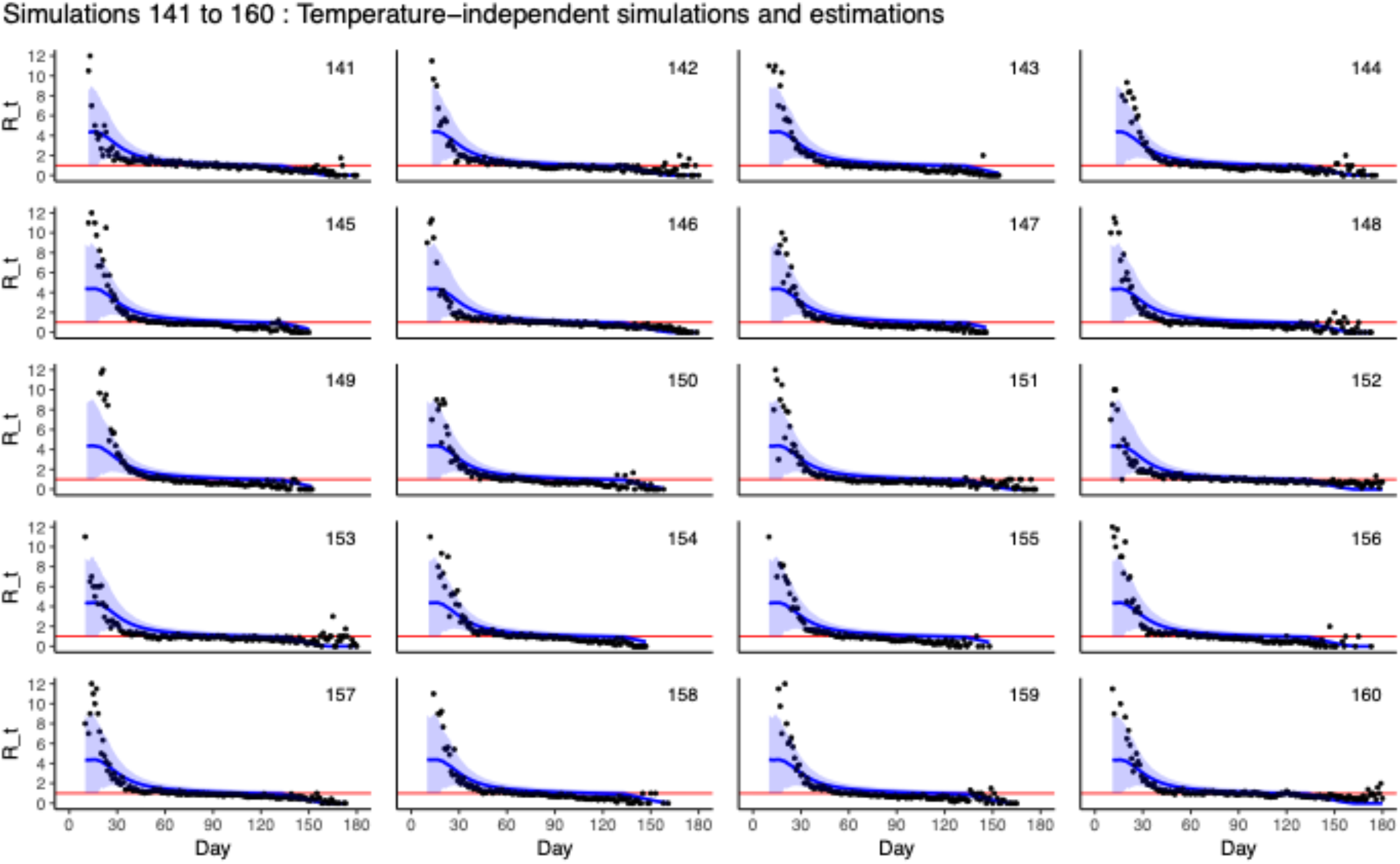
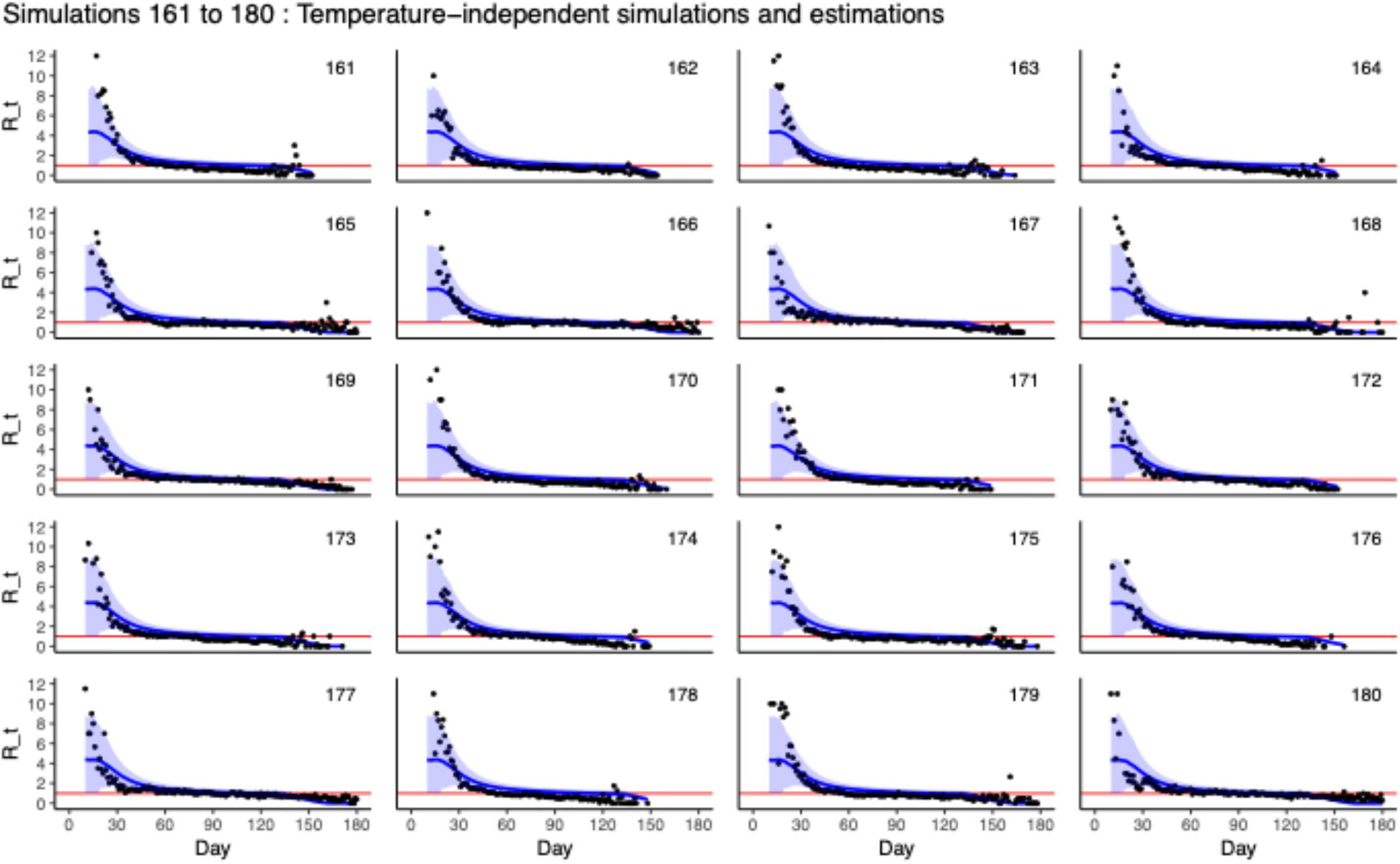
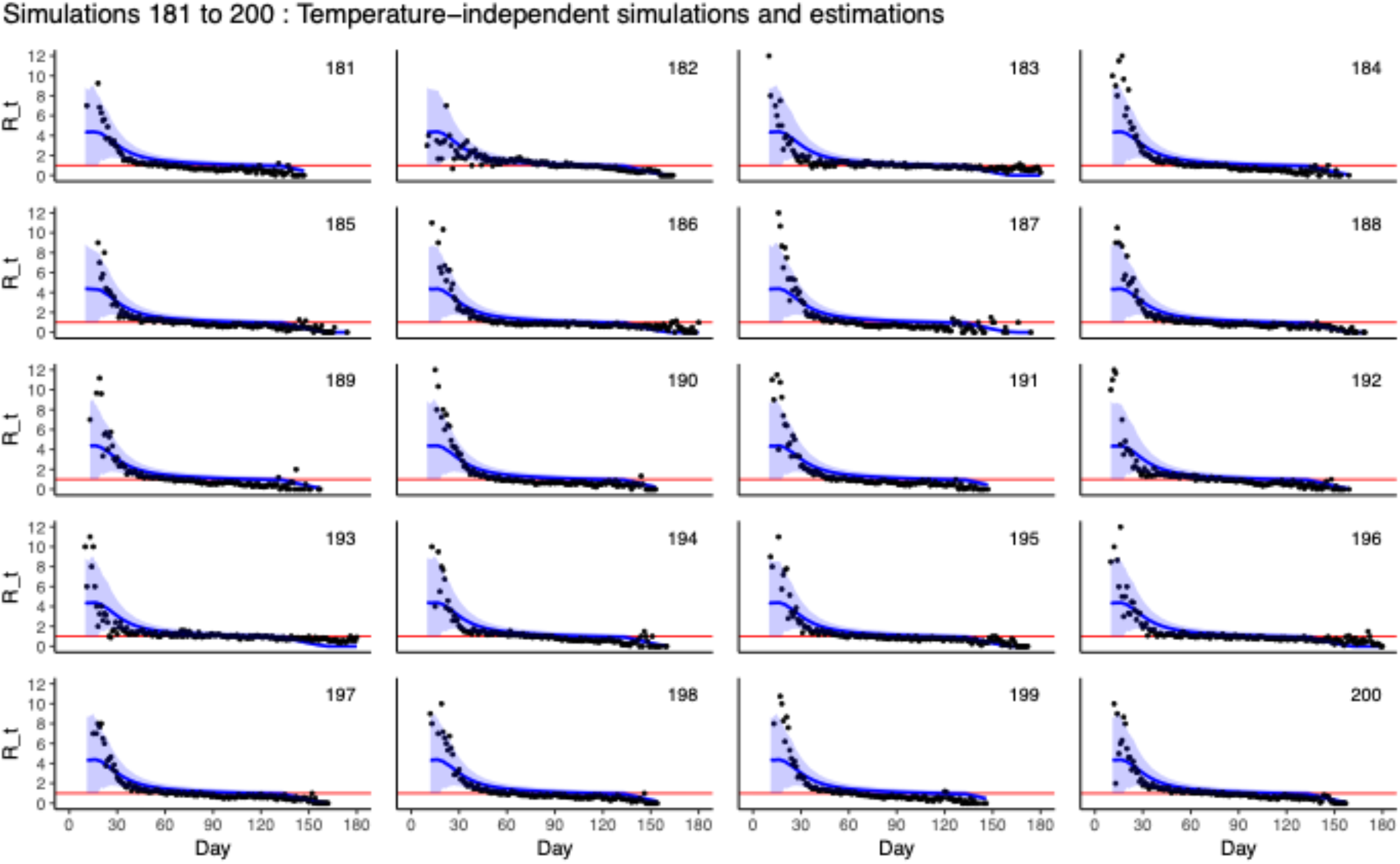
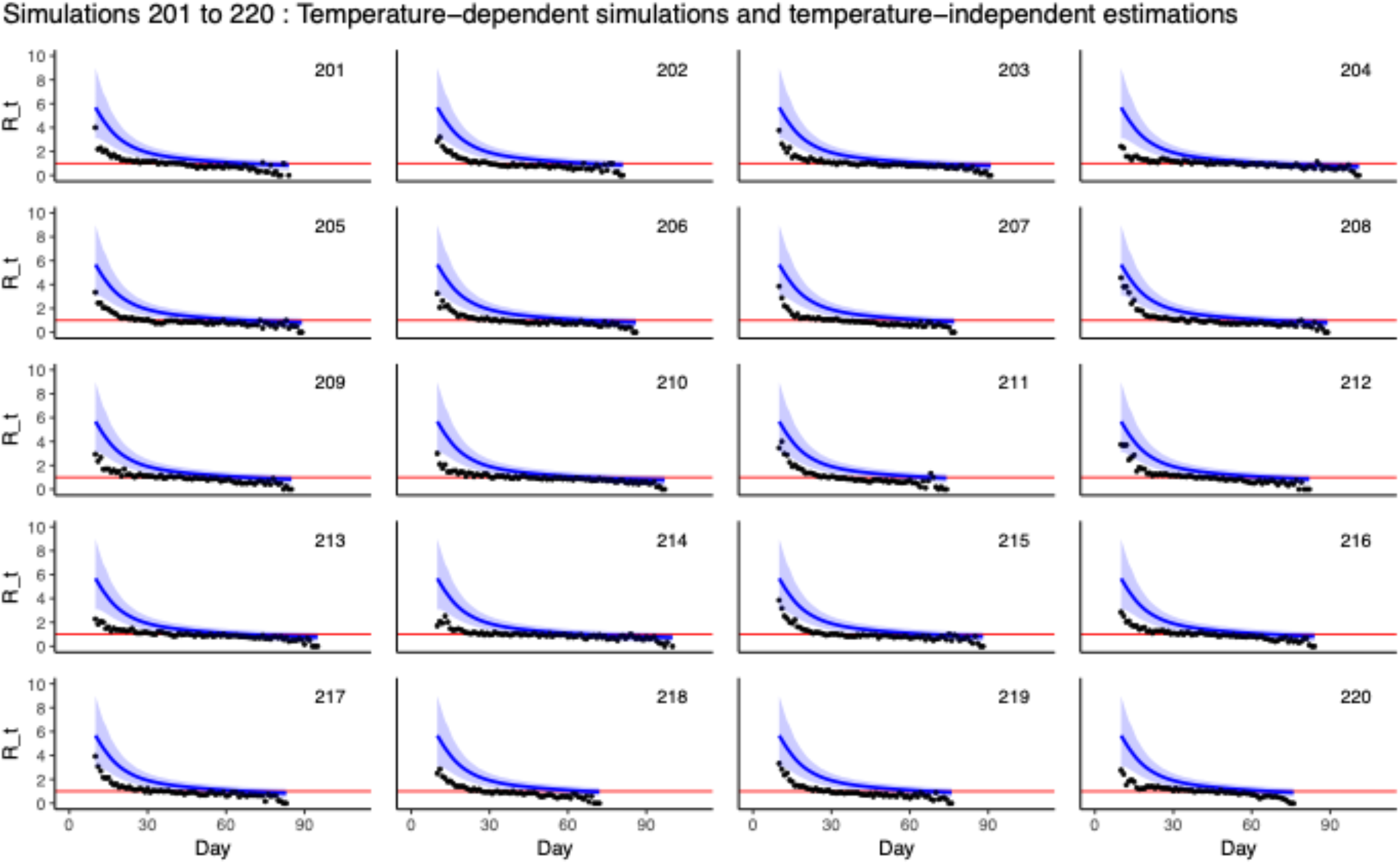
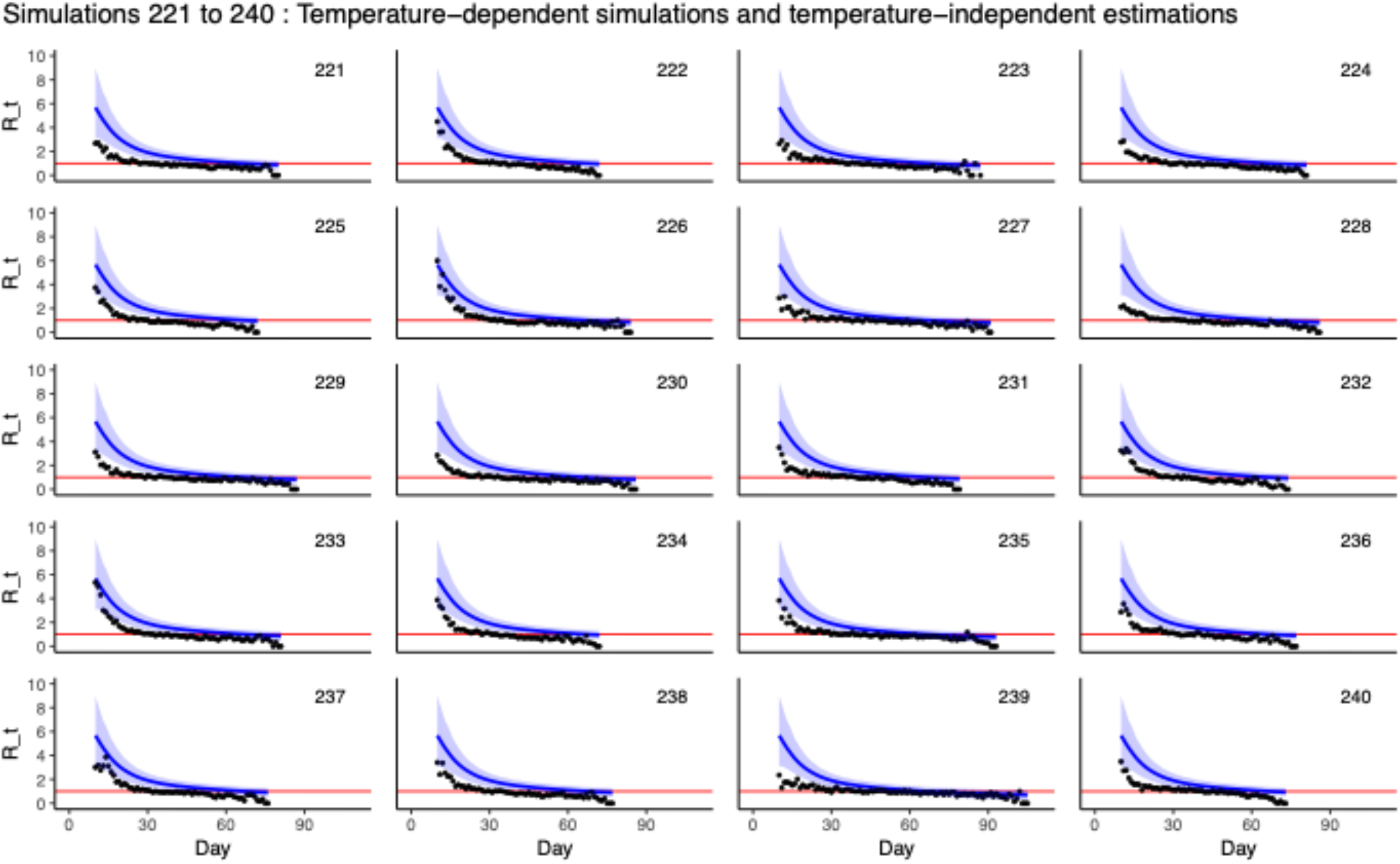
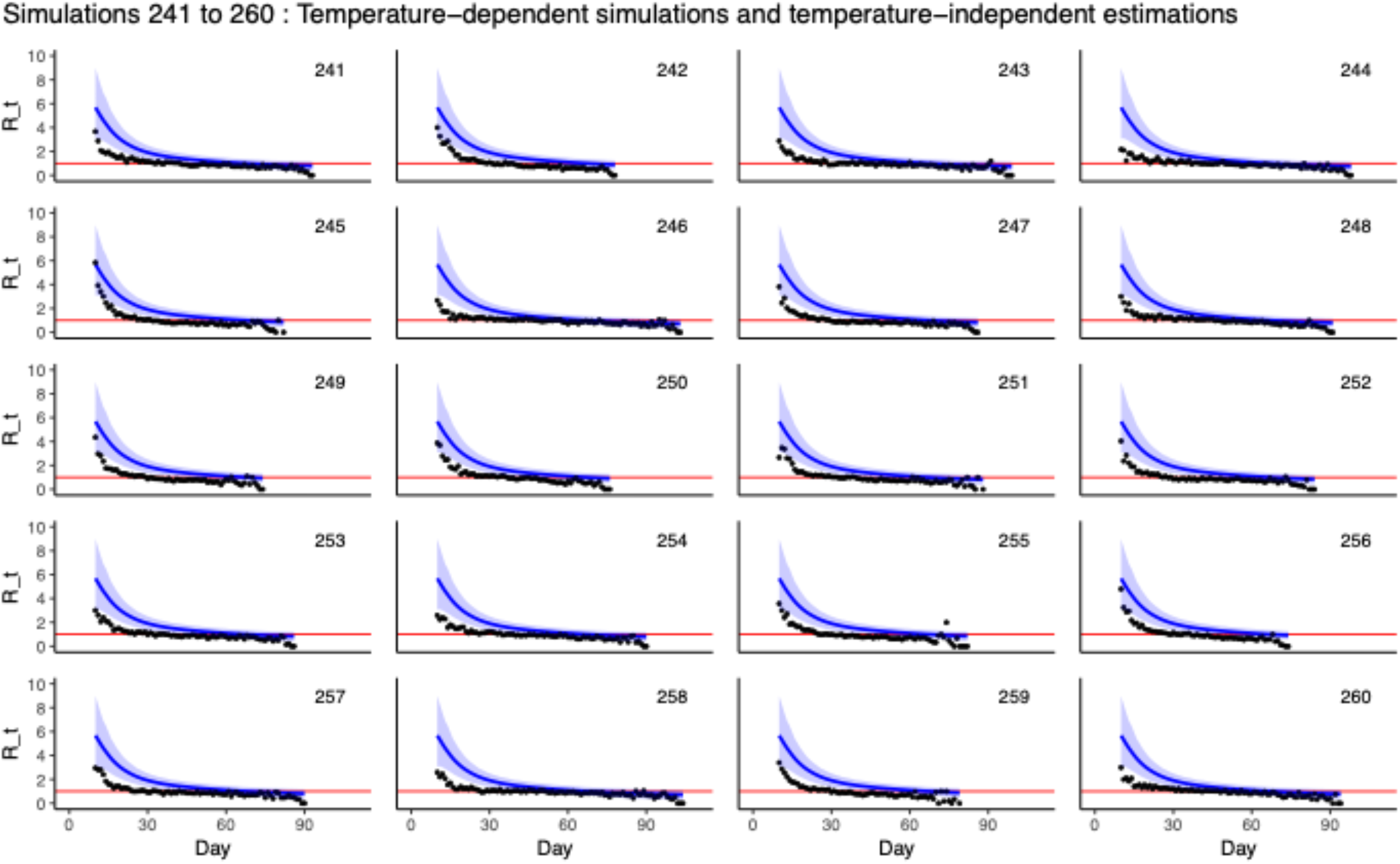
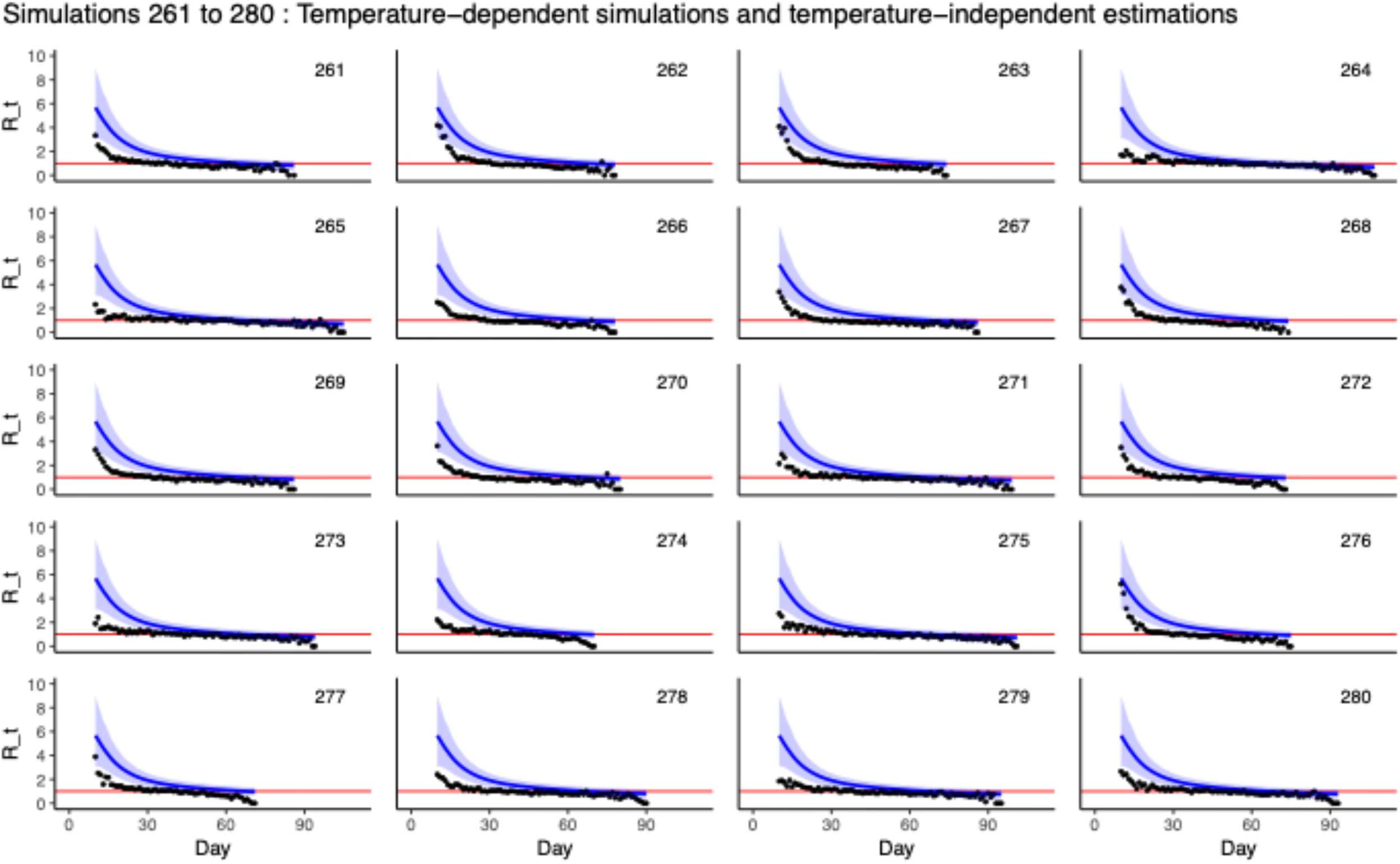
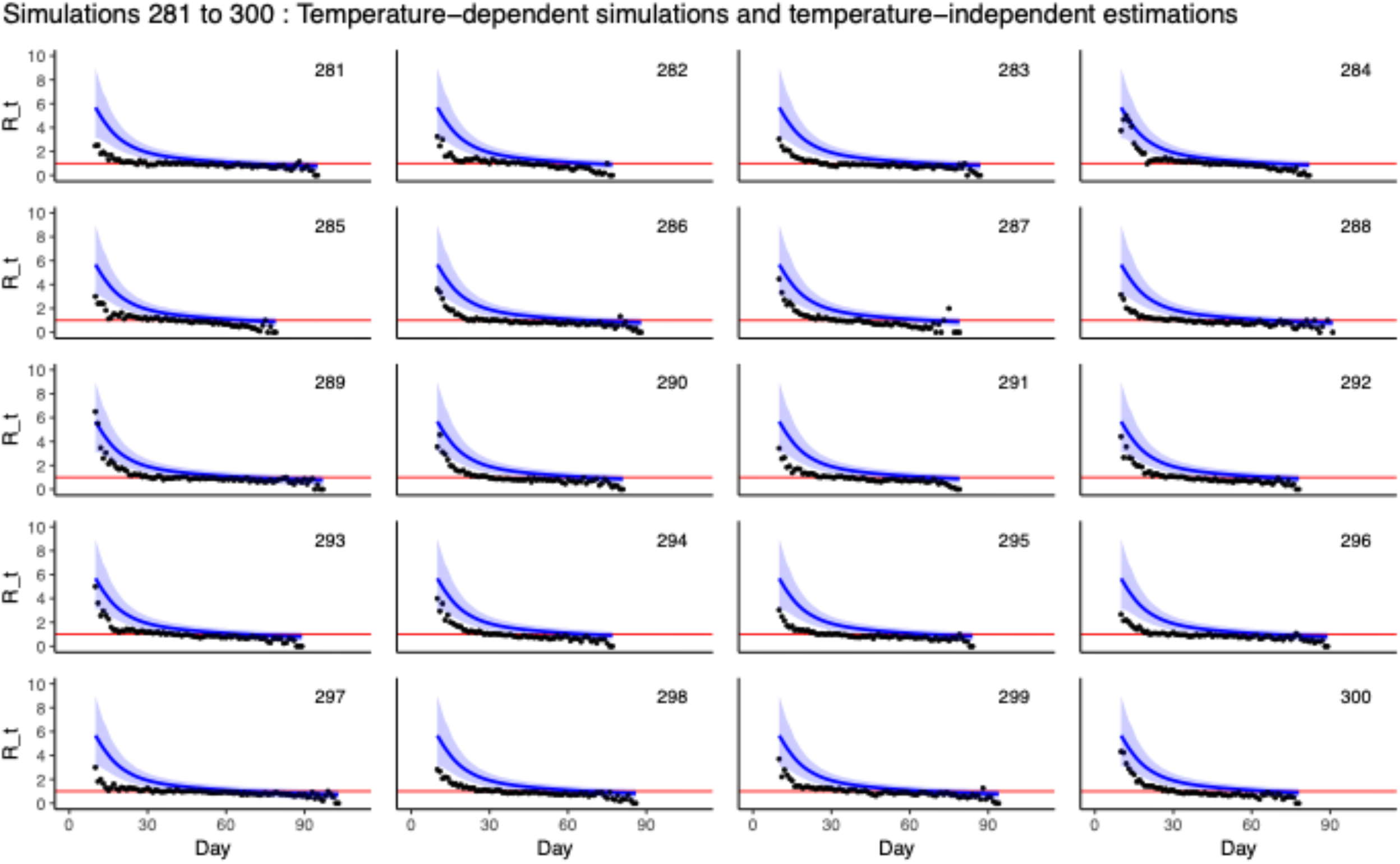
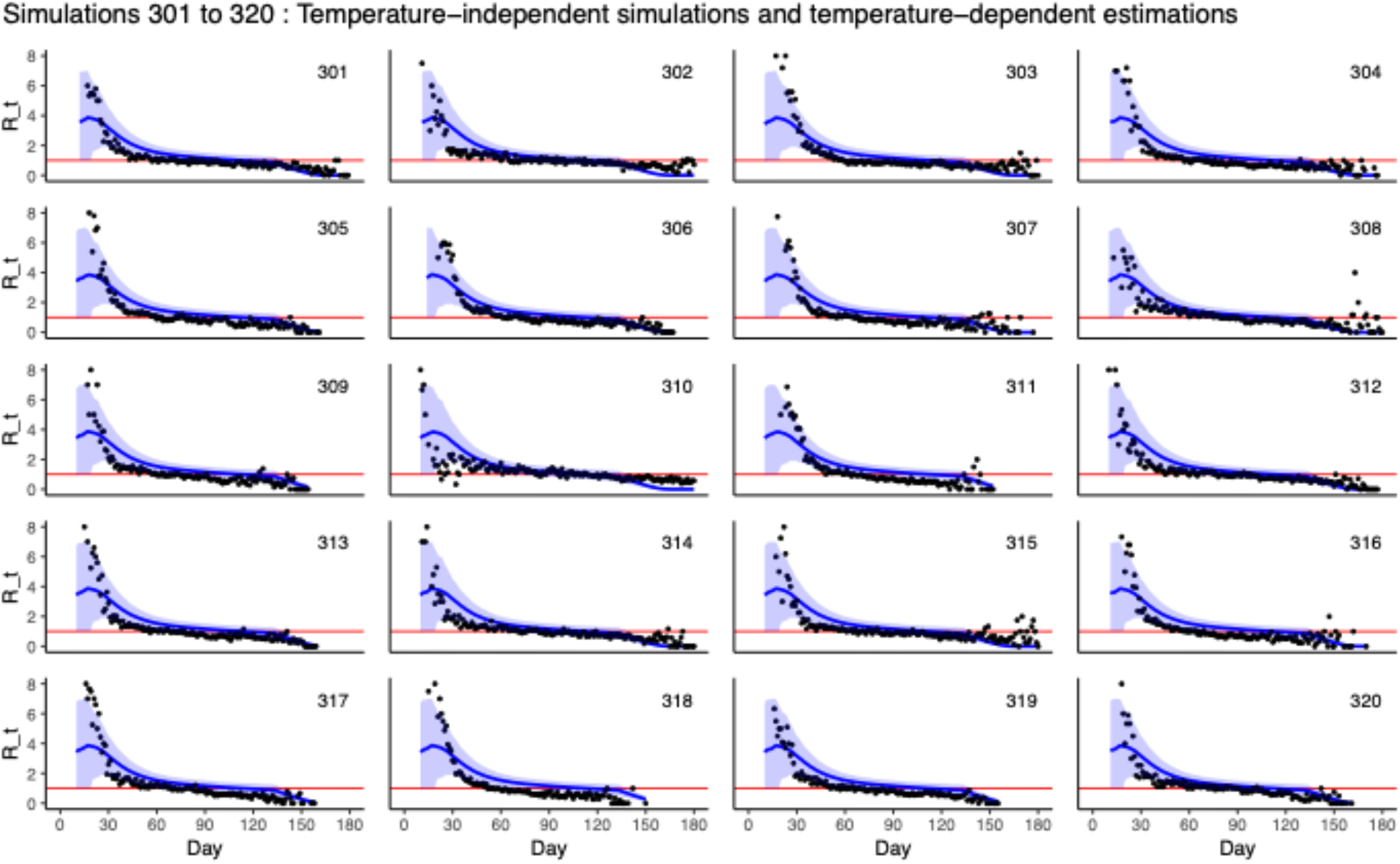
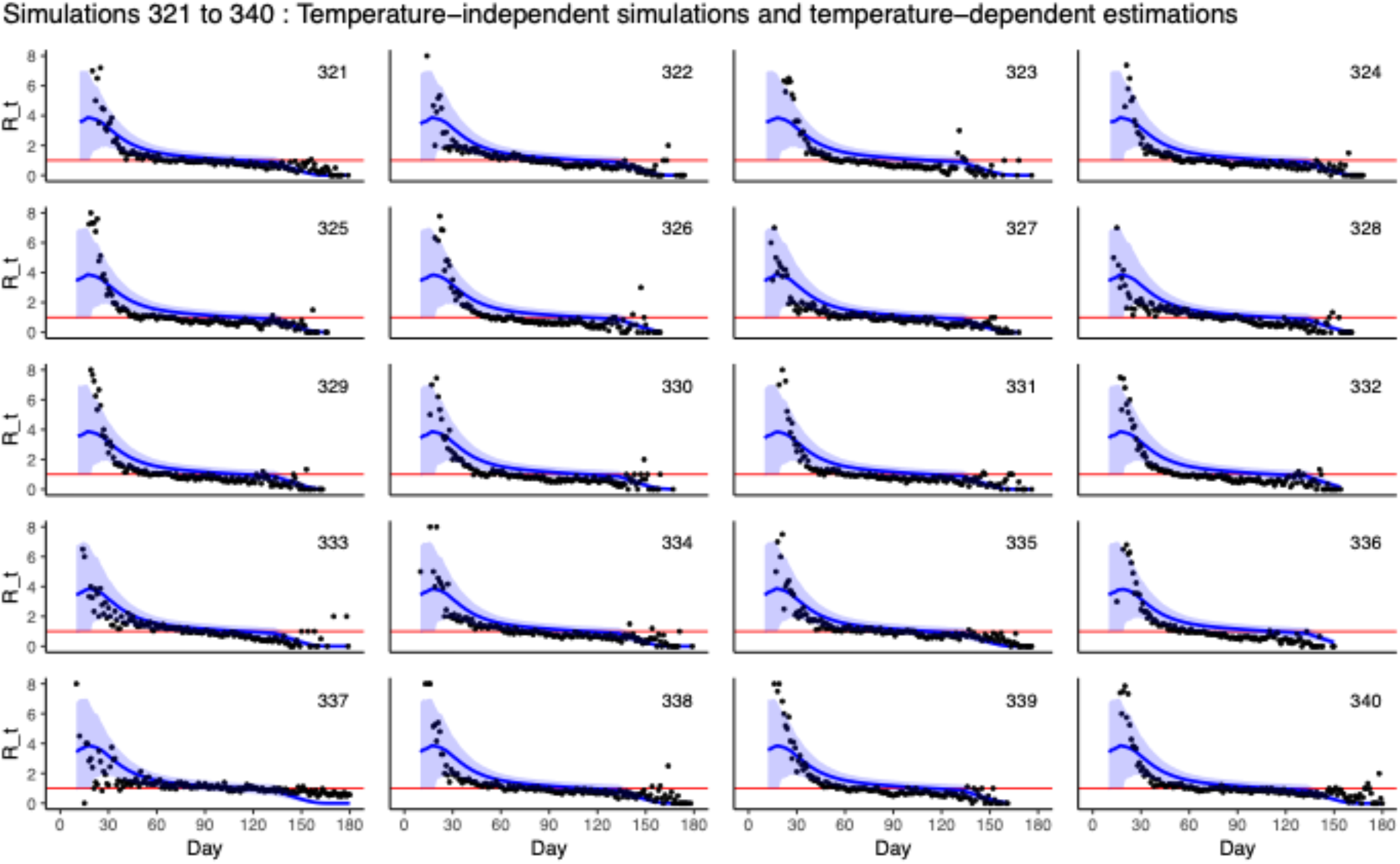
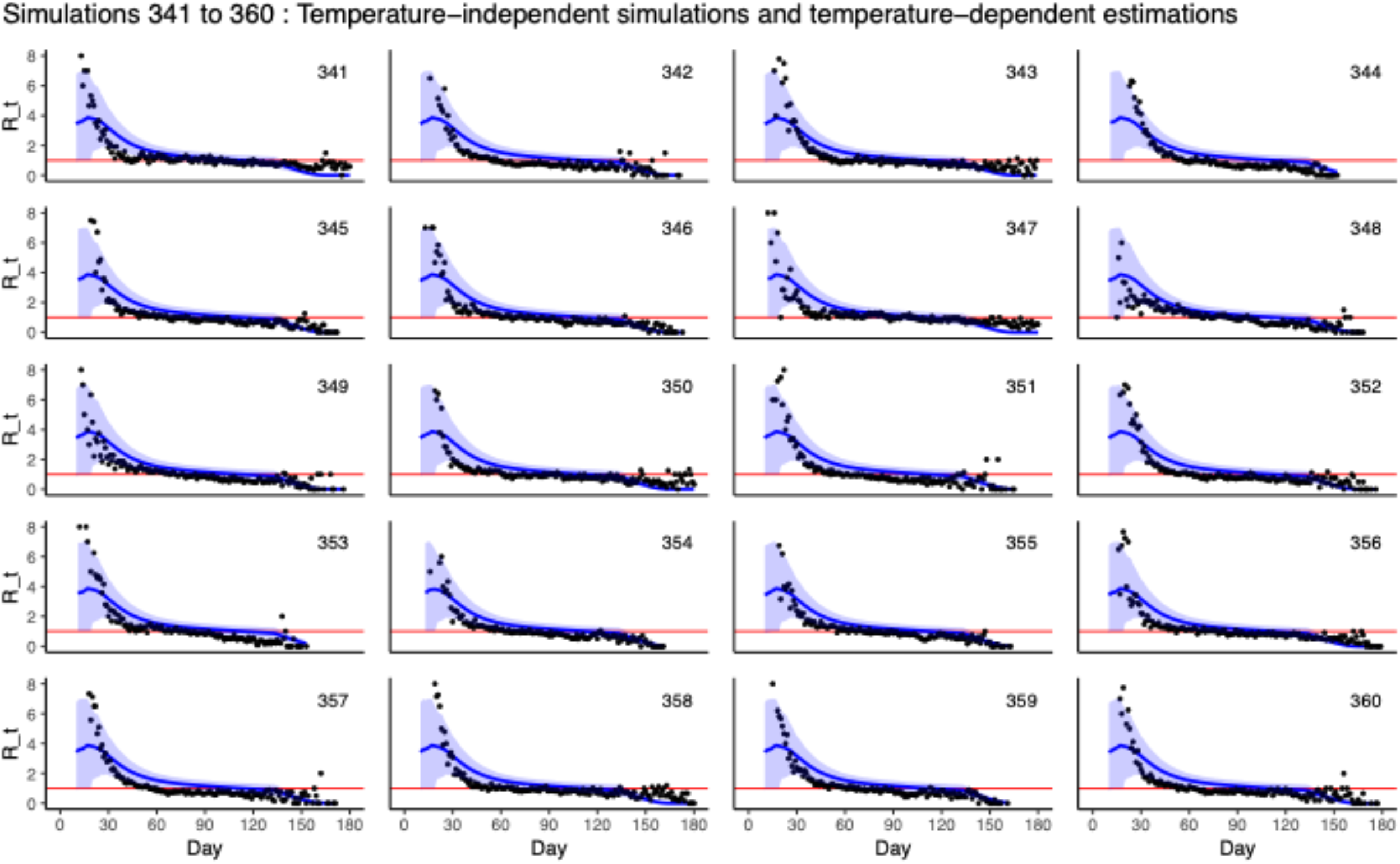
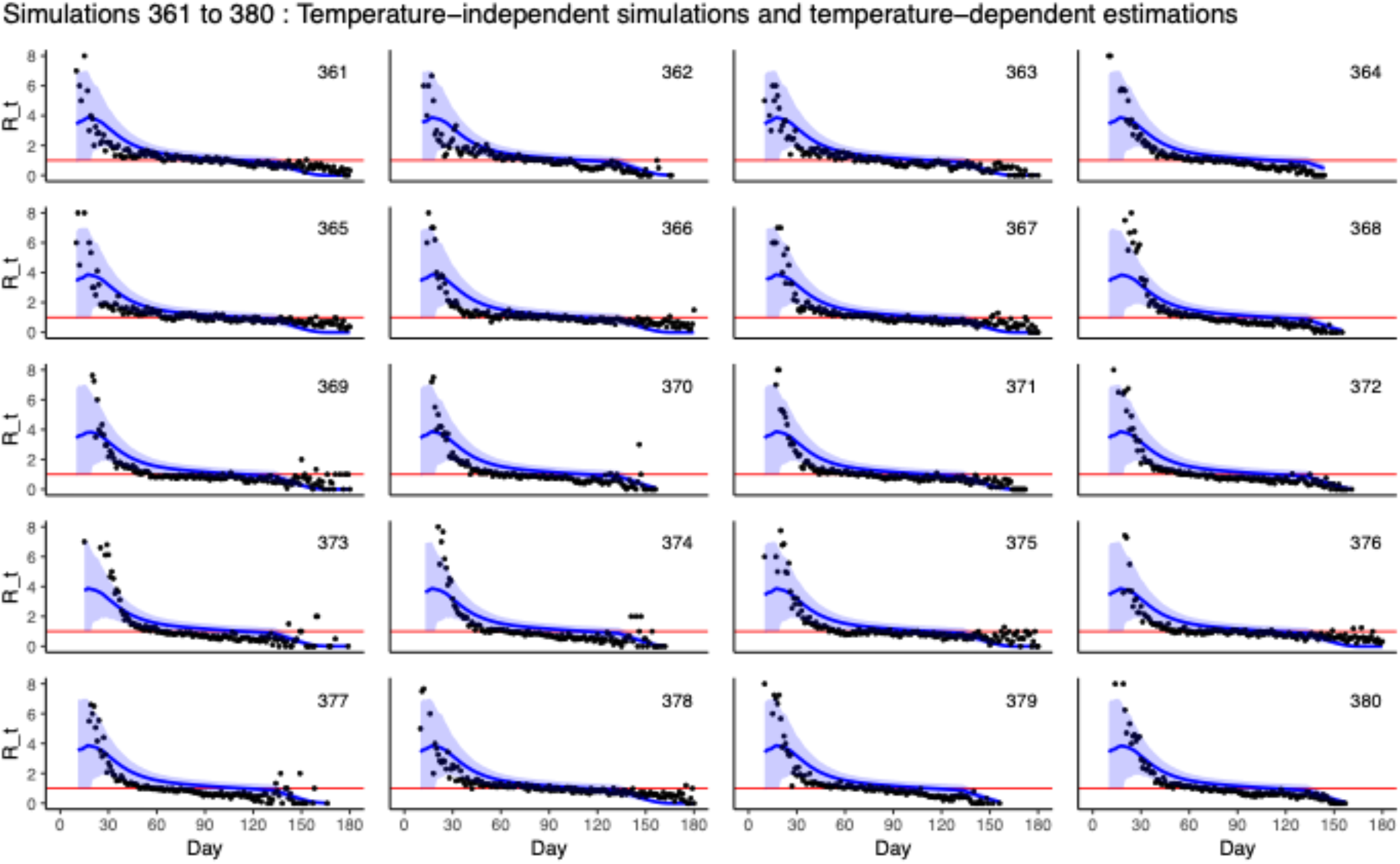
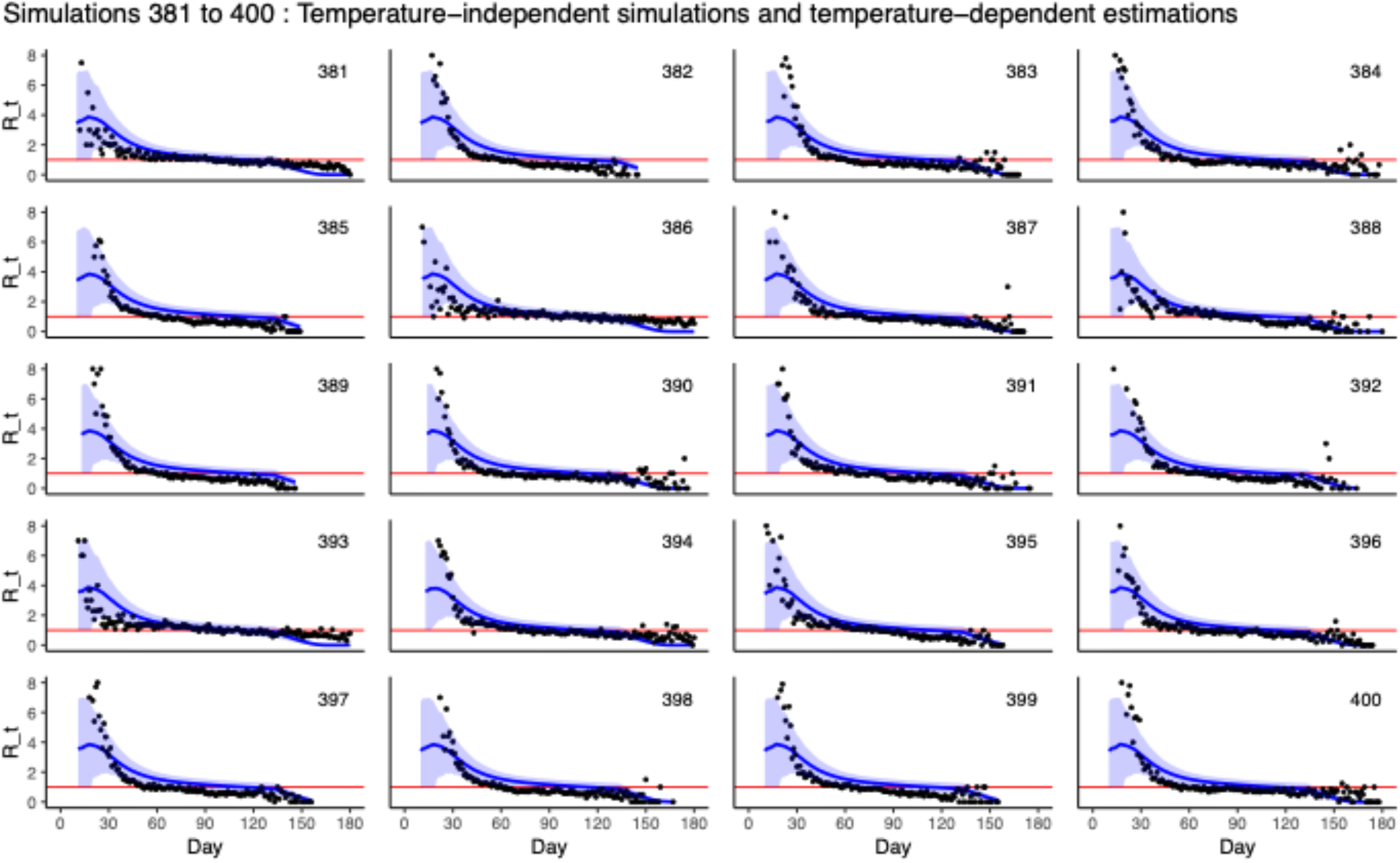
Daily true and predicted *R*_*t*_ values across individual simulations using an exponential decay spatial weighting function. For legibility, 20 simulations are depicted per page. The horizontal red line indicates *R*_*t*_ = 1 and the black points indicate true *R*_*t*_ values. The solid blue line depicts the mean *R*_*t*_ value from 100 bootstrap samples, whereas the shaded blue regions depict the 95% confidence intervals based on 100 bootstrap samples. An exponential decay spatial weighting function was used for these 400 simulations.

**Figure S4.**
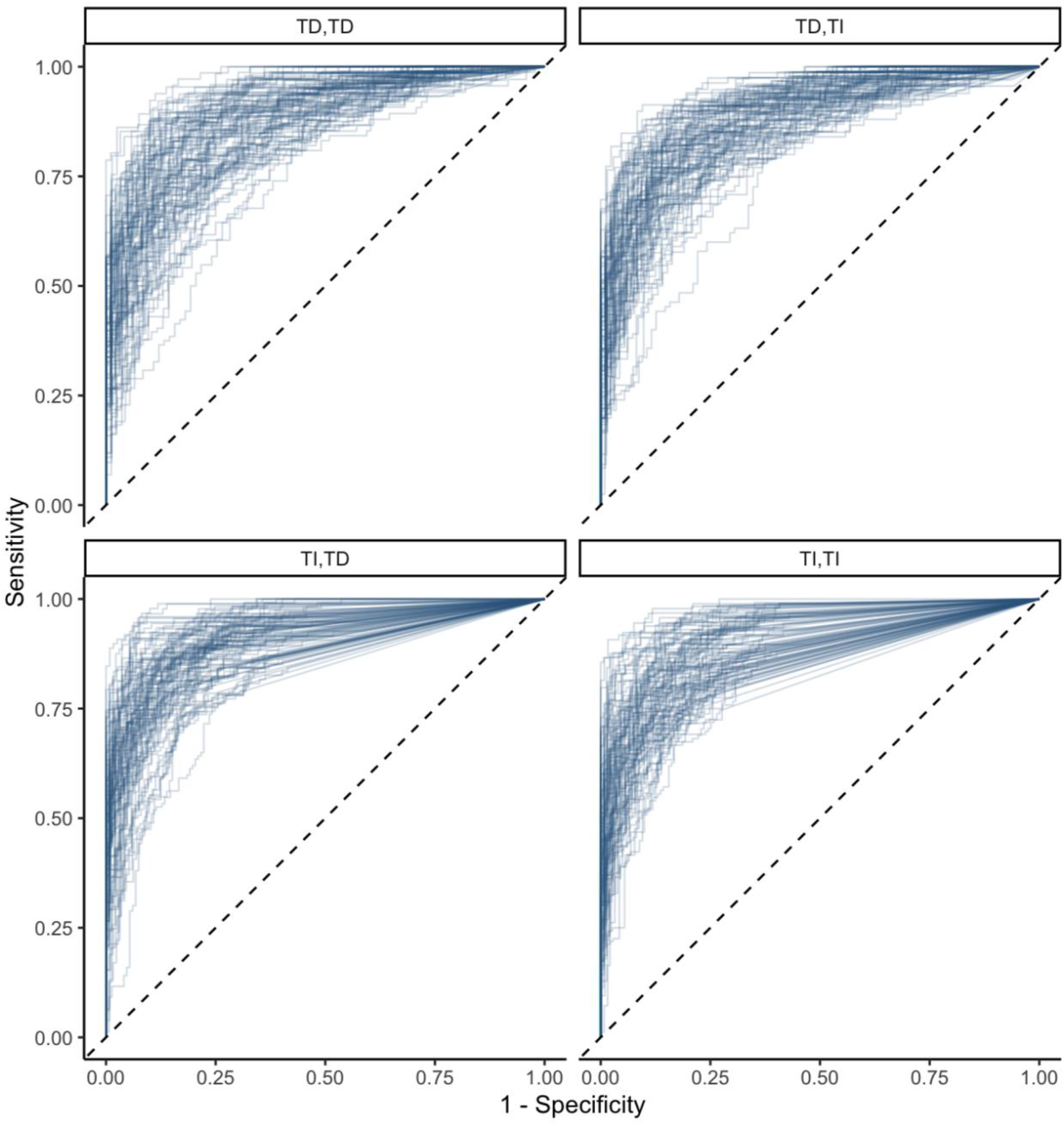
Gaussian decay spatial models’ receiver operating characteristic (ROC) curves for each combination of temperature dependence assumptions as the ground truth and in our estimators, across 100 simulations each. TD,TD: Temperature-dependent generation times as ground truth assumption and temperature-dependent generation times in estimator. TD,TI: Temperature-dependent generation times as ground truth assumption and temperature-independent generation times in estimator. TI,TD: Temperature-independent generation times as ground truth assumption and temperature-dependent generation times in estimator. TI,TI: Temperature-independent generation times as ground truth assumption and temperature-independent generation times in estimator.

**Figure S5.**
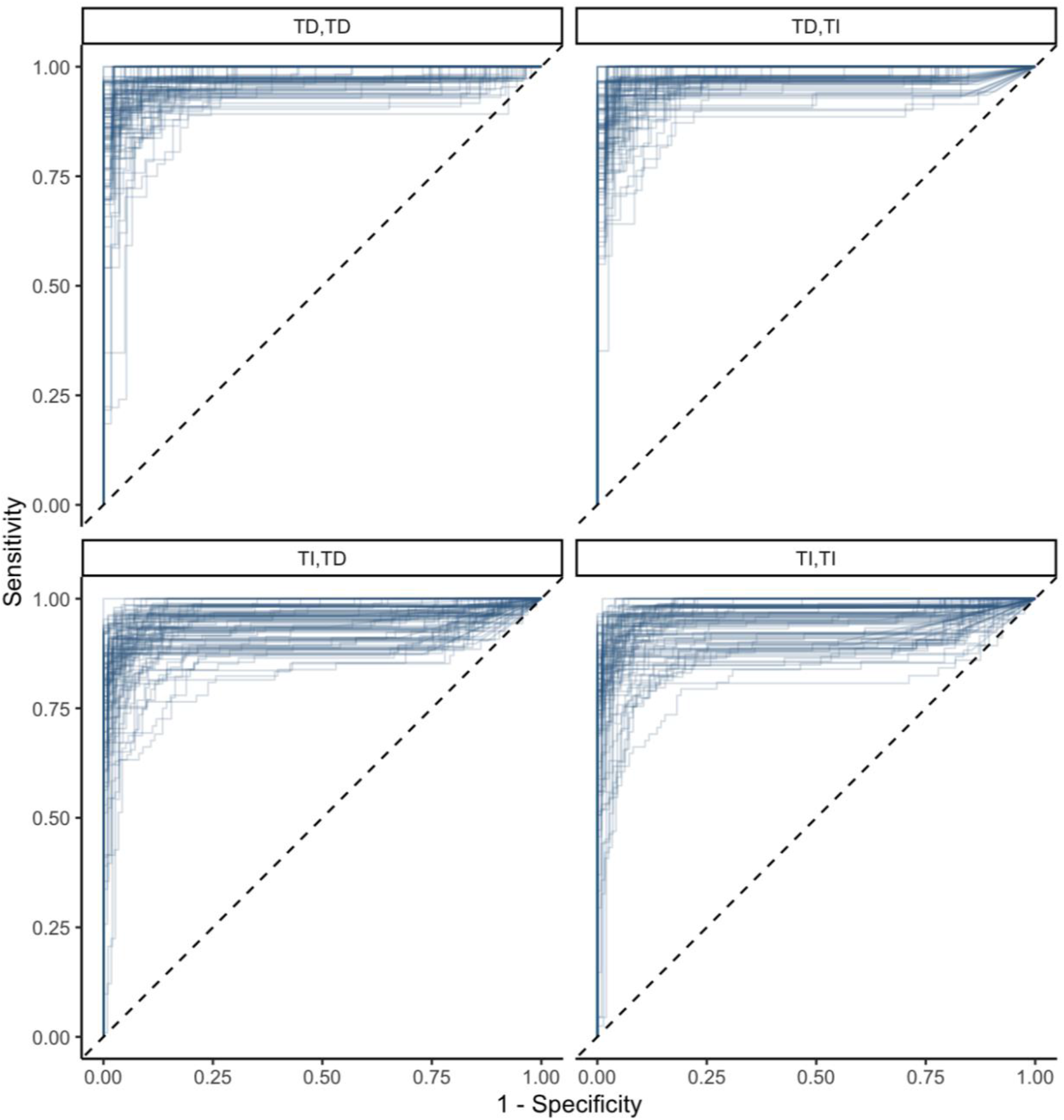
Exponential decay spatial models’ receiver operating characteristic (ROC) curves for each combination of temperature dependence assumptions as the ground truth and in our estimators, across 100 simulations each. TD,TD: Temperature-dependent generation times as ground truth assumption and temperature-dependent generation times in estimator. TD,TI: Temperature-dependent generation times as ground truth assumption and temperature-independent generation times in estimator. TI,TD: Temperature-independent generation times as ground truth assumption and temperature-dependent generation times in estimator. TI,TI: Temperature-independent generation times as ground truth assumption and temperature-independent generation times in estimator.

